# Intratumoral gene delivery of 4-1BBL boosts IL-12-triggered anti-glioblastoma immunity

**DOI:** 10.1101/2025.02.03.636330

**Authors:** Taral R. Lunavat, Lisa Nieland, Sanne M. van de Looij, Alexandra J.E.M. de Reus, Charles P. Couturier, Chadi A. El Farran, Tyler E. Miller, Julia K. Lill, Maryam Schübel, Tianhe Xiao, Emilio Di Ianni, Elliot C. Woods, Yi Sun, David Rufino-Ramos, Thomas S. van Solinge, Shadi Mahjoum, Emily Grandell, Mao Li, Vamsi Mangena, Gavin P. Dunn, Russell W. Jenkins, Thorsten R. Mempel, Xandra O. Breakefield, Koen Breyne

**Author notes:** **Corresponding author**: Koen Breyne, Ph.D., Molecular Neurogenetics Unit, Massachusetts General Hospital, 13^th^ Street, Building 149, Charlestown, MA 02129 USA. These authors contributed equally (Taral R. Lunavat, Lisa Nieland).

## Abstract

The standard of care in high-grade gliomas has remained unchanged in the past 20 years. Efforts to replicate effective immunotherapies in non-cranial tumors have led to only modest therapeutical improvements in glioblastoma (GB).

Here, we demonstrate that intratumoral administration of recombinant interleukin-12 (rIL-12) promotes local cytotoxic CD8^POS^ T cell accumulation and conversion into an effector-like state, resulting in a dose-dependent survival benefit in preclinical GB mouse models. This tumor-reactive CD8 T cell response is further supported by intratumoral rIL-12-sensing dendritic cells (DCs) and is accompanied by the co-stimulatory receptor 4-1BB expression on both cell types. Given that DCs and CD8^POS^ T cells are functionally suppressed in the tumor microenvironments of *de novo* and recurrent glioma patients, we tested whether anti-tumor response at the rIL-12-inflamed tumor site could be enhanced with 4-1BBL, the ligand of 4-1BB. 4-1BBL was delivered using an adeno-associated virus (AAV) vector targeting GFAP-expressing cells and resulted in prolonged survival of rIL-12 treated GB-bearing mice.

This study establishes that tumor antigen-specific CD8 T cell activity can be directed using an AAV-vector-mediated gene therapy approach, effectively enhancing anti-GB immunity.

## INTRODUCTION

Glioblastoma isocitrate dehydrogenase (IDH)-wild type (GB) is the most lethal primary cancer in the central nervous system (CNS) (Louis et al., 2021; Molinaro et al., 2019), with a median survival of 14.7 months after first diagnosis (van Solinge et al., 2022; Zhu et al., 2017). The current standard of care treatment paradigm includes surgical tumor resection, followed by radiotherapy and temozolomide (Stupp et al., 2005). Although the complexity of GB immunology is still being uncovered, GB is generally considered a “cold” tumor typically marked by minimal expression of neoantigens and the presence of various immune checkpoints and immune-inhibitory cytokines that augment the immunosuppressive nature of this cancer (Lin et al., 2024; Segura-Collar et al., 2023). It is believed that even when an anti-tumor immune response develops, it is suppressed not only by tumor cells but also by a ‘hijacked’ tumor microenvironment (TME) (Broekman et al., 2018; Sharma et al., 2023).

Numerous clinical trials have aimed to reinvigorate anti-tumor immunity through targeting immune checkpoint inhibitors (ICI) targeting programmed cell death protein-1 (PD-1) (nivolumab and pembrolizumab), PD-L1 (atezolizumab and durvalumab), or T lymphocyte-associated antigen 4 (CTLA-4) (ipilimumab) (Cloughesy et al., 2019; Filley et al., 2017; Kurz et al., 2018; Nayak et al., 2021; Omuro et al., 2018; Reardon et al., 2020; Schalper et al., 2019). Unfortunately, these ICI strategies did not show therapeutic efficacy in GB patients (Tomaszewski et al., 2019; Wang et al., 2024). More recently, preclinical studies have utilized adeno-associated virus (AAV) vectors to promote anti-tumor immunity. For example, cell-specific AAV vectors targeting endothelial cells in the CNS and expressing the lymphocyte recruiting cytokine LIGHT in the tumor vasculature, induced CD8 T cell infiltration that prolonged survival in murine GB (Ramachandran et al., 2023). In preclinical GB-murine models, AAV type 6 transduction of chemokine CXCL9-expressing astrocytes increased tumor infiltration of cytotoxic lymphocytes, when combined with anti-PD-1 immune checkpoint blockade (von Roemeling et al., 2024).

Although GB remains refractory to immunotherapy, tumors that are equipped with specific tumor antigens could be targeted by immune cells and therefore respond to immunotherapy (Nakagawa et al., 2023). Indeed, encouraging developments have been reported suggesting that the local administration of IL-12, a pro-inflammatory cytokine, can reinvigorate the immune system in the context of recurrent GB in humans (Chiocca et al., 2019). In these studies, the therapeutic effect of IL-12 was enhanced by injecting a replication-incompetent adenovirus vector directly into the tumor resection site. IL-12 expression was restricted by a drug-inducible promoter only activated following oral administration of 20 mg of the blood-brain barrier-permeable drug Veledimex (VDX) (Barrett et al., 2018; Zhang et al., 2015). This gene therapy extended the median survival to 17.8 months in recurrent GB patients without dexamethasone treatment, compared to a median survival of 8.14 months in historical controls. Despite the induction of anti-GB immunity by localized IL-12, patients still progressed over time, and a local increase in PD-L1 was observed (Chiocca et al., 2019). To counteract the induced immune suppression, a combined therapy of IL-12 and the ICI nivolumab was explored; however, this did not lead to extended survival in Phase II clinical trials (Chiocca et al., 2022). Instead of focusing on blocking the IL-12-induced immunosuppressive signaling imposed by tumor cells or the pro-tumoral TME (Agliardi et al., 2021; Chiocca et al., 2019), efforts could be redirected toward enhancing the activity of anti-tumor cells associated with GB tumors.

In GB patients, CD8 T cells are typically present in low numbers, representing only 0.6% of primary-derived tumor tissue (Friebel et al., 2020; Maddison et al., 2021; Nickl et al., 2023), and have a heterogenous phenotype (Wischnewski et al., 2023). The failure of immunotherapy for GB is partly attributed to the suppression of both the accumulation and anti-tumor functions of CD8 T cells (Lin et al., 2024). In patients with malignancies, CD8 T cells can become dysfunctional, or their cytotoxic functions become compromised (Nussing et al., 2020; Philip and Schietinger, 2022). It has emerged that some anti-tumor T cell responses require direct instructions from dendritic cells (DCs), that cross-present tumor-derived antigens (Ags) presented on their major histocompatibility complex (MHC) molecules (Zagorulya and Spranger, 2023). However, in GB, DCs themselves often become suppressed (Friedrich et al., 2023). Therefore, it is necessary to not only stimulate T cells directly but also activate DCs in the GB TME to promote the T cell activity needed for tumor control.

Here, we demonstrate that local administration of recombinant (r)IL-12 gives rise to intratumoral (i.t.) effector-like c CD8^POS^ T cells involved in GB regression. In addition to the direct stimulation of CD8^POS^ T cells by rIL-12, specialized DCs are recruited and activated at the tumor site, further enhancing the CD8^POS^ T cell activity. To support both the function of CD8^POS^ T cells and DCs during the rIL-12 inflammatory response, we aimed to establish a reservoir of 4-1BBL at the tumor. We selected this co-stimulatory molecule because both CD8^POS^ T cells and DCs express its receptor, 4-1BB. An AAV vector was demonstrated to enable the expression of 4-1BBL, mainly in reactive astrocytes within the peritumor region, enhancing the rIL-12-driven survival benefit. Our findings were predominantly tested using relevant syngeneic mouse models with intracranially (i.c.) engrafted CT-2A cells and further validated in GL261 and 005 tumors throughout the study.

## RESULTS

### Intratumorally administered rIL-12 prolongs survival of GB-bearing mice

IL-12 is a proinflammatory cytokine composed of two subunits, IL-12A (p35) and IL-12B (p40), which are covalently linked to form a bioactive IL-12p70 heterodimer complex (Watford et al., 2003). At the tumor site, IL-12 can promote anti-tumor immunity (Keppler et al., 2009; Kilinc et al., 2006), thereby altering the cellular composition of the TME (Rossari et al., 2023). In the context of primary human glioma, high or low expression levels of *IL12A*/B in the tumor did not predict the overall survival or GB patients, which showed a median survival of ∼2 years for each cohort (**Figure 1A**). Single-cell RNA sequencing (scRNAseq) analysis of human gliomas (**Table 1**), encompassing WHO grades II, III, and IV as well as both IDH-mutant and wild-type glioma patients (Louis et al., 2021), suggests that the limited predictive value of IL12A/B may be due to its consistently low expression levels across human gliomas (**Figure S1-A**). The transcript levels of *IL12A*/B in recurrent glioma did not vary from *de novo* tumors (**Figure S1-B**) and were independent of glioma grade (**Figure S1-C**). By processing multiple scRNAseq datasets across various murine GB models (CT-2A, GL261, 005) (**Table 1**) with different genetic and phenotypic profiles (**Table 2**), we confirmed a trend of low *Il12a/b* gene expression in the TME and mouse GB cell lines (**Figure S1-D-E**). The suppression of GB-associated IL-12 can be illustrated with *Il-12b^−/−^*mice, which lack endogenous IL-12 (Magram et al., 1996). Post-i.c. implantation of CT-2A-Firefly Luciferase (FLuc) GB cells, *Il12b^−/−^*mice did not exhibit a survival disadvantage compared to *Il12b^+/+^*mice (**Figure 1B**). FLuc was introduced into CT-2A cells using a lentiviral vector (LVV), enabling *in vivo* bioluminescence imaging (BLI) to monitor tumor engraftment and growth in the mouse brain (**Figure S2-A**). We further quantified IL-12p70 protein levels in the ipsilateral (tumor-implanted) hemispheres by analyzing the tumors of *Il12b^+/+^* and *Il12b^−/−^* mice (**Figure 1C**), concluding that IL-12 was expressed at low levels by the host (*Il-12b^+/+^ 1.2 fg* ± 0.341 and *Il-12b^−/−^ 0.8 fg* ± 0.027, mean ± SEM) and the tumor growth did not elicit an increase in rIL-12 expression in the CNS compared to the unaffected hemisphere.

**Figure 1.**
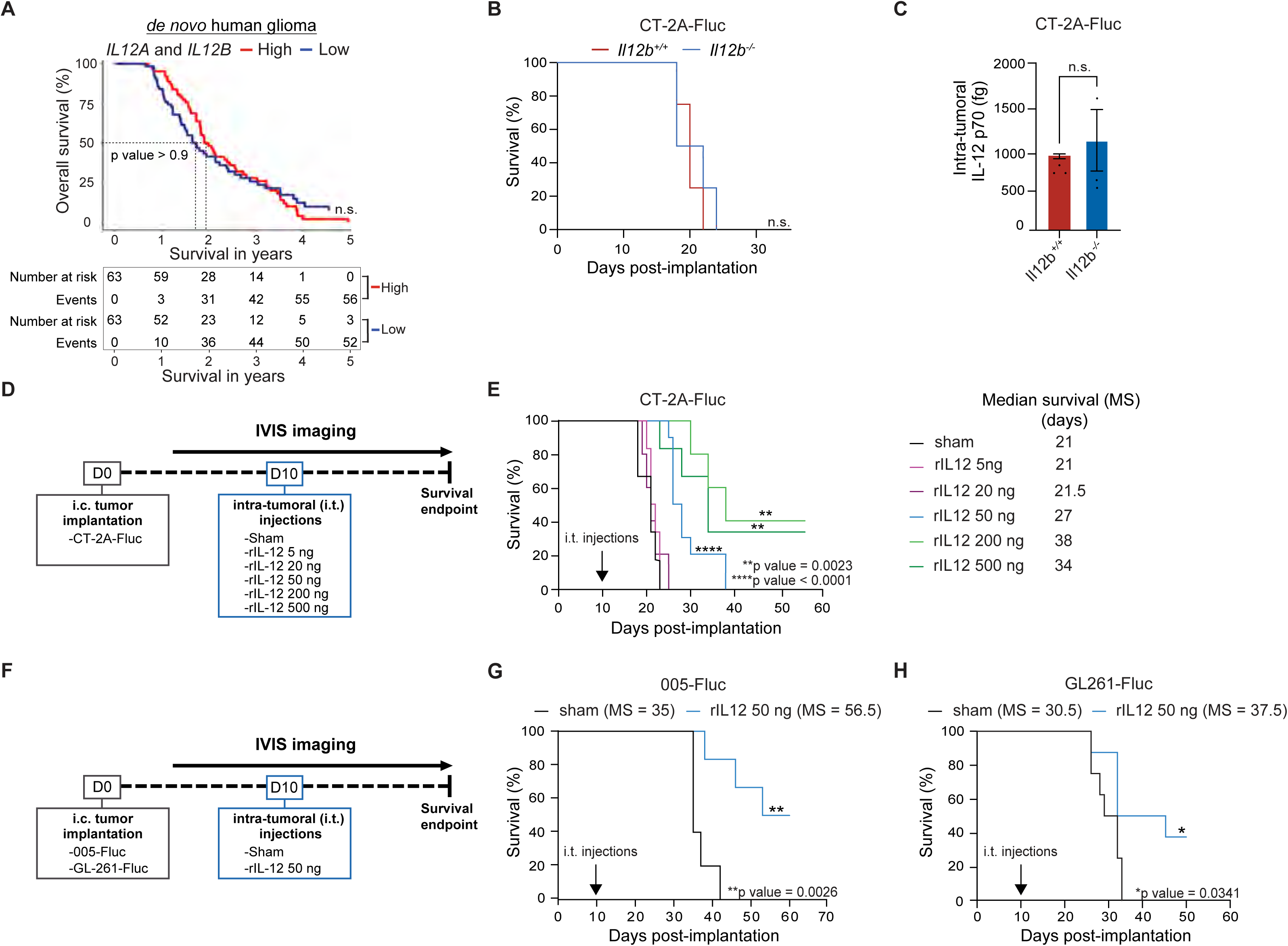
Intratumorally administered rIL-12 prolongs survival of GB-bearing mice. (A) *Survival probability of de novo human glioma differentiating high and low expressing IL12A andIL12B.* Kaplan-Meier survival curves showing the overall survival outcomes over a period of 5 years of a total of 63 glioma patients per group with high (red) or low (low) levels of *IL12A* and *IL12B* combined, each group had a median survival of ∼2 years (Miller et al., 2023). Samples in the top 33% in terms of expression of genes of interest (or module scores) were labeled as “high.” The bottom 33% were considered the “low” group. No differences were observed between groups. Log-rank (Mantel-Cox) test, p-value = 0.9, not significant (n.s). (B) *Survival curves of GB-bearing mice in Il12b^+/+^ and Il12b^−/−^ mice.* Kaplan-Meier survival curves showing no overall survival benefit of CT-2A tumor-bearing *Il12^+/+^* mice (red) compared to *Il12^−/−^*mice (blue) (n=5 mice per group) (median survival of 20 days and 18 days, respectively). Data represents at least two independent experiments. No differences were observed between the groups. Log-rank (Mantel-Cox) test, not significant (n.s). (C) *IL-12 protein levels in GB-containing brain hemisphere.* IL-12 levels femtogram (fg) were determined using Luminex in GB-bearing (CT-2A-FLuc) Il12^+/+^ (red) and Il12^−/−^ (blue) mice. Values were normalized to the non-tumor hemisphere. Data represents three independent experiments and are presented as the mean with SEM (error bars). Unpaired t-test, not significant (n.s). (D) *Schematic illustration of the in vivo experimental set-up.* CT-2A-FLuc GB (100.000) cells were injected intracranially (i.c.) into the left striatum on day 0. Starting on day 7, tumor growth was monitored every 3 to 4 days by in vivo imaging system (IVIS). Based on FLuc levels on days 7 and 10, mice with a similar tumor size were allocated to sham (Fc control) or rIL-12 (recombinant IL-12 conjugated to Fc) -treatment groups (ranging between 5 to 500 ng) on day 10. Sham and rIL-12 solutions were administered intratumorally (i.t.). (E) *Survival of CT-2A tumor-bearing mice with rIL-12 treatment*. Kaplan-Meier survival curves show an rIL-12 dose-response study in mice implanted with (CT-2A-FLuc) GB cells, compared to sham treatment. The arrow indicates the timepoint of i.t. injections of sham or rIL-12. The median survival was significantly increased when GB mice were treated with either 50 ng, 200 ng, or 500 ng (27, 38, and 34 days respectively; **p = 0.0023) compared to sham control 20 days. Median survival for 5 ng and 20 ng rIL-12 were 22.5 and 21.5 days respectively. (50 ng sham, n= 6; 5 ng rIL-12, n=6; 20 ng rIL-12, n=5; 50 ng rIL-12, n=10; 200 ng rIL-12, n=5, and 500 ng rIL-12, n=6). Data represents three independent experiments and were analyzed using the Log-rank (Mantel-Cox) test, **p < 0.01 ****p < 0.0001. (F) *Schematic illustration of the in vivo experimental set-up.* 005-FLuc and GL261-FLuc GB cells (n=100,000) were injected i.c. into the left striatum on day 0. Starting on day 7, tumor growth was monitored every 3 to 4 days by IVIS BLI. Based on FLuc levels on day 7 and day 10, mice with similar tumor sizes were allocated to 50ng sham or 50ng rIL-12 treatment groups on day 10. Sham and rIL-12 were administered i.t. (G) *Survival of rIL-12 treated 005-FLuc bearing mice.* Kaplan-Meier curves displaying the percentage of survival of 005-FLuc bearing mice (100,000 cells at the time of injection) with treatment on day 10 post-tumor injection, comparing i.t. injection of 50 ng rIL-12 (blue) to sham control (black) (n=5-6 mice per group). Approximately ∼50% of the rIL-12 treated mice stayed healthy over 50 days (**p=0.0026). Data represents at least two independent experiments. Data were analyzed using Log-rank (Mantel-Cox) test, **p < 0.01. (H) *Survival of rIL-12 treated GL261-FLuc bearing mice.* Kaplan-Meier curves displaying the percentage of survival of GL261 bearing mice (100,000 cells at the time of injection) with treatment on day 10 post-tumor injection comparing i.t. injection of 50 ng rIL-12 (blue) to sham control (Fc-black) (n=5-6 mice per group). Approximately ∼40% of the rIL-12 treated mice stayed healthy over 50 days (*p=0.0341). Data represents at least two independent experiments. Data was analyzed using Log-rank (Mantel-Cox), *p < 0.05.

To increase IL-12 at the tumor site to therapeutically effective levels, we i.t. injected a single dose testing various doses (5ng, 20ng, 50ng 200ng and 500ng) of murine recombinant IL-12 conjugated to Fc (hereafter referred to as rIL-12) ranging from 5 ng to 500 ng ten days after CT-2A-FLuc engraftment (**Figure 1D**) and compared survival outcomes to mice i.t. injected with sham (Fc without the IL-12 fusion). On day 7, prior to i.t. rIL-12/sham injections, tumors had a similar size across groups based on BLI (**Figure S2-B**), while post-treatment, the BLI signals were altered depending on the treatment (**Figure S2-C**). Together with the evaluation of body weight (**Figure S2-D**), BLI measurements aided us in capturing different treatment responses of the GB-bearing mice over time. Some GB-bearing mice did not respond to treatment (non-responders), while others showed reduced tumor size but eventually died from the tumor (treatment-responders). A third group of treated mice survived the GB implantation (treatment-survivors). Among the treated non-responders, mice exhibited similar outcomes to the sham group, characterized by a steady increase in tumor size and a decline in body weight, indicative of poor health. This response pattern included all GB-bearing mice treated with 5 ng and 20 ng rIL-12. Notably, a cohort of mice, specifically 64% and 33% of the GB-bearing mice treated with 50 ng and 500 ng of rIL-12, respectively, exhibited a similar response as the sham-treated GB-bearing mice. Treatment responders showed delayed tumor growth in response to rIL-12 treatment. A slower increase in the bioluminescence (BLI) signal and a minimal decrease in body mass were observed in the treatment responders compared to the non-responders. The proportion of treatment responders was 36%, 60%, and 33% among the GB-bearing mice treated with 50 ng, 200 ng, and 500 ng of rIL-12, respectively. The treatment-survivors demonstrated favorable outcomes with rIL-12 treatment; 40% and 33% of GB-bearing mice treated with 200 ng and 500 ng rIL-12, respectively, displayed tumor regression concomitant with stable body weight. The mice in this response pattern continued to live for at least 60 days without apparent health concerns (**Figure S2-D**).

The varying responses of mice to different doses of rIL-12 resulted in different survival outcomes. Mice treated with rIL-12 50 ng showed 6 days of improved median survival compared to sham-treated mice. Mice treated with 200 ng and 500 ng benefited 17 and 13 days, respectively (**Figure 1E**). Pathology evaluation of tumor-implanted mice was performed using Hematoxylin & Eosin (H&E) staining, and tumor sizes were quantified across the tested rIL-12 dosages (**Figures S2-E-F**). The i.t. administration of 50 ng rIL-12 was considered the optimal dose for subsequent experiments. While the dosage significantly increased median survival, it was not sufficient to cure tumor-implanted mice, thereby closely reflecting survival outcomes seen in IDH-wild type recurrent GB patients i.t. treated with adenoviral gene therapy delivering IL-12 (Chiocca et al., 2022). Additionally, the 50 ng rIL-12 dosage was chosen throughout our study because it minimizes the risk on rIL-12-associated toxicity and allows for complementary therapies to further enhance the rIL-12-driven survival effect. Systemic toxicity of 50 ng rIL-12 was assessed by comprehensive blood chemistry analysis in mice. No significant elevation in liver biomarkers—including albumin, alkaline phosphatase (ALP), alanine transaminase (ALT), calcium, cholesterol, creatinine, blood urea nitrogen (BUN), globulin, glucose, and phosphorus—was detected after i.t. injection of rIL-12 (50 ng) compared to sham (**Figure S2-G**). The blood tests we conducted indicated no systemic toxicity upon localized administration of rIL-12, in contrast to the previous reported effects of systemic recombinant IL-12 administration (Leonard et al., 1997).

Next, we validated the efficacy of the rIL-12 therapy with the 005-GB cell line, known for its diffuse tumor growth, offering a closer representation of human glioma (Marumoto et al., 2009) as well as the GL261 GB model (**Figure 1F**). I.t. rIL-12 treatment (50 ng) administered 10 days post-tumor injection was effective in 005-FLuc GB-bearing mice treated with rIL-12 and had a 21.5-day improved median survival outcome compared to sham. Importantly, 50% of the mice survived for over 50 days (**Figures 1G and S2-H**). In the GL261 model, the median survival was 37.5 days in the rIL-12-treated mice compared to 30.5 days in sham-treated mice. Approximately 40% of rIL-12 treated tumor-bearing mice survived for over 50 days (**Figures 1H and S2-I**).

Overall, the survival outcomes indicate that i.t. administered rIL-12 increases the lack of endogenous IL-12 in GB tumors and achieves therapeutic effects with 50 ng dosage in multiple syngeneic mouse GB models.

### Identifying the cell types within the GB tumor microenvironment that can trigger an rIL-12-mediated response

IL-12p70 binds to the dimeric receptor composed of the IL-12 receptor β1 (IL12Rβ1) and the IL-12 receptor β2 (IL12Rβ2) subunits, leading to phosphorylation of Tyr693 on the receptor-associated STAT4 transcription factor (**Figure 2A**) (Liu et al., 2005). This phosphorylation promotes STAT4 dimerization, thereby initiating pro-inflammatory signaling (Landoni et al., 2024; Watford et al., 2003). To identify the cells capable of an IL-12p70-mediated anti-GB effect, we analyzed available human glioma scRNAseq datasets (**Table 1**) for the expression of relevant genes (*IL12R*β*1, Il12R*β*2, and STAT4*). Immune cells expressed all three markers in contrast to malignant (tumor) cells, oligodendrocytes, and stromal (vascular) cells which expressed low-to-no levels. Markers were predominantly co-expressed in tumor-associated Natural Killer (NK) cells, CD4 and CD8 T cells (NK/T cells) in *de novo* and recurrent human gliomas (**Figures 2B and S3-A**).

**Figure 2.**
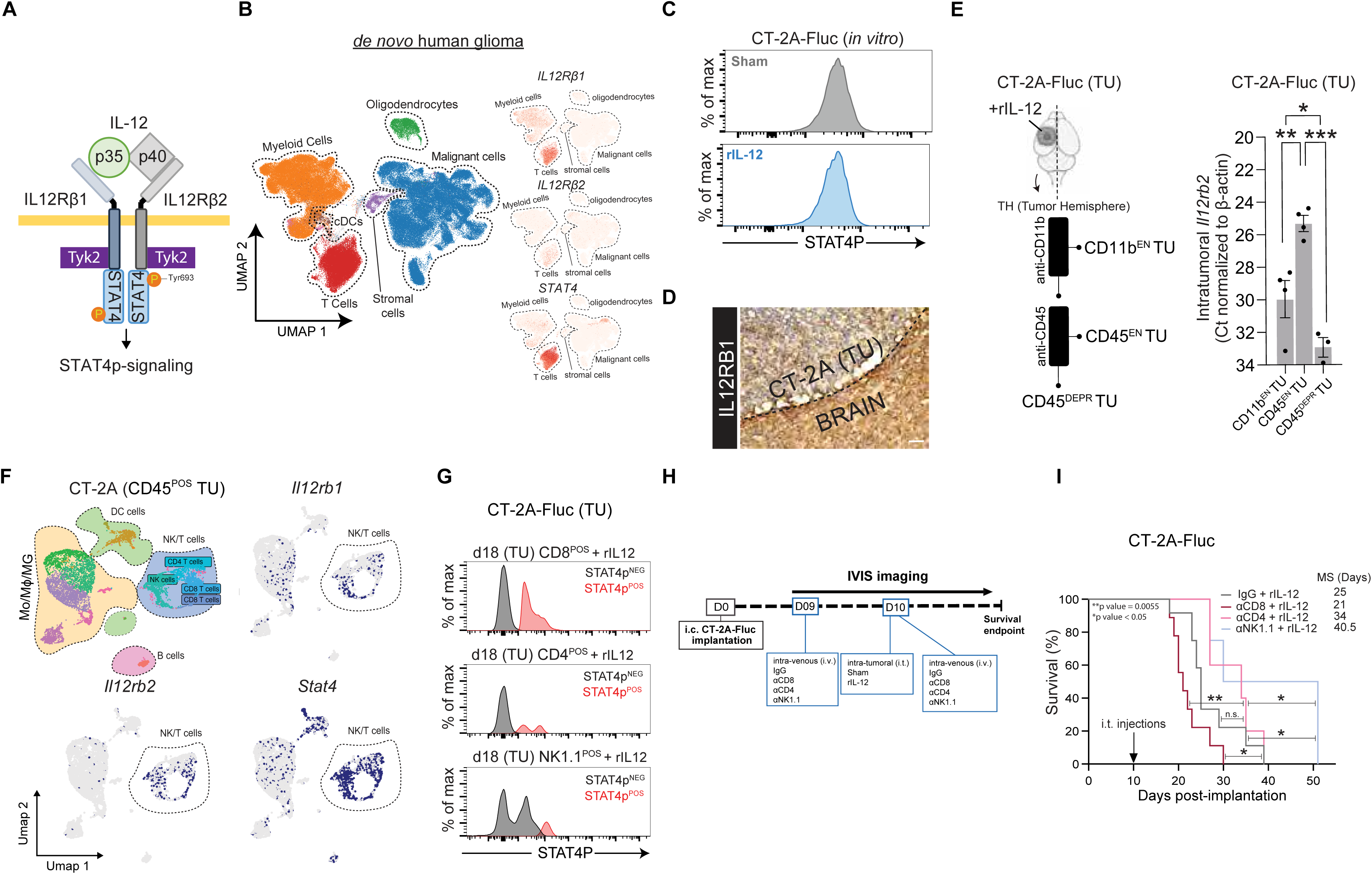
Identifying the cell types within the GB tumor microenvironment that can trigger an rIL-12-mediated response. (A) *Schematic illustration of IL-12R signaling.* IL-12 has two subunits p35 and p40 which bind to IL12 receptor beta (β) 1 and IL12Rβ2 respectively to initiate downstream signaling through tyrosine kinase 2 and phosphorylation of Stat4 proteins. (B) *IL12R*β*1, IL12R*β*2 and STAT4 expression in immune cell populations in de novo human glioma.* Distinct cell types were clustered, annotated and visualized with a high-resolution color-coded UMAP projection (Miller et al., 2023). To visualize IL12Rβ1, IL12Rβ2 and STAT4 expression single cell feature plots were used to compare transcript levels in the immune cell population. (C) *Representative flow cytometry plots of pSTAT4 protein expression in CT-2A-FLuc cultured cells.* CT-2A-FLuc cells were exposed to sham or rIL-12 for 24 hrs, pSTAT4 expression was measured by flow cytometry and no differences were observed. (D) *Positive IL12rb1 receptor staining in brain tissues implanted with a CT-2A tumor.* Immunohistochemistry of IL12rb1 positive cells (brown) in the TME of the CT-2A tumor (TU) cells (blue) (magnification 40x; scale bar = 100 µm). (E) *Decoupling Il-12R expressing Mo/M /MG cells from other immune cells in GB-bearing mouse brains.* A schematic display shows the sequential method used to isolate CD11b enriched (EN) tumor (TU), CD45^EN^ TU and CD45 deprived (DEPR) TU derived from the tumor hemisphere (TH) of mouse brains post-rIL-12 treatment (left). I.t. *Il12rb2* expression was analyzed eight days after rIL12-treatement. *Il12rb2* was expressed significantly higher in CD45^EN^ TU compared to CD11b^EN^ TU (p-value = 0.0043) and CD45^DEPR^ TU cells (p-value = 0.0001). CD45^EN^ TU *Il12rb2* levels were significantly higher (p-value = 0.0136) than CD45^DEPR^ TU cells. Data represent CT values normalized to β-actin. Data represent three independent experiments and are presented as the mean with SEM (error bars). Data were analyzed using one way ANOVA, *p < 0.05, **p < 0.01, ***p < 0.001 (right). (F) *Expression of IL-12rb1 and IL-12rb2 and pStat4 in immune cell populations.* scRNAseq datasets of CD45^POS^-sorted tumor cells derived from mouse GB TU (tumor) (CT-2A, n=3) were analyzed (Tomaszewski et al., 2022). Distinct cell types were clustered, annotated, and visualized with a high-resolution color-coded UMAP projection. To visualize IL-12rb1, IL-12rb2 and pStat4 expression in different datasets, single-cell feature plots were used to display the expression in NK/T cluster (marked in dotted lines). (Chen et al., 2023; Pombo Antunes et al., 2021; Tomaszewski et al., 2019). (G) *pSTAT4 levels in CD8, CD4 T cell and NK cell populations.* CD8, CD4 T cells and NK1.1 cells were isolated from CT-2A-FLuc tumor (TU) bearing, rIL-12 treated mice on day 18 post-tumor implantation. Representative flow plots showed pSTAT4^POS^ and pSTAT4^NEG^ levels as the percentage of max. All three cell types express pSTAT4. (H) *Schematic illustration of the T cell depletion strategy at the tumor site.* On day 0, 100,000 GB cells (CT-2A-FLuc) were implanted i.c. into the left striatum. Anti-CD8 or IgG control was injected intravenously (i.v). on day 9 (50 µg, retro-orbitally). On day 10, mice were injected with 50ng rIL-12 or sham control i.t. at the tumor (TU) site and anti-CD8 or IgG control was injected i.v. (100 µg, retro-orbitally) to deplete endogenous CD8^POS^ T cells systemically. (I) *Importance of CD8^POS^ T cell recruitment for survival benefit in anti-GB therapy with rIL12.* Kaplan-Meier curves showing survival outcome of tumor-bearing mice injected with IgG and rIL-12 (grey), with CD8 depletion (anti-CD8) and rIL-12 (solid red), with CD4 depletion (anti-CD4) and rIL-12 (solid pink) and with NK cells depletion (anti-NK1.1) and rIL-12 (solid light blue) (n = 6-8 mice per group). IgG control treated with rIL-12 had a median survival (MS) of 25 days, whereas anti-CD8 had a median survival of 21 days, anti-CD4 34 days, and anti-NK 40.5 days. IgG control did not differ compared to anti-CD4 but had a significantly improved survival compared to anti-CD8 (p-value = 0.0372), and anti-NK (p-value = 0.0479). anti-CD8 had significant improved median survival compared to anti-NK (p-value = 0.0122) and anti-CD4 (p-value = 0.0055). Data represent two independent experiments and were analyzed using Log-rank (Mantel-Cox) test, *p < 0.05 **p < 0.001.

Consistent with these findings, we did not expect rIL-12 to have a direct effect on GB tumor cells. In fact, *Il12rb1* was expressed only at low levels *in vitro*, and *Il12rb*2 was not detected in the murine tumor cells (**Figure S3-B**). Additionally, upon *in vitro* exposure to 50 ng rIL-12 or sham, GB cell proliferation was not affected over the course of 5 days (**Figure S3-C**), nor did it alter the Tyr693 STAT-4 phosphorylation (pSTAT4) levels altered (**Figures 2C and S3-D**). This was confirmed in brain sections of tumor-bearing mice, where IL12Rβ1 expression was minimally present within the CT-2A tumor and predominantly localized at the tumor periphery (**Figure 2D**), as compared to the negative control (**Figure S3-E**).

To screen the immune cell populations involved in rIL-12-mediated anti-GB immunity, we enzymatically dissociated rIL-12-treated CT-2A-FLuc tumors and serially enriched for immune cell populations from the TME through affinity columns (anti-CD11b and anti-CD45) (**Figure 2E - Left**), followed by qRT-PCR analysis for *Il12rb2* expression (**Figure 2E - Right**). *Il12rb2* transcript levels in CD45 enriched (EN) tumor (TU) cells (immune cells derived from tumor after anti-CD11b depletion and anti-CD45 column enrichment) were 30.80-fold and 192.72-fold higher compared to CD11b^EN^ TU (immune cells derived from tumor cells after anti-CD11b column enrichment) and CD45 deprived (DEPR) TU cells (cells derived from tumor after anti-CD11b and anti-CD45 depletion) respectively (**Figure 2E - Right**). In line with the human glioma scRNAseq analysis, we found that rIL-12 sensing CD45^EN^ TU cells from murine GB primarily consisted of NK/T cells. We concluded this by screening for *Ifng* transcript levels, as this cytokine is primarily secreted by activated NK/T cells (Tau et al., 2001). This was validated through immune cell analysis of available scRNAseq datasets (**Table 1**) from mice implanted with different tumor cell lines (CT-2A, 005, and GL261) (**Figure S3-F**). Similarly, *Ifng was* mainly detected in the CD45^EN^ TU fraction of the CT-2A TME following rIL-12 treatment (45.65- and 64.21-fold higher than CD11b^EN^ and CD45^DEPR^, respectively) (**Figure S3-G**).

We confirmed that the expression profiles of relevant IL-12R genes (*Il12rb1*, *Il12rb2*, and *Stat4*) are similar between human gliomas (**Figure 2B**) and the murine GB cell lines (CT-2A, GL261, and 005) (**Figures 2F and S3-H**). The NK/T cluster including NK cells, CD4 T cells, and CD8 T cells exhibited the highest expression of *Stat4* (20-40%) compared to the other cells (∼1-5%) in murine GB models (GL261, CT-2A and 005) (**Figure S3-I**). Based on pSTAT4 levels, we confirmed that post-rIL-12 treatment, all NK/T cell types have the potential to activate an IL-12 mediated immune response against the GB cells (**Figure 2G**). pSTAT4^POS^ cells were found in 12.6% of the CD8 T cells, 3.7% of the CD4 T cells, and 2.3% of the NK1.1 cells in the TME of a CT-2A tumor treated with rIL-12 (**Figure S3-J**). To identify which NK/T cell types contribute to the anti-GB effect of i.t. administered rIL-12, we depleted CD4, CD8, or NK cells systemically in rIL-12-treated GB mice (**Figure 2H**). Survival analysis of CT-2A tumor-bearing mice treated i.t. with rIL-12 and intravenously (i.v.) with anti-CD4, anti-CD8, or anti-NK1.1 revealed that only CD8 T cell depletion significantly reduced the rIL-12-mediated anti-GB response compared to the control (IgG) (**Figure 2I**). Conversely, anti-NK1.1 treated mice with rIL-12 showed a 37% improved survival compared to control (IgG), implying that NK cells can suppress the effect of rIL-12 therapy. Anti-CD4 treatment had no effect on the GB survival post-rIL12 treatment. This was confirmed by increased tumor growth based on BLI measurements and a reduction in weight with anti-CD8 as compared to IgG control in rIL-12-treated GB mice (**Figure S3-K**). No differences in BLI were observed for anti-CD4 and anti-NK1.1 depleted mice compared to IgG control. NK1.1 cell-depleted mice gained weight, while CD4 T cell-depleted mice lost weight over time (**Figure S3-L**). We next verified if the TME of an rIL-12-treated CT-2A tumor was changed upon i.v. anti-CD8 administration with flow cytometry of CD11b^EN^ and CD45^EN^ TU cells. CD8^POS^Thy1.2^POS^ T cells were detected only in the CD45^EN^ TU cell fraction of our non-depleted control (IgG) and not in the mice that received i.v. anti-CD8 injections (**Figure S3-M**). CD11b^EN^ TU cells did not contain CD8^POS^Thy1.2^POS^ T cells. CD8 T cell depletion in an i.t. rIL-12 GB-bearing mouse can also be monitored by analyzing the periphery. Indicating that the CD8 T cells were depleted not only in the brain but also in the spleen (**Figure S3-M**). Additionally, on days 11 and 18 post-tumor implantation (1 and 8 days after the last i.v. injection with anti-CD8, respectively) lower *Cd8b* levels in the blood were observed due to the depletion, while this was not the case on day 7 (three days before rIL-12 treatment and two days before the first injection with anti-CD8) (**Figure S3-N**).

Taken together, our data demonstrates that CD8 T cells within the TME of GB tumors are the key effectors driving tumor reduction during a rIL-12-mediated response.

### Identifying GB-associated DCs with the potential to stimulate CD8^POS^ T cells during IL-12 immunity

While *Il12^+/+^* and *Il12^−/−^* mice showed no difference in survival (**Figure 1B**), we hypothesized that cells in the tumor vicinity may still express IL-12, albeit insufficiently for anti-tumor immunity. Here, we investigated which cell types produce IL-12, their additional anti-tumor capabilities for aiding CD8^POS^ T cells, and their representation at the tumor site. *Il12b* was predominantly expressed by immune cells in the CT-2A mouse model (**Figure S4-A**). In the human glioma (*de novo* and recurrent) scRNAseq dataset, we found that *IL-12B* was mainly expressed in the DC cluster (**Figure S4-B**). DCs are known to instruct tumor antigen-reactive T cells, including CD8^POS^ T cells, to proliferate and activate their cytotoxic machinery when necessary (Fu and Jiang, 2018). When CD8^POS^ T cells isolated from GL261-tumors were co-cultured with naïve DCs, no effect of rIL-12 could be observed, as assessed by IFN-γ secretion (**Figure S4-C**). However, when DCs were stimulated to cross-present the GL261-neoepitope peptide mImp3 (Schaettler et al., 2023) before exposure to rIL-12, the mouse tumor-derived CD8^POS^ T cells secreted 7-times higher levels of IFN-γ compared to sham control (**Figure 3A**). This indicates that rIL-12 acts as a stimulatory cytokine for GB-associated CD8^POS^ T cells recognizing the DC MHC-I-tumor antigen peptide complex via their T cell receptors. Hence, we screened scRNAseq datasets of human glioma tissue (**Figure S4-D-E**) and murine GB models (CT-2A and GL261) (**Figures 3B-C and S4-F-G**) to identify DCs in the TME that might be involved in tumor antigen cross-presentation at the tumor site and whether they are equipped to modulate CD8^POS^ T cell activity and/or proliferation through co-stimulatory factors and/or cytokine production. Of note, the murine 005 dataset (**Table 1**) contained no relevant DC information. In our analysis, *H2-D1* encoding H-2Db, an MHC-I class that binds our mImp3 peptide (Johanns et al., 2016), was highly expressed by a specific DC state in the CT-2A and GL261 models (**Figures 3B and S4-F**) that was distinct from the conventional classical (c)DC1/2, and pDCs (**Figures 3C and S4-G**). This DC state encodes high levels of genes important for MHC-I antigen processing and cross-presentation (such as *Psme2* and *TapbpI*) but showed lower expression of genes (including *Cd74*) involved in MHC-II-mediated antigen presentation. Interestingly, these MHC-I-expressing DCs co-expressed high levels of CD8^POS^ T cell activity modulating factors, including *Il12b* (encoding IL-12p40), inhibitory CD8^POS^ T cell factors, including *Cd274* (encoding PD-L1), and co-stimulatory factors, including *Tnfsf9* (encoding 4-1BBL) (**Figures 3B and S4-F**). The DCs of interest have an *Il2b* signature (**Figures 3C-D and S4-G-H**) and also express migratory markers, including *Fscn1, Ccr7*, which are important in facilitating DC migration to tumor-draining lymph nodes to shuttle between lymph nodes and *Ccl22,* involved in interactions between T regulatory cells and DCs (Liu et al., 2021) (**Figure 3E and S4-I**). Drawing conclusions about this cell type in human glioma datasets is challenging however due to the low number of MHCI-expressing DCs in these tumors (**Figures S4-D-E**). Specifically, in the *de novo* human glioma datasets, no *CCR7* expression in the DC cluster was observed. In one recurrent human GB dataset that contained *CCR7* expressing DCs, we also observed MHC-I components, stimulatory factors including *IL-12B,* inhibitory factors (e.g., *CD274*), and co-stimulatory factors (e.g., *CD80*) that match our murine findings (**Figure S4-E**). We hypothesize that the detection of CCR7^POS^ DCs in the second human recurrent dataset is due to its enrichment for immune cells prior to scRNAseq processing, whereas the other dataset did not perform this enrichment.

**Figure 3.**
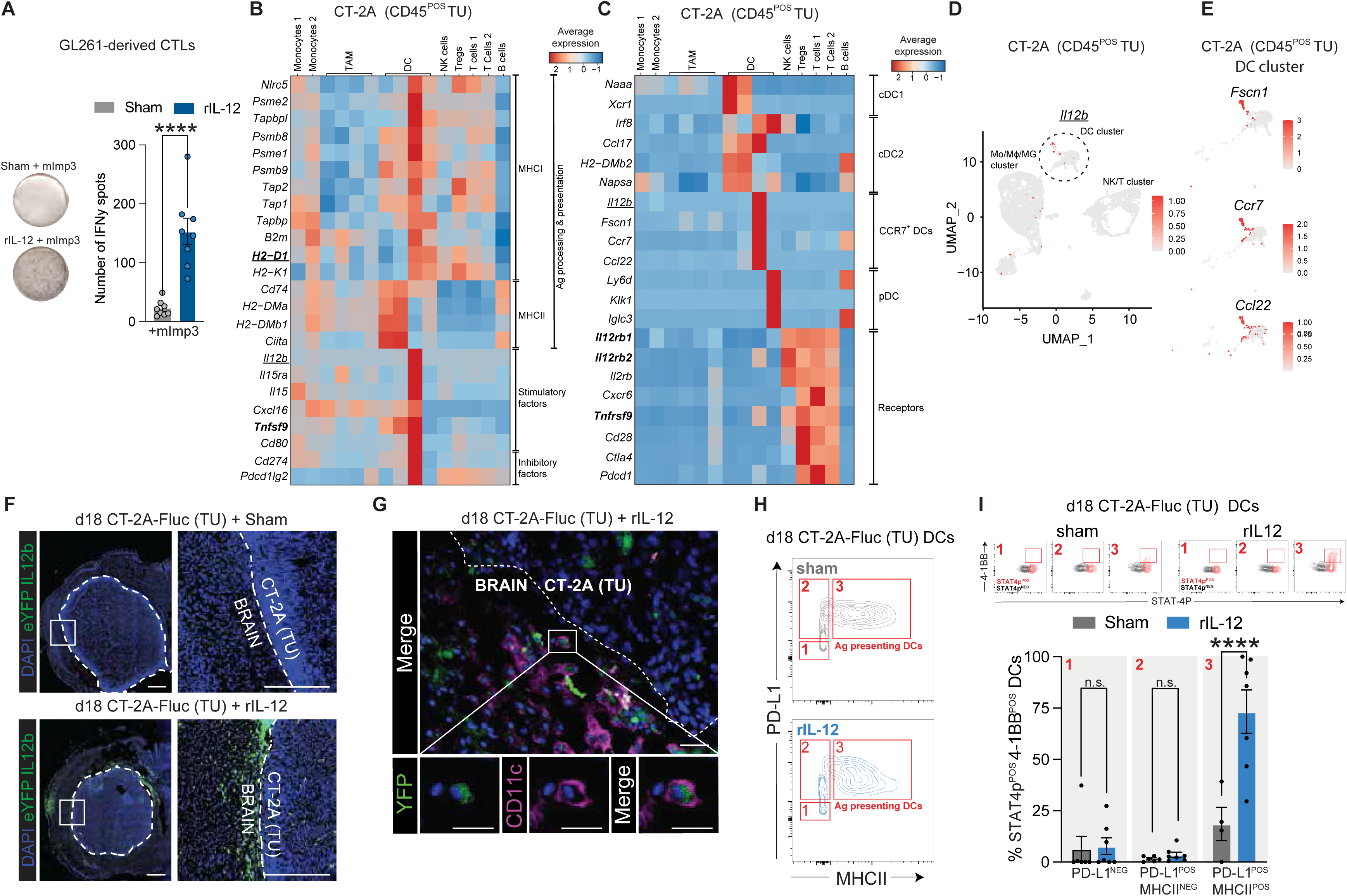
Identifying GB-associated DC states with the potential to stimulate CD8^POS^ T cells during IL-12 immunity. (A) *Quantification of IFN-*γ *activation of co-cultured CD8^POS^ T cells with naïve splenocytes.* Representative images of the IFN-γ Elispot assay demonstrate an increased number of spots when CD8POS T cells isolated from a GL261 primary CD8^POS^ T cells, isolated from GL261-bearing mice brain, were co-cultured with naïve splenocyte-derived splenocytes and exposed to mImp3, a GL261-specific neopeptide, for 24 hrs. This increase was observed only in the presence of recombinant IL-12 (rIL-12) and not with the sham control. The accompanying bar graph quantifies the number of spots (mean number of spots: 22 for sham; 153 for rIL-12). Each spot corresponds to an IFN-γ-releasing cell. Data represent two independent experiments and are presented as the mean with SEM (error bars). Data were analyzed using an unpaired t-test, ****p < 0.0001. (B) *Regulatory factors of interest for CD8^POS^ T cell differentiation by DCs*. Heatmap showing co-expression of genes that are generated by DCs in TME involved in tumor antigen cross presentation (Tomaszewski et al., 2022) and modulate CD8^POS^ T cell activity through stimulatory factors and/or inhibitory factors. *H2-D1* encoding for H-2Db, an MHC-I class was expressed by a specific DC state in the CT-2A model. MHC-I-expressing DCs co-expressed high levels of CD8^POS^ T cell activity modulating factors, including *Il12b* (encoded by IL-12p40), inhibitory factors, including *Cd274* (encoding for PD-L1), and co-stimulatory factors, including *Tnfsf9* (encoding for 4-1BBL). (C) *IL-12 expression profile matches CCR-7^POS^-DCs*. IL12b was highly expressed in subcluster CCR-7^POS^-DCs which had marked expression of migratory factors *Fsn1, Ccr7* and *Ccl22* (Tomaszewski et al., 2022). Il12b cells expressing its receptors were found to be expressed in the NK/T cell clusters. Plasmacytoid DC (pDC). (D) *IL-12 expression restricted to the state of the DCs.* UMAP clustering shows expression of IL12b in distinct population of the DC cluster in CT-2A (CD45^POS^) TU (Tomaszewski et al., 2022). (E) *Visualization of CCR-7^POS^-DCs in scRNAseq dataset*. The cells positive in panel D (marked with a dotted line) match with the dendritic-specific markers *Fscn1, Ccr7,* and *Ccl22* (Tomaszewski et al., 2022). (F) *IL-12 expressing cells in the TME of CT2A-FLuc GB*. High numbers of YFP expressing cells were observed in both the CT-2A tumor border (white dotted line) as well as the tumor itself in IL-12b-YFP reporter mice treated on day 10 post-tumor implantation with sham control or 50 ng rIL-12. Mice were sacrificed on day 18 post-tumor implantation (4x magnification, scale bar = 50 µm). (G) *CD11c^POS^ IL-12 expressing DCs are recruited to the GB TME*. DCs were stained for CD11c and expressed YFP (10x magnification, scale bar = 10 µm; 40x magnification, scale bar = 50 µm). (H) *Enhanced expression of antigen-presenting cells after IL-12 treatment.* Contour plots of MHC-II expression versus PD-L1 in DCs on day 18 post-tumor implantation treated with sham (grey) or rIL-12 (blue) showing PD-L1^NEG^ (panel 1), PD-L1^NEG^MHC-II^POS^ (panel 2), PD-L1^POS^MHC-II^POS^ (panel 3), representing activated DCs. (I) *Quantification of the percentage of pSTAT4^POS^ cells*. Cells were pre-gated for 4-1BB versus pSTAT4 comparing sham and rIL-12 treated conditions (top). Quantification of pSTAT4^POS^ expressing cells in 4-1BB^POS^ DCs showed an increased expression of 54.71% (SEM ± 15.41) in cells pre-gated for PD-L1^POS^MHC-II^POS^ (panel 3) were found in rIL-12 treated cells compared to sham (n=6-7 mice per group) (bottom). Data represents two independent experiments and are presented as the mean with SEM (error bars). Data were analyzed using multiple comparison two way ANOVA, ****p < 0.0001, not significant (n.s.).

Based on our scRNAseq analysis, *Il12b*^POS^ DCs were found equipped with IL-12R machinery (mainly *Il12rb2*) and thus have the potential to be affected by rIL-12 similar to the CD8^POS^ T cells (**Figure 3C and S4-G**). Therefore, we assessed their involvement in rIL-12-mediated immunity at the GB site in our preclinical mouse model. Using IL-12b^YFP^ reporter mice (Reinhardt et al., 2006) to label *Il-12b*^POS^ DCs, we observed an accumulation of IL-12b^YFP^ cells at the CT-2A tumor site upon i.t. rIL-12 injection, but not when administering sham (**Figures 3F and S4-J**). Tumor-associated IL-12b^YFP^ positive cells were confirmed to be DCs as they co-expressed CD11c (**Figure 3G**). Next, we verified the *Il12b*^POS^ DC activity upon rIL-12 treatment of a GB tumor. Tumor-interacting and antigen-presenting CD11c^POS^ DCs in our CT-2A model were identified by the PD-L1 and MHC-II markers, respectively (**Figure 3H**). *Il12b*^POS^ DCs expressing 4-1BB (*Tnfrsf9*) (**Figures 3C and S4-G**) dominated the PD-L1^POS^MHC-II^POS^ population (**Figure 3I - TOP**). These cells also displayed most of the rIL-12 reactivity based on pSTAT4, the downstream signaling event of the IL-12 receptor pathway (Bacon et al., 1995). rIL-12 treatment resulted in a 54.71% ± 15.41 (mean difference ± SEM) increase of pSTAT4^POS^4-1BB^POS^ DCs at the tumor site compared to sham (**Figure 3I – BOTTOM**).

Taken together, our findings suggest that DCs accumulate and are activated at the tumor site by exogenous rIL-12. These intratumoral-affected DCs have distinct signatures compared to other DCs, have migratory markers, and properties that can modulate the CD8^POS^ T cell response, including providing stimulatory signals (e.g., co-stimulatory molecules and cytokines) to tumor-antigen-targeting CD8^POS^ T cells.

### Intratumoral injected rIL-12 increases the number of effector-like CD8^POS^ T cells at the tumor site

We have shown that the survival benefit from i.t. rIL-12 is driven by CD8^POS^ T cells and that local DCs may provide the necessary signals to guide their activity. Here we investigate whether rIL-12-induced, tumor-associated CD8^POS^ T cells have the potential to sense cues from rIL-12-activated DCs to enhance their functionality. CD8^POS^ T cells were present at the periphery of the CT-2A tumor i.t. treated with either rIL-12 or sham (**Figures 4A and S5-A**). Multiparametric flow cytometry was performed on CD45^EN^ TU cells to quantify CD8^POS^ T cells at the tumor hemisphere. rIL-12 treatment showed a 10-fold increase in CD8^POS^ T cells compared to sham-treated GB mice (**Figure 4B**). CD8^POS^ T cell accumulation in the tumor appeared to be tumor-targeted, as we could not detect CD8^POS^ T cells in the contralateral hemisphere of both rIL-12 or sham-treated mice. Interestingly, the number of CD8^POS^ T cells was consistently low in our sham-treated tumors, underscoring the immunosuppressive nature of the CT-2A model (Khalsa et al., 2020). This conclusion is further supported by evaluating survival of mice that received sham treatment after CT-2A-implantation comparing CD8^POS^ T cell-depleted and non-depleted mice, no differences in survival were found (**Figure S5-B**), and no differences in BLI or mice weights measured over time between groups (**Figure S5-C**). This was also observed in the human glioma survival analysis as high CD8a expression did not result in improved survival outcomes compared to CD8a-low-expressing gliomas (**Figure S5-D**).

**Figure 4.**
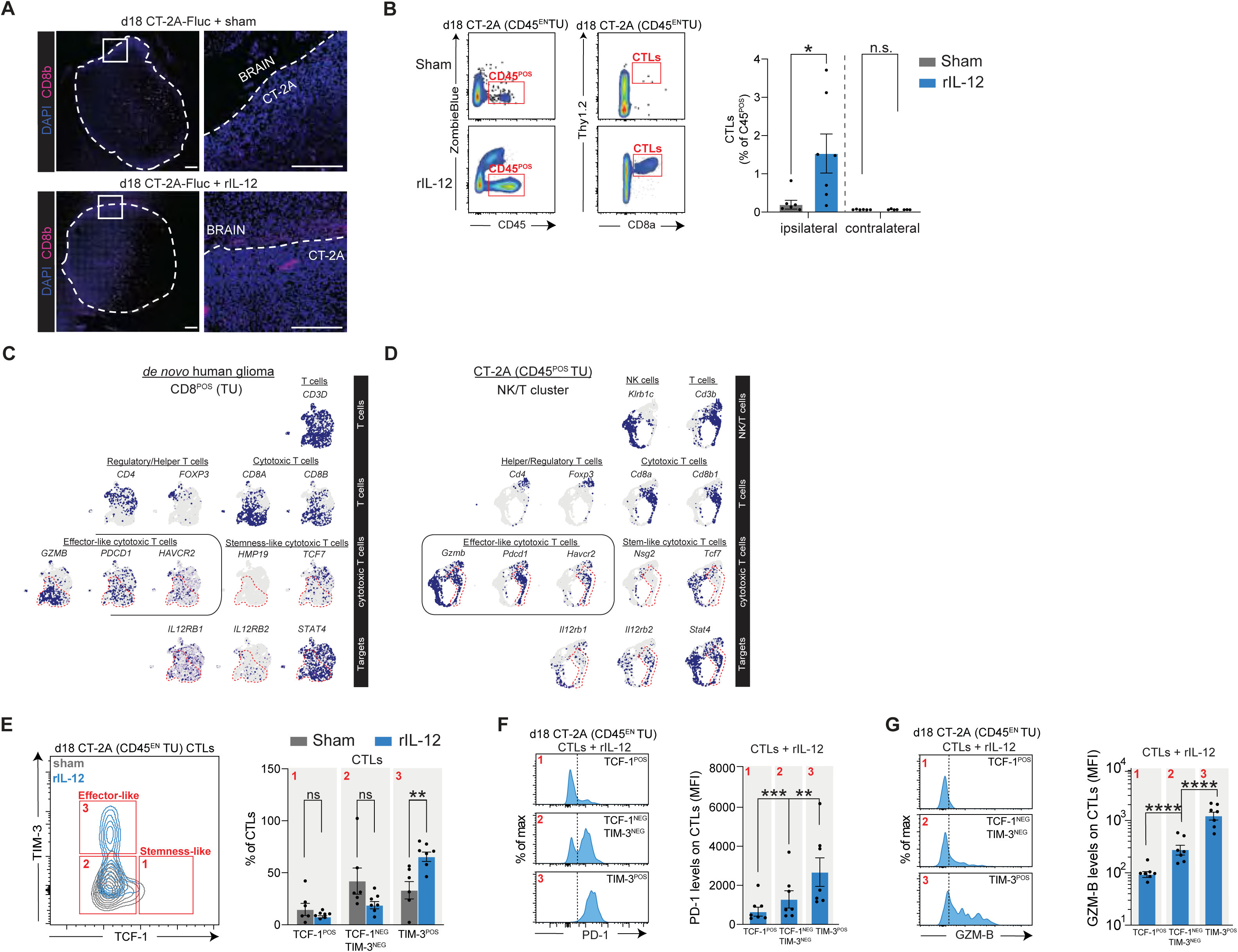
Intratumoral injected rIL-12 increases the number of effector-like CD8^POS^ T cells at the tumor site. (A) *Higher number of CD8^POS^ T cells around rIL-12 treated CT-2A-FLuc tumor.* Immunohistochemistry of CD8b at the tumor border (white dotted line) after 18 days of CT-2A-FLuc implantation comparing sham and rIL-12 treated mice. (Magnification 20x; scale bar 100 µm). (B) *Increase of CD8^POS^ T cells at tumor site with intratumoral rIL-12.* Representative flow cytometry plots of CD45^EN^ CT-2A tumor cells show the gating for live cells based on uptake of the viability dye ZombieBlue staining. Isolation of Thy1.2^POS^ and CD8^POS^ T cells, pre-gated for CD45^POS^CD11b^NEG^ in brains of tumor-bearing mice on day 18 post-tumor (CT-2A CD45^EN^ TU) implantation, comparing rIL-12 treated and sham control (left). The brain tissue was enzymatically digested and enriched for CD45^EN^ tumor cells. The bar graph represents the quantification of CD8^POS^ T cell numbers in ipsilateral and contralateral hemispheres (n=6 mice per group) as a percentage of single cells in rIL-12 or sham-treated mice (right, bar graph). Data represent two independent experiments and are presented as the mean with SEM (error bars). Data were analyzed using unpaired t-test, *p < 0.05. (C) *Differentiating stem- and effector-like CD8^POS^ T cells in de novo human glioma.* scRNAseq analysis classifies T cells based on *CD3D* expression, respectively. Helper and regulatory T cells, expressing *CD4* and *FOXP3* genes, were discriminated from the CD8^POS^ T cells by marked *CD8A/B* expression. In the CD8^POS^ T cell cluster, we observed naïve and stem-like CD8^POS^ T cells, expressing *TCF7* (encoding TCF-1) and *HMP19* genes, that were different from the CD8^POS^ T cells with an effector-like phenotype expressing *HAVCR2* (encoding TIM-3), *PDCD1* (encoding for PD-1), *GZMB* (encoding for cytotoxic granzyme-B). Effector-like T cells are marked with a red dotted line. (D) *Differentiating stem- and effector-like CD8^POS^ T cells in CT-2A murine GB*. scRNAseq analysis of CT-2A (CD45^POS^ TU) cells (Tomaszewski et al., 2022), distinguishes NK cells from T cells based on *Klrb1c* and *Cd3b* expression, respectively. Then, helper and regulatory T cells, expressing *Cd4* and *Foxp3* genes, were discriminated from the CD8^POS^ T cells by marked *Cd8a/b* expression. In the CD8^POS^ T cell cluster, we observed naïve and stem-like CD8^POS^ T cells, expressing *Tcf7* (encoding TCF-1) and *Nsg2* genes, that were different from the CD8^POS^ T cells with an effector-like phenotype expressing *Havcr2* (encoding TIM-3), *Pdcd1* (encoding for PD-1), *Gzmb* (encoding for cytotoxic granzyme-B) and *Tnfrsf9* (encoding for 4-1BB). Effector-like T cells are marked with a red dotted line. (E) *Increased CD8^POS^ T cell differentiation upon rIL-12 treatment of GB*. Overlaid contour plots of TCF-1 expression against TIM-3^POS^ comparing rIL-12 (blue) and sham (grey) of CD8^POS^ T cells on day 18 post-tumor (CT-2A CD45^EN^ TU) implantation (left). Quantification of the percentage of CD8^POS^ T cells comparing TCF-1^POS^ (panel 1 – effector-like), TCF-1^NEG^TIM-3^NEG^ (panel 2) and TIM-3^POS^ (panel 3 – stemness-like) showed a 32.2% increase of TIM-3^POS^ after rIL-12 (65.5%) treatment compared to sham (33.2%) (n=6-7 mice per condition, right bar graph). Data represent two independent experiments and are presented as the mean with SEM (error bars). Data were analyzed using two way ANOVA **p < 0.01, not significant (n.s.). (F) *PD-1 differentiation marker is highly expressed in effector-like CD8^POS^ T cells.* Percent of maximum PD-1 expression within TCF-1^POS^ (panel 1), TCF-1^NEG^TIM-3^NEG^ (panel 2) and TIM-3^POS^ (panel 3) populations on day 18 post-tumor (CT-2A CD45^EN^ TU) implantation, of CD8^POS^ T cells when treated with rIL-12 (left). Quantification of flow cytometry PD-1^POS^ after rIL-12 treatment within TCF-1^POS^ (panel 1 – MFI 2679), TCF-1^NEG^TIM-3^NEG^ (panel 2 – MFI 1280) and TIM-3^POS^ (panel 3 – MFI 652) populations. TIM-3^POS^ cells express significantly more PD-1 compared to TCF-1^POS^ and TIM-3^NEG^TCF-1^NEG^ cells. In contrast, TIM-3^NEG^TCF-1^NEG^ CD8^POS^ T cells express significantly more PD-1^POS^cells, compared to TCF-1^POS^ ones (right). The black dotted line represents the FMO. Data represent two independent experiments and are presented as the mean with SEM (error bars). Data were analyzed using two way ANOVA, **p < 0.01, ***p < 0.001. (G) *Cytotoxic GZM-B is highly expressed in effector-like CD8^POS^ T cells. Percent* of maximum GZM-B expression within TCF-1^POS^ (panel 1), TCF-1^NEG^TIM-3^NEG^ (panel 2) and TIM-3^POS^ (panel 3) populations on day 18 post-tumor (CT-2A CD45^EN^ TU) implantation, of CD8^POS^ T cells when treated with rIL-12 (left). Quantification of flow cytometry of GZM-B after rIL-12 treatment within TCF-1^POS^ (panel 1 – MFI 1229), TCF-1^NEG^TIM-3^NEG^ (panel 2 – MFI 277.3) and TIM-3^POS^ (panel 3 – MFI 93.5) populations (right). (n=5-6 mice per group). The black dotted line represents the FMO. Data represent two independent experiments and are presented as the mean with SEM (error bars). Data were analyzed using two way ANOVA ****p < 0.0001.

CD8^POS^ T cell activity is important for accumulation and proliferation of local CD8^POS^ T cells and is necessary for tumor regression. However, the immunosuppressive nature of GB reduces CD8^POS^ T cell activity and can drive CD8^POS^ T cells into a hypo- or dysfunctional state (Siddiqui et al., 2019). Countering this tumor-enforced immunosuppressive program has been a major interest in the field as it is crucial for sustained anti-tumor activity by CD8^POS^ T cells (Vella et al., 2023). First, we screened for activity markers in glioma-associated CD8^POS^ T cells using scRNAseq datasets derived from human glioma tumors (both *de novo* and recurrent) (**Figures 4C and S5-E-F-G**) and murine GB models (CT-2A, GL261 and 005) (**Figures 4D and S5-H-I**). Furthermore, we verified whether these markers changed during an rIL-12-mediated response with flow cytometry. Immune cells such as NK cells and T cells were distinguished based on specific markers such as expression of *KLRB1C* marking NK cells, while T cells are marked by *CD3D/E* expression. Helper and regulatory T cells were identified through their expression of the *CD4* and *FOXP3* genes, whereas CD8^POS^ T cells were marked by the expression of *CD8A*/*B*. The CD8^POS^ T cell population has distinct subtypes such as the stem-like memory CD8^POS^ T cells, which have self-renewing capabilities, and the effector-like CD8^POS^ T cells, which carry out cytotoxic functions targeting cells presenting tumor antigens (Koh et al., 2023). We observed the expression of signature genes *TCF7* (encoding TCF-1) and *HMP19* (encoding NSG2) in naïve and stem-like CD8^POS^ T cells. In contrast, effector-like CD8^POS^ T cells express genes such as *HAVCR2* (encoding TIM-3), *PDCD1* (encoding PD-1), and *GZMB* (encoding granzyme-B), which are markers of their cytotoxic function and inhibitory state (Ando and Araki, 2022; Hudson et al., 2019; Lak et al., 2022; Utzschneider et al., 2016). In addition, we observed co-expression of *IL12RB1*, *IL12RB2,* and *STAT4,* markers of interest in effector-like CD8^POS^ T cells (**Figures 4C and S5-E**). The same states and expression patterns of markers were found in the murine models (**Figures 4D and S5-H-I**).

An initial screening was performed on isolated CD45^EN^ TU cells to assess whether rIL-12 affects CD8^POS^ T cell activity. This was explored with the markers *Pdcd1* (encoding for PD-1) (Honda et al., 2014), *Gzmb* (encoding for granzyme-B) (Best et al., 2013), and *Cd101* (encoding for CD101) (Hudson et al., 2019), which are signatures for CD8^POS^ T cells, tumor antigen reactivity, cytotoxicity, and differentiation, respectively (**Figure S5-J**). Upon rIL-12 treatment the markers PD-1, GZM-B, and CD101 increased by 5.60-, 3.0- and 2.9-fold, respectively compared to sham-treated tumors (Watowich et al., 2023; Zhou et al., 2020) (**Figure S5-K**). We then performed an in-depth characterization of CD8^POS^ T cell states during rIL-12 treatment with a multiplex flow cytometric analysis of CD45^EN^ TU cells. Eighteen days post-tumor implantation (equivalent to eight days post-rIL-12/sham treatment), 32.3% ± 9.125 (mean difference ± SEM) more TCF-1^NEG^ TIM-3^POS^ CD8^POS^T cells were observed, as compared to the sham-treated GB mice, indicating a transition of CD8^POS^ T cells from a stem-like to an effector-like state (**Figure 4E**). A prerequisite for effector-like CD8^POS^ T cells is that they engage with a tumor antigen-(cross)presenting cell, which provides the necessary signals for them to acquire cytotoxic potential. PD-1 is transiently upregulated on CD8^POS^ T cells upon their interaction with a cross-presented antigen via their T cell receptor. Upon rIL-12 treatment, the TCF-1^NEG^TIM-3^POS^ CD8^POS^ T cells expressed the highest levels of PD-1 (4-fold higher compared to TCF-1^POS^TIM-3^NEG^ and 2-fold higher compared to TCF-1^NEG^TIM-3^NEG^), indicating that this CD8^POS^ T cell state had experienced prolonged or repeated engagement with their T cell receptor compared to the other states (Honda et al., 2014) (**Figures 4F and S5-L**). Next, we checked the tumor-killing potential of the TCF-1^NEG^TIM-3^POS^ CD8^POS^ T cells by determining their cytotoxic GZM-B levels. GZM-B levels gradually increased along the TCF-1-to-TIM-3 differentiation axis upon rIL-12 treatment, accentuating cytotoxic activity in TCF-1^NEG^TIM-3^POS^ CD8^POS^ T cells (GZM-B MFI levels were 13-fold higher compared to TCF-1^POS^TIM-3^NEG^, and 4-fold higher compared to TCF-1^NEG^TIM-3^NEG^) (**Figure 4G**).

In sum, the above data demonstrates that i.t. rIL-12-administration results in an increased number of CD8^POS^ T cells at the tumor site that progress toward an effector-like state.

### Effector-like CD8^POS^ T cells at the GB border sustain 41BB expression post-rIL-12 treatment

Effector-like CD8^POS^ T cells require various signals to become activated and functional, including antigen recognition (signal 1), co-stimulatory signals (signal 2), and cytokine signaling (signal 3) (Joshi and Kaech, 2008; Prokhnevska et al., 2023). We have shown that tumor-associated CD8^POS^ T cells were susceptible to MHC-I-mediated antigen cross-presentation (signal 1) and rIL-12-based cytokine stimulation (signal 2), but we did not verify co-stimulation (signal 3) (**Figure 3A**). Previously, we suggested that rare *Il12b*^POS^ DCs could provide these signals in a GB tumor, including co-stimulatory signals, and that these functions are increased during rIL-12 treatment (**Figures 3B and I**). Here, we verified whether the rIL-12-stimulated CD8^POS^ T cell states (e.g., effector-like CD8^POS^ T cells) have the machinery to bind to co-stimulatory factors of rIL-12-activated DCs. The scRNAseq analysis (*Tnfrsf9* in **Figures 3C and S4-E-G**) on *Il12b*^POS^ DCs and CD8^POS^ T cells (Vinay and Kwon, 2014; Watts, 2005) suggested similar expression of IL-12R and o-stimulatory receptor 4-1BB (Goodwin et al., 1993). This suggests that stimulation of the 4-1BB ligand (4-1BBL) provided by intratumoral *Il12b*^POS^ DCs at the tumor site (*Tnfsf9* in **Figures 3B and S4-F**), could potentially act on both CD8^POS^ T cells and DCs, thereby strengthening the anti-tumor response. Because 4-1BBL can bind to, and activate, 4-1BB on both cell types, similar to rIL-12 binding to IL12R on CD8^POS^ T cells and DCs (Il12rb2 in Figures 3C and S4-E-G), it suggests a potential dual role in enhancing immune responses. This pattern of dual-like activity was not observed with other co-stimulatory molecules examined, such as CD80, which exhibited more limited or cell-type-specific effects.

4-1BB is less enhanced in GB tumors than in healthy brain tissue (**Figure S6-A**). The lack of a significant increase of 4-1BB in *de novo* glioma tumors is a potential reason why no survival benefit was observed between patients with low or high expression levels (**Figure 5A**). The findings for 4-1BB are also paralleled by those for its ligand, 4-1BBL (**Figures S6-A-B**). Interestingly, with our murine GB model, we observed that 4-1BB^POS^ CD8^POS^ T cells were mainly detected at the tumor site (**Figures 5B-C and S6-C**) and not in the spleen (**Figure S6-D**). We confirmed that effector-like CD8^POS^ T cells at the tumor site can express *TNFRSF9* (encoding for 4-1BB) (**Figure 5D**). Our findings in the *de novo* human glioma datasets were validated in recurrent human glioma (**Figure S5-E**) and in CT-2A, GL261, 005 murine models (**Figures 5D and S5-H-I**). We further characterized the 4-1BB levels in the CD8^POS^ T cell states upon rIL-12 treatment with flow cytometry. TCF-1^NEG^TIM-3^POS^ CD8^POS^ T cells or effector-like T cells expressed ∼5-fold more 41BB compared to TCF-1^POS^TIM-3^NEG^ or TCF-1^NEG^TIM-3^NEG^ (**Figures 5E**).

**Figure 5.**
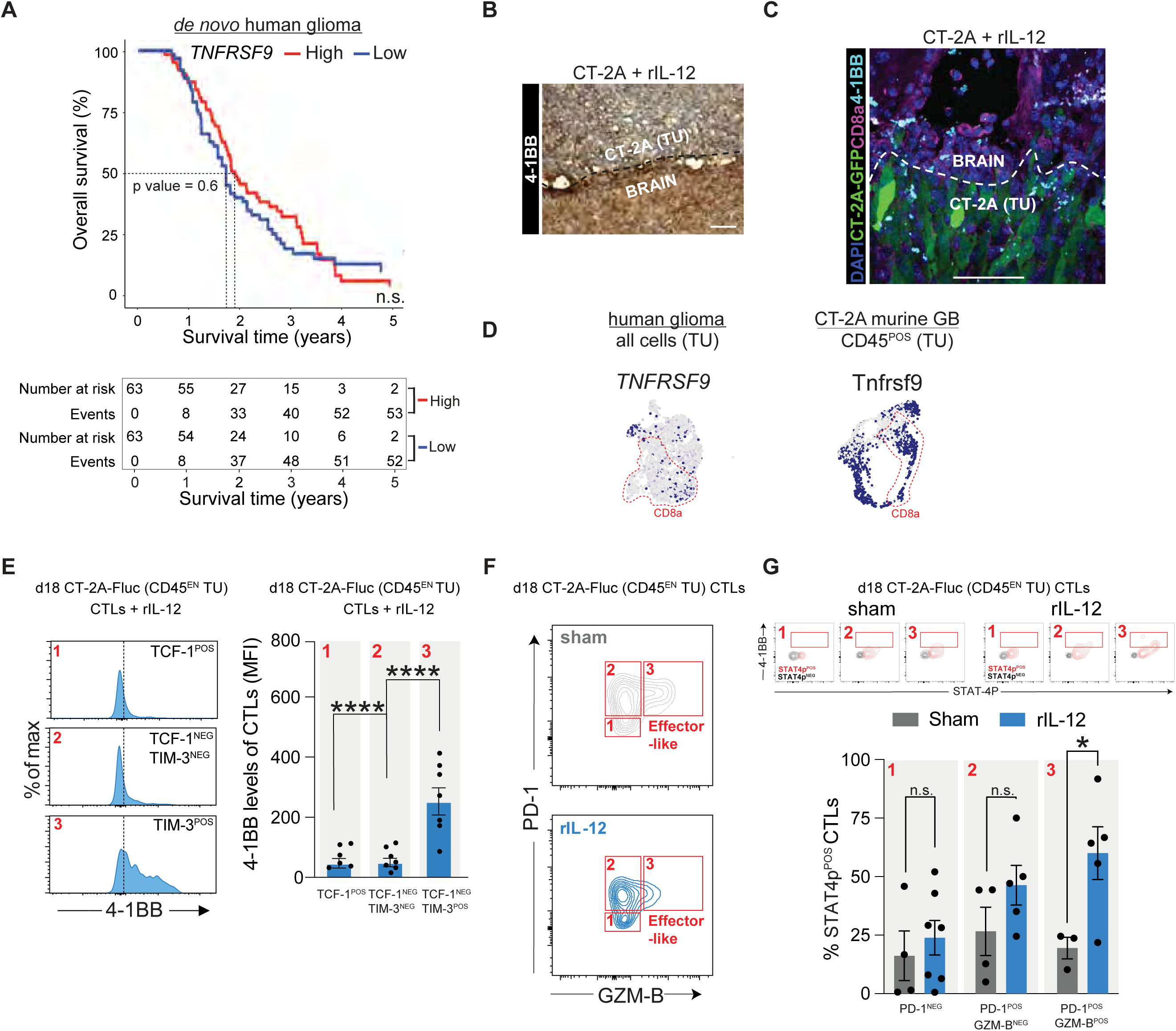
In line with rIL-12 activated DCs, effector-like CD8^POS^ T cells at the GB border maintain 41BB expression post-rIL-12 treatment. (A) *Survival probability of de novo human glioma differentiating high and low TNFRSF9.* Kaplan-Meier survival curves showing the survival outcomes over a period of 5 years of 63 glioma patients per group with high (red) or low (blue) levels of *TNFRSF9*, each group had a median of ∼2 years (Miller et al., 2023). No differences were observed between groups. Log-rank (Mantel-Cox) test, p-value = 0.6, not significant (n.s). (B) *Immunohistochemistry shows 4-1BB expression at the tumor border.* The CT-2A-bearing brain was stained by HRP for 4-1BB. (Magnification 20x; scale bar 100 µm). (C) *CD8^POS^ T cells expressing 4-1BB are recruited at the tumor upon IL-12 treatment*. Immunofluorescence shows that 4-1BB (cyan) was expressed in CD8^POS^ T cells (pink) post-rIL-12 treatment at the CT-2A-GFP (green) tumor border (white dotted line). (20x magnification, scale bar = 50 µm). (D) *4-1BB expression*. scRNAseq analysis displaying *TNFRSF9* expression in all cells (TU) in human glioma and *tnfrsf9* expression in CD45^POS^ (TU) cells in CT-2A murine GB (Tomaszewski et al., 2022). The red dotted line represents the CD8a population. (E) *4-1BB^POS^ cells are effector-like CD8 ^POS^ T cells.* Percent of maximum 4-1BB expression (left histogram plot) within TCF-1^POS^ (panel 1), TCF-1^NEG^TIM-3^NEG^ (panel 2) and TIM-3^POS^ (panel 3) populations on day 18 post-tumor (CT-2A CD45^EN^ TU) implantation of CD8^POS^ T cells when treated with rIL-12. Quantification by flow cytometry of 4-1BB (right bar graph) after rIL-12 treatment within TCF-1^POS^ (panel 1 – MFI: 251.9), TCF-1^NEG^TIM-3^NEG^ (panel 2 – MFI 48.4) and TIM-3^POS^ (panel 3 – 46.3) populations. (n=4-7 mice per group). (The black dotted line represents the FMO). Data represent two independent experiments and are presented as the mean with SEM (error bars). Data were analyzed using two way ANOVA, ****p < 0.0001. (F) *4-1BB^POS^ cells are more present in STAT4 phosphorylated CD8^POS^ T cells.* Counter plots of GZM-B expression against PD-1 within PD-1^POS^ (panel 1), PD-1^NEG^GZM-B^NEG^ (panel 2) and GZM-B^POS^, representing effector-like cells (panel 3) populations on day 18 post-tumor (CT-2A CD45^EN^ TU) implantation of CD8^POS^ T cells when treated with sham (top) or rIL-12 (bottom). (G) Cells were pre-gated for 4-1BB versus pSTAT4 comparing sham and rIL-12 treated conditions (top). Quantification of flow cytometry as shown in bar graphs of percentage of 4-1BB^POS^ cells post-rIL-12 treatment showing STAT4 phosphorylation of CD8^POS^ T cells within PD-1^NEG^ (panel 1), PD-1^NEG^GZM-B^POS^ (panel 2) and PD-1^POS^GZM-B^POS^ (panel 3) populations (bottom). (n=4 mice per group). Data represent two independent experiments and are presented as the mean with SEM (error bars). Data were analyzed using two way ANOVA, *p < 0.05.

In analogy to the 4-1BB^POS^ (representing *Il12b*^POS^) DC analysis, we tested whether the 4-1BB^POS^ CD8^POS^ T cells are rIL-12 responsive by analyzing STAT4-phosphorlyation. In effector-like CD8^POS^ T cells, as identified by PD-1 and GZM-B markers (**Figure 5F**), we detected a 4-1BB^POS^ subpopulation that was enriched for STAT4 phosphorylation (pSTAT4^POS^ cells) (**Figure 5G - TOP**). rIL-12 treatment led to a 41.4% ± 1.131 (mean difference ± SEM) increase in pSTAT4^POS^ in this 4-1BB^POS^ effector-like CD8^POS^ T cell subpopulation compared to the sham condition (**Figure 5G – BOTTOM**). In other CD8^POS^ T cell states (either PD-1^NEG^GZM-B^POS^ or PD-1^NEG^ CD8^POS^ T cells), low numbers of CD8^POS^ T cells co-expressing 4-1BB were observed. Therefore, STAT4 phosphorylation did not increase compared to sham control (**Figure 5G**).

Taken together, our findings demonstrate that effector-like CD8^POS^ T cells at the tumor site during rIL-12 treatment express the 4-1BB receptor, making them susceptible to activation by 4-1BBL.

### Anti-tumor immunity triggered by the combined rIL-12 and 4-1BBL immune stimuli enhanced CD8^POS^ T cell-mediated survival in GB-bearing mice

Clinical trials with therapeutic i.t. IL-12 expression suggested that the activity of CD8^POS^ T cells was rapidly diminished due to the PD-L1-rich GB environment (Chiocca et al., 2019). Rather than inhibiting immunosuppression with ICI, which failed to enhance IL-12-mediated survival in patients (Cloughesy et al., 2019; Filley et al., 2017; Kurz et al., 2018; Nayak et al., 2021; Omuro et al., 2018; Reardon et al., 2020; Schalper et al., 2019), here we aim to boost survival by providing a co-stimulatory molecule to support tumor-associated CD8^POS^ T cells directly, or indirectly through DCs. The co-stimulatory factor *TNFSF9* (encoding 4-1BBL) was selected because it was less induced in GB tissue, to the extent that patients with detectable *TNFSF9* (4-1BBL) expression levels did not gain in survival (**Figures S6-A-B**).

First, we determined whether our mouse model would be able to mimic the high levels of PD-L1 at the GB site during rIL-12 treatment. In rIL-12 and sham-treated tumors, we confirmed increased levels of PD-L1 in our murine GB tumors. This marker was mainly found on tumor cells (represented predominantly in CD45^DEPR^ TU cell fraction), where we observed ∼125-fold higher expression of *Cd274* (encoding for PD-L1) compared to the immune cell compartment (represented by both the CD11b^EN^ and CD45^EN^ TU cells) (**Figure 6A**). We next addressed whether rIL-12-activated tumor-associated 4-1BB^POS^ immune cells (CD8^POS^ T cells and DCs) could be stimulated to further enhance anti-tumor regression in PD-L1-rich GB tumors. To explore this, we designed a lentiviral vector (LVV) encoding murine 4-1BBL (m4-1BBL) (**Figure 6B**). LVV-encoded m4-1BBL could be distinguished from endogenous m4-1BBL through an N-terminal 3xFLAG-tag, which in the recombinant protein localizes to the intracellular-facing side, ensuring it does not interfere with 4-1BB binding. A T2A protease cleavage site separating mCherry fluorescent reporter transgene was included to confirm the transduction of our tumor cells. An inactive mimic LVV (LVV null), encoding mCherry but lacking the 3xFLAG-tag and m4-1BBL, was designed as a control. Following LVV m4-1BBL transduction of CT-2A-FLuc and 005-FLuc cells, mCherry^POS^ cells were sorted via FACS and confirmed to overexpress m4-1BBL compared to LVV null cells. Our qRT-PCR analysis indicated a 30- and 38-fold increase of *Tnfsf9* (encoding m4-1BBL) in both CT-2A and 005 GB cell lines, respectively, compared to the controls (LVV null and non-transduced) (**Figure 6C**). A western blot analysis was performed using 3xFLAG-tag detection to confirm that the transgene was expressed as a full-length protein with the expected size of 37.5 kDa (**Figure 6D**). We validated the uniform recombinant protein expression in the transduced cells through colocalization of mCherry fluorescence with anti-3xFLAG-tag and anti-m4-1BBL staining (**Figure 6E**). Importantly, although *in vitro* all transduced cells expressed the construct, m4-1BBL and 3xFLAG-tag expression was not uniformly observed throughout the entire CT-2A-FLuc-m4-1BBL tumor on day 18 post-tumor implantation. We hypothesize that this might be due to transgene instability in tumor cells, promoter silencing, or potential overgrowth by small numbers of non-expressing 4-1BBL GB cells, rather than an effect of rIL-12 treatment. The latter was tested *in vitro* using a cell viability assay on CT-2A-FLuc-m4-1BBL and CT-2A-FLuc-null cell lines exposed to either rIL-12 or sham and did not observe any significant differences (**Figure S7-A**).

**Figure 6.**
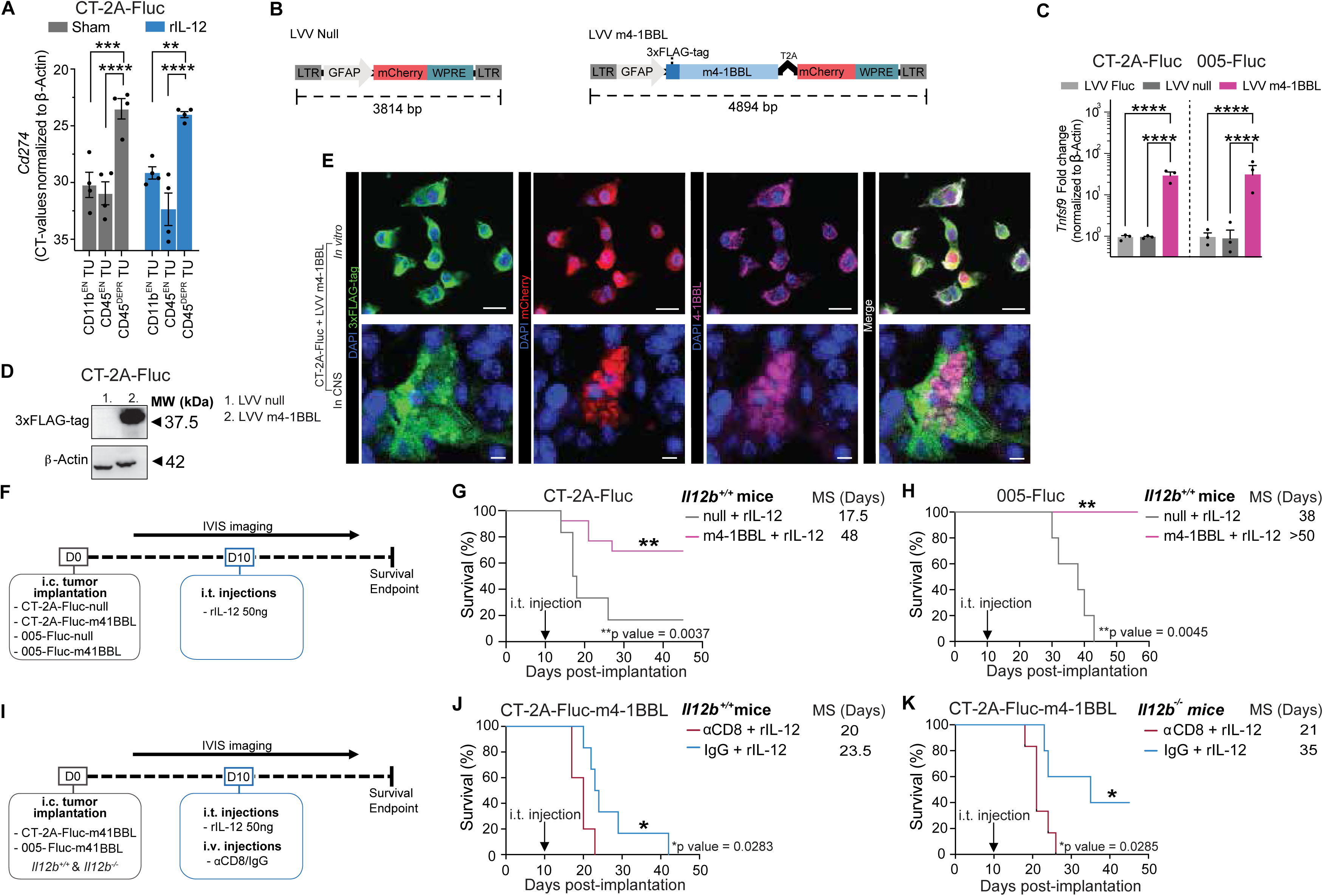
Anti-tumor immunity activated by combined rIL-12 and m4-1BBL immune stimuli increased the survival of GB-bearing mice. (A) *GB tumors in mice express high levels of PD-L1*. CD11b^EN^, CD45^EN^ and CD45^DEPR^ TU populations were isolated from CT-2A-FLuc tumor-bearing hemisphere of brains using CD11b beads and CD45 beads. *Cd274* (the PD-L1 gene) was expressed at significantly higher levels in CD45^DEPR^ TU cells. No differences were observed between sham (grey) and rIL12 treatment (blue). Gene expression levels were normalized to β-actin. (n=4 mice per group). Data represent four independent experiments and are presented as the mean with ± SEM (error bars). Data were analyzed using one way ANOVA, **p < 0.01, ***p< 0.001, ****p < 0.0001. (B) *Lentiviral m4-1BBL constructs.* Schematic display of m4-1BBL lentiviral vectors (LVV); LVV null, a GFAP promotor followed mCherry and WPRE; LVV m41BBL containing mCherry labelled m4-1BBL and 3xFLAG-tag driven by a GFAP promotor. (C) *m4-1BBL expression in GB mouse cells.* CT-2A-FLuc and 005-FLuc cells transduced with the LVV m4-1BBL showed significant enhanced gene expression levels of *TNfsf9* (encoding for 4-1BBL) compared to cells transduced with LVV null and non-transduced cells, normalized to β-Actin. Data represent three independent experiments and are presented as the mean with ± SEM (error bars). Data were analyzed using one way ANOVA, ****p < 0.0001. (D) *m4-1BBL protein expression in GB mouse cells*. 3xFLAG-tag protein levels (37.5 kDa) were only present in CT-2A-FLuc cells transduced with the LVV m4-1BBL compared to cells transduced with LVV null, normalized to β-Actin. 3xFLAG-tag detection enabled detection of transgene m4-1BBL and not endogenous 4-1BBL. (E) *Homogenous m4-1BBL expression in transduced GB mouse cell line and tumor-bearing CNS*. Immunofluorescence images of m4-1BBL overexpressing CT-2A cells post-LVV m4-1BBL transduction in culture stained for DAPI (blue) 3xFLAG-tag (green), mCherry (red) and 4-1BBL (pink) with a merged image (scale bar = 50 µm). Tumor-bearing mouse brains confirmed transgene expression (mCherry-positive cells) co-localized with 3xFLAG-tag and m4-1BBL. (40x magnification, scale bar = 50 µm). (F) *Experimental outline to test local expression of m4-1BBL and rIL-12 treatment*. The *in vivo* approach is schematically displayed; CT-2A-FLuc-null; CT-2A-FLuc-m4-1BBL; 005-FLuc-null; 005-FLuc-m4-1BB cells were implanted i.c., mice were treated i.t. with rIL-12 or sham (PBS or Fc-control) 10 days after tumor implantation. (G) *Survival benefit of local m4-1BBL expression in CT-2-FLuc-bearing mice post-rIL-12 treatment*. Kaplan-Meier curves showing survival outcomes following treatment of CT-2A-FLuc-control with rIL-12 (solid grey) or CT-2A-FLuc-m4-1BBL treated with rIL-12 (solid pink) (n= 4-5 mice per group). Mice injected with CT-2A-FLuc-m4-1BBL tumor cells treated with rIL-12 (50 ng) had a median survival of 48 days (p-value = 0.0037) compared to mice implanted with tumor cells lacking m4-1BBL, median survival of 17.5 days. Data represent at least two independent experiments and are presented as the mean with ± SEM (error bars). Data were analyzed using Log rank (Mantel-Cox) test **p < 0.01. (H) *Survival benefit of local m4-1BBL expression in 005-FLuc-bearing mice post-rIL-12 treatment*. Kaplan-Meier curves showing survival outcomes following treatment of 005-FLuc-control with rIL-12 (solid grey) or 005-FLuc-m4-1BBL treated with rIL-12 (solid pink) (n= 4-5 mice per group). Mice injected with 005-FLuc-m4-1BBL tumor cells treated with rIL-12 (50 ng) had a 100% survival (p-value = 0.0045) compared to mice implanted with tumor cells lacking m4-1BBL, median survival of 38 days. Data represent at least two independent experiments and are presented as the mean with ± SEM (error bars). Data were analyzed using Log rank (Mantel-Cox) test **p < 0.01. (I) *Experimental outline to test CD8 T cell dependency of m4-1BBL and rIL-12 combination treatment.* Schematic display shows intravenous (i.v.) injection with or without CD8 T cell depletion (αCD8 or IgG control, respectively) on day 9 and day 10 (50 µg and 100 µg on day 9 and 10, respectively) post-tumor (CT-2A-FLuc-m4-1BBL, 005-FLuc-m4-1BBL) implantation. Mice were injected i.t. with rIL-12 50 ng on day 10. (J) *GB mouse survival benefit of m4-1BBL and rIL-12 combination treatment is CD8 T cell dependent.* Kaplan-Meier curves of a total of 12 *Il12^+/+^* mice showing survival outcomes of CT-2A-FLuc-m4-1BBL tumor-bearing mice all treated with rIL-12 i.t., after treatment with αCD8 (red) or IgG control (blue). Mice (n=5-6 mice per group) treated with IgG control had a median survival of 23.5 days (p-value = 0.0283), compared to 20 days for mice treated with αCD8. Data represent two independent experiments and are presented as the mean with ± SEM (error bars). Data were analyzed using Log-rank (Mantel-Cox) test *p < 0.05. (K) *GB mouse survival benefit due to CD8 T cell recruitment of m4-1BBL, and rIL-12 combination treatment is not dependent on endogenous IL-12.* Kaplan-Meier curves of a total of 12 *Il12^−/−^* mice showing survival outcomes of CT-2A-FLuc-m4-1BBL tumor-bearing mice all treated with rIL-12, after treatment with αCD8 (red) or IgG control (blue). Mice (n=5-6 mice per group) treated with IgG control had a median survival of 35 days (p-value = 0.0285), compared to 21 days for mice treated with αCD8. Data represent at least two independent experiments and are presented as the mean with ± SEM (error bars). Data were analyzed using Log-rank (Mantel-Cox) test *p < 0.05.

Next, we assessed the survival rates of GB-bearing mice in response to rIL-12 and m4-1BBL was assessed (**Figure 6F**). Mice brains were engrafted with CT-2A-FLuc-m4-1BBL or CT-2A-FLuc-null GB cells and i.t. treated with rIL-12 or sham on day 10 post-implantation. Mice implanted with CT-2A-FLuc-m4-1BBL tumors showed a prolonged median survival of 30.5 days and 70% of the mice survived >50 days, compared to those implanted with CT-2A-FLuc-null cells (**Figures 6G and S7-B-C**). The survival advantages following rIL-12 treatment with the co-stimulatory signal m4-1BBL were confirmed in the 005-GB mouse model, showing that all mice implanted with 005-FLuc-m4-1BBL survived for >50 days compared to mice implanted with 005-FLuc-null cells which had a median survival of 38 days (**Figures 6H**). For the CT-2A model, we also verified the effect of m4-1BBL without rIL-12 treatment. The median survival of CT-2A-FLuc-m4-1BBL implanted mice was extended by 27 days compared to the CT-2A-FLuc-null implanted mice (**dotted lines in Figure S7-B**), suggesting that high levels of i.t. m4-1BBL also can counter the immunosuppressed TME. Mice that survived after initial implantation of CT-2A-FLuc-m4-1BBL or 005-FLuc-m4-1BBL cells were rechallenged with a second tumor (CT-2A-FLuc and 005-FLuc, respectively). Interestingly, only five out of thirteen re-implanted mice developed new tumors in the CT-2A GB model, and 62% did not regrow tumors upon re-implantation, suggesting that protective immunity had developed during the rejection of m4-1BBL-expressing tumors. The 005-FLuc re-implanted mice also did not regrow tumors (**Table 3**).

To evaluate the impact of co-stimulatory 4-1BBL on CD8^POS^ T cells during an rIL-12-mediated anti-tumor response, mice were systemically depleted of CD8^POS^ T cells with anti-CD8 mAb to avoid accumulation at the CT-2A-FLuc-m4-1BBL tumor site (**Figure 6I**). Improved survival of 3.5 days was observed for non-depleted (IgG control) compared to the T cell-depleted (anti-CD8) CT-2A-FLuc-m4-1BBL implanted mice treated with i.t. rIL-12 (**Figures 6J and S7-D**). Next, we tested if augmenting co-stimulation with m4-1BBL in *Il12b^−/−^*mice induced tumor regression independently of endogenous IL-12p70. Similar to *Il12b^+/+^* mice, rIL-12 treatment improved median survival compared to the sham treatment in *Il12b^−/−^* mice implanted with CT-2A-FLuc-m4-1BBL cells (**Figure S7-E-F**), and depletion of CD8^POS^ T cells reversed the increase of survival (**Figures 6K and S7-G**). Interestingly, all *Il12b^−/−^* sham-treated mice died despite m4-1BBL overexpression, which was not the case for the *Il12b^+/+^*sham-treated mice (**Figure S7-B and S7-E**). This indicates that it is necessary for m4-1BBL to act on functional Il12b^POS^ DCs at the tumor site to mediate its effects, as tumor regression and improved survival were dependent on their presence. This was not the case in rIL-12-treated *Il12b^−/−^* mice, which had similar survival rates as rIL-12-treated *Il12b^+/+^* mice, suggesting that in this setting 4-1BBL mainly acts on CD8^POS^ T cells and its anti-tumor effects is less dependent on its activity on functional *Il12b*^POS^ DCs (**Figure S7-H**).

Taken together, these findings indicate that increased local expression of the co-stimulatory factor 4-1BBL has the potential to support the cytotoxic activity of CD8^POS^ T cells associated with GB during rIL-12 treatment. Augmenting local 4-1BBL expression supplements the co-stimulation that is normally provided by rare tumor-associated DCs.

### rIL-12 administration combined with AAVF vector-mediated delivery of m4-1BBL in GFAP^POS^ cells increases survival in GB-bearing mice

We hypothesized whether a therapeutically relevant AAV vector-based gene therapy approach could effectively deliver 4-1BBL to enhance IL-12-mediated CD8^POS^ T cell activity. We focused on targeting GFAP^POS^ cells, as GFAP marker is strongly present in GB tumors in patients (Zhang et al., 2023). Elevated expression of GFAP was observed in malignant cells, predominantly in astrocyte-like cells (Couturier et al., 2020; Neftel et al., 2019) (**Figure S8-A-B**). Therefore, we compared three GB mouse models (CT-2A, 005, GL261) with varying levels of GFAP expression levels in both the tumor and the peritumoral region (**Figures 7A and S8-C**). Although both our tumor cell lines and primary astrocytes were confirmed to express *Gfap* (**Figures 7B and S8-D**), we expect that non-dividing or slowly dividing GFAP^POS^ cells in the peritumoral regions are most important for our strategy, as there they co-localize with tumor-associated DCs and CD8^POS^ T cells, in addition to having the highest potential for extended AAV-vector transgene expression. To confirm that non-tumor GFAP^POS^ cells could be targeted by a GFAP-driven transgene, AAVF-*GFAP*-GFP was i.c. injected in mice (**Figure S8-E**).

**Figure 7.**
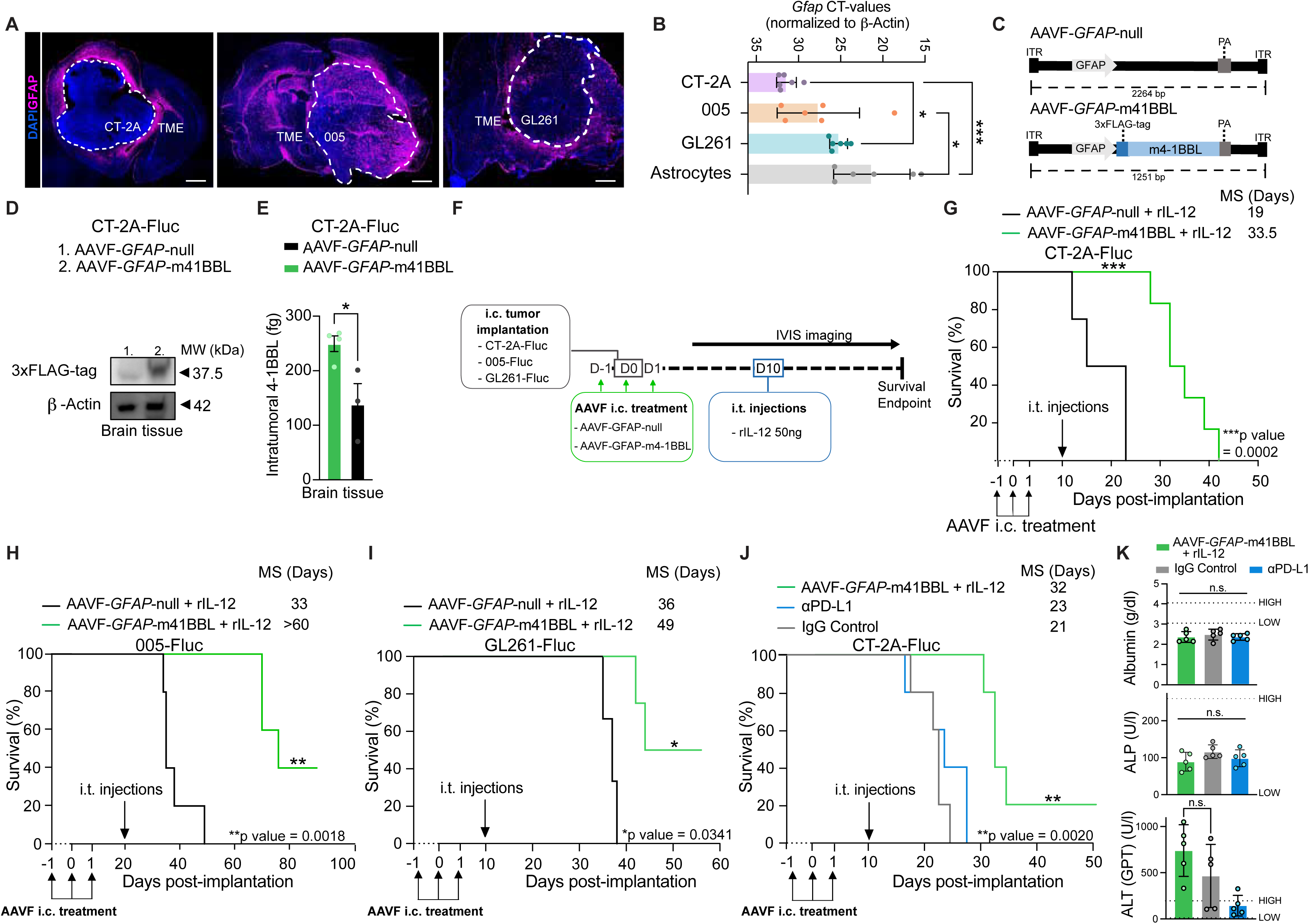
rIL-12 administration combined with AAVF-mediated delivery of 4-1BBL in GFAP^POS^ cells as a therapeutic intervention. (A) *GFAP^POS^ cell expression profile in the TME is dependent on the type of mouse GB cell line*. Immunofluorescence showing GFAP^POS^ astrocytes at the tumor border (whited dashed line) of CT-2A (left), 005 (middle) and GL261(right) tumors 18 days post-implantation. With 005 and GL261 tumors the GFAP^POS^ cells were retrieved in the brain tumor cell mass. (4x magnification, scale bar = 5 µm). (B) *Endogenous GFAP expression in mouse GB cell lines and astrocytes.* qRT-PCR analysis measuring *Gfap* expression levels for GB cell lines, CT-2A and 005, GL261 and primary brain-derived astrocytes (n=6/condition). Data represent three independent experiments and are presented as the mean with ± SEM (error bars). Data were analyzed using one way ANOVA, *p<0.05, ***p<0.001. (C) *AAVF vector constructs to deliver m4-1BBL to tumor site.* Schematic representation of the 4-1BBL AAVF (AAVF-*GFAP*-m4-1BBL) and control AAVF (AAVF-*GFAP*-null) constructs. The m4-1BBL-3xFLAG-tag and m4-1BBL are under a GFAP promotor with a poly(A) signal after the coding sequence. In the control AAVF-*GFAP*-null, the GFAP promoter and poly(A) signal were connected without the presence of intervening sequences. (D) *m4-1BBL protein expression at tumor site.* 3xFLAG-tag protein was only detected in brains injected with AAVF-*GFAP*-m4-1BBL-3xFLAG-tag (37.5 kDa) as normalized to β-Actin (42 kDa) by western blot analysis. No fragmentation of the transgenic product was observed. (E) *4-1BBL protein levels in GB-containing brain hemisphere.* 4-1BBL levels were determined in femtogram (fg) using Luminex in GB-bearing (CT-2A-FLuc). Data represents three independent experiments and are presented as the mean with SEM (error bars). Unpaired t-test, *p < 0.05. (F) *Graphic depiction of the treatment scheme of AAVF-GFAP-m4-1BBL experiments.* m4-1BBL-coding or AAVF-null vectors were injected i.t. at three time points; one day prior to tumor implantation (CT-2A-FLuc, 005-FLuc, GL261-FLuc), at the time of tumor implantation, and 1-day post-tumor implantation. rIL-12 was injected i.t. on day 10 post-implantation at the tumor site, and mice were followed by IVIS every 3-4 days. (G) *Survival benefit with AAV-mediated delivery of m4-1BBL in rIL-12 treated CT-2A-FLuc-bearing mice.* Kaplan-Meier curves displaying the percentage of survival of CT-2A-FLuc-bearing mice (12,500 cells at the time of injection) comparing AAVF-*GFAP*-m4-1BBL (green) and AAVF-*GFAP*-null (black) vectors both treated with rIL-12 (n= 4-6 mice per group). AAVF*-GFAP*-m4-1BBL rIL-12 treated had a median survival of 33.5 days (p-value = 0.0002) compared to AAVF-*GFAP*-null with a median survival of 19 days. Data represent at least two independent experiments and are presented as the mean with ± SEM (error bars). Data were analyzed using the Log-rank (Mantel-Cox) test, ***p < 0.001. (H) *Recovery of survival benefit with AAV-mediated delivery of m4-1BBL into delayed rIL-12 treatment of 005-FLuc-bearing mice.* Kaplan-Meier curves displaying the percentage of survival of 005-FLuc-bearing mice (50,000 cells at the time of injection) comparing AAVF-*GFAP*-m4-1BBL (green) and AAVF-*GFAP*-null (black) vectors both treated with rIL-12 on day 20 post-tumor implantation (n=5 mice per group). AAVF-*GFAP*-m4-1BBL rIL-12 treated had a median survival of >60 days (p-value = 0.0018) compared to AAVF-*GFAP*-null with a median survival of 33 days. Data represent at least two independent experiments and are presented as the mean with ± SEM (error bars). Data were analyzed using Lo-rank (Mantel-Cox) test, **p < 0.01. (I) *Survival benefit with AAV-mediated delivery of m4-1BBL in in rIL-12 treated GL261-FLuc-bearing mice.* Kaplan-Meier curves displaying the percentage of survival of GL261-FLuc-bearing mice (50,000 cells at the time of injection) comparing AAVF-*GFAP*-m4-1BBL (green) and AAVF-*GFAP*-null (black) vectors both i.t. treated with rIL-12 on day 10 post-tumor implantation (n=4 mice per group). AAVF-*GFAP*-m4-1BBL rIL-12 treated mice had a median survival of 49 days (p-value = 0.0341) compared to AAVF-*GFAP*-null with a median survival of 36 days. Data represent at least two independent experiments and are presented as the mean with ± SEM (error bars). Data were analyzed using the Log-rank (Mantel-Cox) test, *p<0.05. (J) *The survival advantage of mice treated with AAVF-GFAP-m4-1BBL, and rIL-12 compared to* α*PD-L1 therapy in CT-2A-FLuc-bearing mice.* Kaplan-Meier curves show the percentage of survival of CT-2A-FLuc-bearing mice (50,000 cells at the time of injection). AAVF-*GFAP*-m4-1BBL injected on days −1, 0 and 1 and rIL-12 i.t. treated on day 10 post-tumor implantation (green) with αPD-L1 (blue), and IgG control (grey) (n=5 mice per group). αPD-L1 and IgG control groups were treated i.p. with 200ug in 100ul volume on days 3, 5 and 14, post-tumor implantation. AAVF-*GFAP*-m4-1BBL and rIL-12 treated mice had a median survival of 32 days, significantly improved (p-value = 0.0020) compared to the median survivals of mice treated with αPD-L1 23 days and IgG control 21 days. Data represent one independent experiment and are presented as the mean with ± SEM (error bars). Data were analyzed using the Log-rank (Mantel-Cox) test, **p < 0.01. (K) *Blood pathology toxicity analysis post-therapy.* The blood of mice (n=5 per group) that received AAVF-*GFAP*-m4-1BBL and rIL-12 (green), αPD-L1 (blue), or IgG control (grey) was tested for toxicity markers systemically including Albumin, ALP and ALT, no significance differences were observed. Blood was collected retro-orbitally on day 15 post-tumor implantation. Data represent one independent experiment and are presented as the mean with ± SEM (error bars). Data were analyzed using one way ANOVA, not significant (n.s.).

AAV vector constructs with a *GFAP* promoter, were designed to encode m4-1BBL-3xFLAG-tag (AAVF-*GFAP-*m4-1BBL) and a control, lacking the transgene (AAVF-*GFAP-*null) (**Figure 7C**). These cassettes were packaged into an AAVF capsid, selected for its robust transduction of astrocytes in the peritumoral region (Beharry et al., 2022). To validate full-length recombinant m4-1BBL in i.c. injected AAV vector-treated mice with CT-2A-FLuc tumors, western blot analysis was performed with an anti-FLAG-tag antibody (**Figure 7D**). We confirmed that 37.5 kDa m4-1BBL was expressed in AAVF-*GFAP-*m4-1BBL-treated tumor brain samples and was not present in the AAVF-*GFAP*-null condition. Additionally, the concentration of i.t. m4-1BBL was measured with Luminex and showed 45% increased levels in the tumor hemisphere of mice treated with AAVF-*GFAP-*m4-1BBL compared to AAVF-*GFAP*-null (**Figure 7E**).

AAVF-*GFAP-*m4-1BBL and AAVF-*GFAP*-null were tested in mice i.c. engrafted with three different syngeneic GB cell lines (CT-2A-FLuc, 005-FLuc, and GL261-FLuc) and treated with either rIL-12 or sham (**Figure 7F**). The treatment strategy involved three i.c. injections of AAVF vectors over 3 days, within a timeframe that would not trigger anti-AAVF immunogenicity (Verdera et al., 2020) but still guaranteed sufficient m4-1BBL expression at the tumor site. These GB cell lines have a different growth rates and survival profiles when implanted in mice (**Figures 1E-G-H and S2-D-H-I**). To have comparable tumor size among models, rIL12 treatment was given at around half the expected survival time post-tumor implantation. CT-2A-FLuc and GL261-FLuc were treated on day 10 post-implantation, and 005-FLuc cells were on day 20. The latter grows more slowly and has a diffuse tumor phenotype that is more reflective of human GB. Interestingly, despite equal BLI signals at these time points between the CT-2A and 005 models (day 10 and 20, respectively), delaying the i.t. rIL-12 injection in the 005 model rendered it less susceptible to rIL-12 (**Figure S8-F**). In three murine GB models, AAVF-*GFAP-*m4-1BBL treatment combined with rIL-12 administration improved median survival compared to AAVF-*GFAP-*null combined with rIL-12 (**Figures 7G-H-I and S8-G-H-I**). A 14.5-day survival benefit was observed in the CT-2A-FLuc model. For mice implanted with 005-FLuc and GL261-FLuc, survival rates of 40% up to 80 days and 45% up to 60 days were observed, respectively. The improved survival advantage for GL261 and 005 GB models compared to the CT-2A model could be due to the lower number of activated astrocytes at the CT-2A border and the lower *gfap* expression of the CT-2A cells (**Figures 7A-B**). However, we could confirm that *in vivo* astrocytes were more amenable to our AAV vector-mediated transgene expression than tumor cells when analyzing the tumor border of the treated tumors (**Figure S8-J-K**). We confirmed transgene expression in GFAP^POS^ cells at the tumor border (indicated by the white dotted line) through the detection of the 3xFLAG-tag following AAVF-*GFAP-*m4-1BBL treatment, which was absent in the AAVF-*GFAP-*null condition (**Figures S8-J-K**) despite the observation that *in vitro* exposure of our AAVF-*GFAP-*m4-1BBL/null demonstrated that tumor cells were able to be transduced by our vectors (**Figures S8-L**).

We compared the AAVF-*GFAP*-m4-1BBL and rIL-12 co-therapy to another immunomodulatory approach, anti-PD-L1 treatment (**Figure 7J and S8-M**). Following previously reported regimen (Bowman-Kirigin et al., 2023), 200 μg anti-PD-L1 or IgG control were administered intraperitoneally (i.p.) on days 7 and 14 post-CT-2A-FLuc implantation. AAVF-*GFAP*-m4-1BBL and rIL-12 treatment prolonged the median survival by 9 and 11 days compared to mice treated with either anti-PD-L1 or IgG control, respectively. This represents an improvement of ∼40%, over anti-PD-L1 monotherapy, showing the advantage of this gene therapy approach with rIL-12. Liver markers, including Albumin, ALT, and ALP, showed no significant differences between mice administered with AAVF-*GFAP*-m41BBL, IgG control, and anti-PD-L1, indicating that our gene therapy-induced minimal detectable systemic toxicity (**Figure 7K**).

These results demonstrate that combining rIL-12 with gene therapy delivering the co-stimulatory factor m4-1BBL prolongs the survival rate of GB-bearing mice more effectively than anti-PD-L1 therapy.

## DISCUSSION

Tumor-reactive CD8^POS^ T cells are both rare and often dysfunctional in glioblastoma (GB) tumors (Friebel et al., 2020; Maddison et al., 2021; Nickl et al., 2023). Efforts to restore their functionality have largely been unsuccessful to date (Chiocca et al., 2019; Meister et al., 2022). The accumulation and activity of CD8^POS^ T cells, which are essential for effective tumor clearance, are tightly regulated to minimize damage to healthy tissues and immunopathology by balancing stimulatory signals through the TCR and cytokine as well as co-stimulatory receptors with inhibitory signals binding to the immune checkpoint and metabolite CTL receptors (Zagorulya and Spranger, 2023). To compensate for the inhibitory and immunosuppressive signaling dominant in GB and shift the balance in favor of activating signals in the TME, we therapeutically augmented the pro-inflammatory cytokine rIL-12 and the co-stimulatory factor 4-1BBL. Although both factors are well-described in oncology, little is known about their effect on GB. This is partly due to the fact that historically the initial enthusiasm for pro-inflammatory agents such as IL-12 has declined due to its association with dose-dependent systemic toxicity observed in animal studies (Zou et al., 1995) and clinical trials (Chiocca et al., 2019; Motzer et al., 2001). Recently, alternative approaches for safely administrating therapy with pro-inflammatory stimuli have shown potential, including spatially controlled delivery and short-term administration methods (Barnwal et al., 2023; Nguyen et al., 2020; Venkatas and Singh, 2022). Therefore, we focused on administering a single low dose of soluble rIL-12 at the tumor site and utilizing AAV vector technology to locally display 4-1BBL, evaluating their anti-tumor effects and improved survival in preclinical GB mouse models.

Tumor-associated CD8^POS^ T cell responses are regulated by intratumoral DCs. DCs are well-equipped to process and convert information from their surroundings into tailored instructions to guide T cell responses (Cabeza-Cabrerizo et al., 2021). DC-derived stimulatory signals driving T cell proliferation and activation include 1) Ag peptide-mediated presentation by MHC molecules (e.g., MHC-I and MHC-II), 2) co-stimulatory ligands (e.g., CD80, CD86 and 4-1BBL), and 3) cytokines (e.g., IL-12 and type I IFNs) (Eisenbarth, 2019; Fu and Jiang, 2018; Habib-Agahi et al., 2007; Zagorulya and Spranger, 2023). However, in GB tumors, DC function and numbers are suppressed (Friedrich et al., 2023), which has implications for anti-tumor immunity (Tooley et al., 2022). We questioned whether the lack of anti-tumor reactivity was due to insufficient DC instructions, dysfunction of CD8^POS^ T cells, or both. This was investigated with isolated murine GB-associated CD8^POS^ T cells from mouse brains. Tumor-infiltrating CD8^POS^ T cells retained the ability to secrete IFN-γ following MHC-I presentation of mImp3 antigen by naïve DCs (Schaettler et al., 2023), indicating the presence of a tumor-Ag-specific TCR. However, this response was only achieved when the CD8^POS^ T cells were stimulated with rIL-12, suggesting that the intricate engagement between DCs and CD8^POS^ T cells necessary for activation is insufficient in the GB TME.

We then explored which functional DC states, when functional, provide the necessary Ag cross-presentation and immunomodulatory signals to CD8^POS^ T cells in GB. By processing and analyzing of available scRNAseq datasets indicated that a discrete CCR7^POS^ DC state in GB tumors expresses *Il12b* and high levels of the MHC-I machinery, along with high levels of co-stimulatory molecules and pro-inflammatory cytokines essential for CD8^POS^ T cell activation. The CCR7 signature suggests that these DCs are migratory (Liu et al., 2021), involved in capturing tumor-associated Ags within the TME and subsequently trafficking to the draining lymph nodes to present Ags to CD8 T cells (Eiraku et al., 2018; Eisenbarth, 2019). These tumor-retained DCs that we referred to here as *Il12b*^POS^DCs arise from cDC precursors and have been differently named in other tumor types, including DC3, migratory DCs, mRegDCs, as well as CCR7^POS^ DCs, LAMP3^POS^ DCs, mature DCs, or activated DCs (Di Pilato et al., 2021; Lee et al., 2024). Here, we found that in addition to their IL-12 production potential (Brandum et al., 2021), other key immune modulators were expressed by these DCs, such as co-stimulatory IL15/RA, CXCL16, 4-1BBL, CD80, and inhibitory PD-L1/2 (Ziblat et al., 2024). Interestingly, the receptors for IL-12 and 4-1BBL products were also intrinsically expressed in this DC state, suggesting that there might be a positive feedback loop to enhance their activation and thereby promote a more robust anti-tumor immune response of CD8^POS^ T cells. Indeed, when exogenous rIL-12 was supplied to the tumor, it induced *Il12b*^POS^DC accumulation and pSTAT4, suggesting that DCs sense IL-12, or their activation could be indirectly through other cells that can sense IL-12. This has been reported by other studies where IL-12R in DCs stimulates autocrine signaling to maintain IL-12 expression (Grohmann et al., 1998; Nagayama et al., 2000; Tugues et al., 2015). A similar effect can be envisioned for exogenous 4-1BBL expression and 4-1BB receptor expressing *Il12b*^POS^DCs (Macdonald et al., 2014). The underperformance of *Il12b*^POS^DCs in GB, despite their presence in scRNAseq datasets, warrants further investigation. Our data suggests that these tumor-residing DCs are rare in mouse GB and even more limited in *de novo*/recurrent glioma patient datasets, detectable only through specific enrichment protocols. Next to their limited presence compared to other types of cancer, such as melanoma, *Il12b*^POS^ DCs also express the highest PD-L1 levels whenever they become activated, imposing suppression on CD8^POS^ T cells. These observations, together with the possibility that both DCs and CD8^POS^ T cells are not abundant enough to contribute effectively to the anti-tumor immune response, likely explain why GB tumors with high versus low rIL-12 or 4-1BBL expression do not show differences in patient survival. Therefore, survival data may not reflect the potential effect of supplementing exogenous IL-12 or 4-1BBL, which could provide the necessary signals to activate immune pathways that are otherwise absent.

Besides affecting DCs, IL-12 and 4-1BBL have been suggested to activate CD8^POS^ T cells (Reithofer et al., 2021), resulting in the release of cytotoxic molecules, such as granzymes (Lasek et al., 2014), but can also affect other immune cells, such as NK cells (Bosch et al., 2019; Parihar et al., 2002), and regulatory/helper cells (Cao et al., 2009). With the aid of human and murine scRNAseq datasets, we confirmed the expression of IL12Rβ1/2 and 4-1BB in these cell types and, with flow cytometry, we demonstrated that they have the potential to phosphorylate STAT4 upon rIL-12 treatment. Nonetheless, antibody-based depletion of CD8 T cells, CD4 T cells, and NK cells in our GB models illustrated that CD8^POS^ T cells effectuate the rIL-12-induced immunity and not the other cell types. This aligns with findings in melanoma models in transgenic mice, where it was demonstrated that NK and CD4 T cells could potentially have a supportive function but are less likely to be directly involved in IL-12-mediated anti-tumor immunity (Kerkar et al., 2011). In the context of human GB, the VDX trials showed that besides the IFN-γ signature, predominantly the number of CD8 T cells, and not the CD4 T cells, increased at the tumor site, not the CD4 T cells (Chiocca et al., 2019). Upon local rIL-12 exposure of the GB tumor, the recruited CD8^POS^T cells differentiated towards a more effector-like CD8^POS^ T cell state. This effector-like state exhibited high levels of PD-1, GZM-B, and 4-1BB expression and increased STAT-4p upon a rIL-12 trigger. Overall, our data suggest that rIL-12 influences both DCs and T cells. Specifically, CD8^POS^ T cells are directly affected by rIL-12, while their activity is indirectly supported by *Il12b*^POS^ DCs, which provide the necessary additional signals.

Given that CD8^POS^ T cells at the tumor site express 4-1BB and IL-12Rβ1/2 and are tumor-antigen specific, their activation is likely limited by the scarcity or immunosuppressed state of *Il12b*^POS^ DCs. To overcome this limitation, we investigated an alternative model that delivers stimulatory signals through mechanisms independent of DCs. We demonstrated that 4-1BBL-overexpression by tumor cells increased the survival of rIL-12 treated animals, and we could recapitulate these results with AAV vector-mediated m4-1BBLgene therapy. It was shown with CD8 depletion in *Il12^+/+^* and/or *Il12^−/−^*mice that the effect was CD8^POS^ T cell mediated and not dependent on functional DCs. Other attempts with a combination of IL-12 and 4-1BBL have, to our knowledge, only been attempted with other cancers, such as colon carcinoma (Chen et al., 2000), liver metastasis (Martinet et al., 2000), and melanoma (Huang et al., 2010).

In contrast to tumoricidal drugs that act on tumor cells directly, AAV vector-based immunotherapy can also act in the vicinity of the tumor as the transgene products can work indirectly, e.g., through CD8^POS^ T cells (Binnewies et al., 2018). Recently, a successful AAV vector therapy approach has been deployed targeting endothelial cells in the vasculature of GB with reduced CD8^POS^ T cells (Ramachandran et al., 2023). In our strategy, 4-1BB^POS^ CD8^POS^ T cells recruited by rIL-12 were stimulated by AAV vector-mediated delivery. 4-1BB stimulation with agonist antibodies has shown promising effects on CD8^POS^ T cells in patients with advanced solid tumors (Melero et al., 2023), B cell lymphoma (Souza-Fonseca-Guimaraes et al., 2016), or pancreatic cancer (Gulhati et al., 2023). Our strategy created a continuous reservoir of 4-1BB stimulation at the tumor site, where CD8^POS^ T cells reside. This was achieved by packaging the AAV vector with an astrocyte-tropic AAVF capsid and using a GFAP promoter to drive the transgene, with GFAP being elevated in slow-dividing reactive astrocytes associated with the tumor (Beharry et al., 2022; Hanlon et al., 2019; Yao et al., 2022) and, to a certain extent, in proliferating tumor cells. This reduced expression in tumor cells is likely due to factors such as increased proliferation contributing to AAV vector genome dilution (Colella et al., 2018). GB models with higher GFAP expression in and around the tumor, such as GL261 and 005, performed better than CT-2A, which exhibited limited GFAP expression. Our vector was injected into the tumor cavity over three consecutive days to achieve sufficient therapeutic transgene expression and avoid immune-mediated elimination of the AAVF capsid. The boost of rIL-12 immunity by our local transgene expression prolonged overall survival in all our tested GB models and was shown to be more effective than ICI, such as anti-PD-L1.

In conclusion, we have demonstrated an immuno-gene therapy combination strategy whereby CD8^POS^ T cells were activated at the tumor site with limited cues from tumor-associated DCs, leading to increased survival in GB tumor-bearing mice. These findings are crucial for GB patients, as they potentially offer improved outcomes compared to ICI therapies and support future studies using models with human GB tissues.

## METHODS

### Experimental model and subject details

#### Animals

All animal experiments were performed in agreement with ethical guidelines of the National Institutes of Health for the Care and Use of Laboratory Animals. Experiments were conducted under the oversight of the Massachusetts General Hospital Institution Animal Care and Use Committee (IACUC). C57BL/6J were purchased from Charles River Labs (IACUC protocol 2009N000054). T.R.M. provided *Il12b^tm1.1Lky^/J* (IL-12 p40-YFP) mice (Reinhardt et al., 2006) and the B6.129S1-*Il12b^tm1Jm^*/J (IL-12p40 KO) mice (Magram et al., 1996). Animals were maintained in specific pathogen-free facilities at Massachusetts General Hospital (MGH) with unlimited access to water and food under a 12-hr light/dark cycle. To study the immunomodulatory effects of rIL-12 and 4-1BBL on GB *in vivo,* C57BL/6J adult male and female mice were randomly assigned to each group.

#### In vivo bioluminescence analysis

*In vivo* tumor growth in brain was monitored by Firefly luciferase by BLI using a Xenogen *in vivo* 200 Imaging System (IVIS) (PerkinElmer). D-Luciferin (Gold Biotechnology) was reconstituted by adding 50 ml of 1x sterile phosphate-buffered saline (PBS) to the lyophilized pellet. 100 μl working solution was injected i.p. in mice. Imaging was acquired 5 min after injection and analysis was performed using Living Image software 4.3.1 (PerkinElmer).

#### Cell culture

The National Cancer Institute (NCI) provided mouse GB cells (CT-2A, GL261 and 005) syngeneic with strain C57BL/6J. HEK293T cells were purchased from ATCC (Locarno et al., CRL-3216™). Cells were cultured at 37°C in a 5% CO_2_ humidified incubator. CT-2A cells were cultured in Dulbecco’s modified Eagle’s medium (DMEM; Corning) supplemented with penicillin (100 units/mL) and streptomycin (100 mg/mL) (P/S) (Corning) and 10% fetal bovine serum (FBS) (Gemini Bioproducts, West Sacramento, CA). GL261 cells were cultured in Roswell Park Memorial Institute (RPMI) (Corning) with 10% FBS and 1% p/s. 005 cells were cultured in DMEM Nutrient Mixture F-12 (DMEM/F-12, Gibco Thermofisher). DMEM/F-12 was supplemented with 1% penicillin/streptomycin, B-27 supplement (1x, Gibco Thermofisher), heparin (Sigma-Aldrich) (2 μg/mL), epidermal growth factor (EGF, R&D system) (20 ng/mL), and fibroblast growth factor (FGF, Peprotech) (20 ng/mL). Cells tested negative for mycoplasma contamination at periodic intervals throughout the study (Mycoplasma PCR Detection Kit G238; ABM, Richmond, BC, Canada).

For *in vivo* experiments, GB cells (CT-2A, GL261, and 005) cells were stably transduced with an LVV vector to express FLuc (Addgene #108542) and were used for all subsequent *in vivo* experiments.

The FLuc plasmid was obtained from Addgene (Cat#108542) and was transfected into HEK293T cells along with capsids and packaging material for lentivirus production. HEK293T cells were cultured for 24 hrs and fresh media was provided 72 hrs after transfection for lentivirus production. The conditioned media was spun down at 300 x g for cell debris removal. The supernatant was filtered through a 0.2 µm filter and the virus was pelleted at 330,000 x g for 2 hrs. All virus preparations were aliquoted and kept frozen at −80 degrees until use. The FLuc virus was transduced into CT-2A, GL261 and 005 cells and transduced cells were selected with blasticidin.

To study the effect of 4-1BBL, GB cells (CT-2A-FLuc and 005-FLuc) were stably transduced with an LVV to express 4-1BBL under the GFAP promoter tagged 3xFLAG-tag and with fluorescent label mCherry or the control vector lacking 4-1BBL.

### Method details

#### Intracranial tumor implantation

Adult mice were anesthetized using 2.5% isoflurane (USP, Baxter Healthcare Corporation) in 100% oxygen via a nose cone and placed on a warm pad to avoid hypothermia. A total of 5 × 10^4^ CT-2A-FLuc were suspended in 1 µL Opti-MEM (Gibco, Waltham, MA). In total 2 µL of the cell suspension was then implanted into the left striatum of C57BL/6J mice, IL-12 p40-YFP or IL-12p40 KO (*Il12^−/−^*) mice using a Hamilton syringe (Sigma-Aldrich, Germany) and automatic stereotaxic injector (Stoelting, Wood Dale, IL) with a flow rate of 0.2 µl/min for 10 min. In reference to bregma, three coordinates for stereotactic implantation were chosen: anterior-posterior (AP) = 2.0 mm, medial-lateral = 0.5 mm, and dorsal-ventral = 2.5 mm. Overall survival of the mice was based on 20% weight loss, presence of apparent distress, or actual death. Tumor growth in mice was assessed by measuring BLI using IVIS (PerkinElmer, Waltham, MA) every three or four days starting from day 7 after tumor implantation.

#### Survival studies

##### IL-12 and AAVF vector treatment

For CT-2A-FLuc tumors, ten days after intracranial injection, mice were treated with either (5, 20, 50, 200, or 500 ng) rIL-12-FC (Adipogen - Cat#: CHI-MF-11112-C025) or FC (Adipogen - Cat#: CHI-HF-210IG1-C100) (50 ng) sham control by i.t. injections using a Hamilton syringe (Sigma-Aldrich, Germany) and automatic stereotaxic injector (Stoelting, Wood Dale, IL) with a flow rate of 0.2 µl/min for 10 min at the coordinates used for tumor implantations. For GL261-FLuc and 005-FLuc tumors, 50 ng rIL-12 or sham were injected i.t at day 10 post-tumor implantation.

For three mouse GB models (CT-2A, GL261, 005) AAVF vectors (AAVF-m41BBL and AAVF-null) were intracranially injected in a volume of 5 µL (5.0 × 10^13 genome copies (gc)/mL at three time points. One day prior to tumor implantation (day −1), on day 0 (tumor implantation) and one day post-tumor implantation (day 1) the mice were injected with the AAVF vector using a Hamilton syringe (Sigma-Aldrich, Germany) and automatic stereotaxic injector (Stoelting, Wood Dale, IL) with a flow rate of 0.2 µl/min for 10 min. In reference to bregma, three coordinates for stereotactic implantation were chosen: anterior-posterior (AP) = 2.0 mm, medial-lateral = 0.5 mm, and dorsal-ventral = 2.5 mm. The same coordinates were used for all three AAVF vector injections.

##### Depletion of CD8, CD4 T cells or NK1.1 cells

To deplete CD8 T cells, CD4 T cells or NK1.1 cells in C57BL/6J and *Il12^−/−^*mice, endogenous CD8, CD4 T or NK1.1 ells were depleted by i.v. injection of anti-mouse CD8β antibody (Bioxcell, Clone Lyt 3.2) anti-mouse CD4 antibody (Bioxcell, Clone GK1.5), anti-mouse NK1.1 antibody (Bioxcell, Clone PK136) or rat IgG2b isotype control (Bioxcell, Clone LTF-2) on day 9 (50 µg) and day 10 (100 µg) post-tumor implantation. On day 10, rIL-12 or sham was injected i.t. at the tumor site and mice were sacrificed on day 18 for flow cytometry of dissociated brain cells.

##### Anti-PD1/PD-L1

CT-2A tumor bearing mice were administered with IP injections as days 3, 5 and 14, with anti-PD-L1 (Leinco Technologies, clone 10F.9G2) or with rat IgG2b isotype control (Bioxcell, Clone LTF-2) at a dose of 200 µg/mouse in a volume of 100ul. Tumor growth was measured by IVIS every 3-4 days and mice were euthanized based on 20% weight loss, presence of apparent distress, or actual death.

Whole blood was retro-orbital collected ∼400 µL and was sent to pathology core on day 15 for pathology toxicology analysis.

##### Retroorbital blood collection

To identify the depletion of CD8 T cells in the blood post-i.v. injection of anti-CD8 antibody, 100 µl retro-orbital blood was collected via 1.2mm size glass capillaries (world precision instruments) in EDTA tubes to avoid coagulation on days 7, 11 and 18 post-tumor implantation. The collected blood was further processed immediately for RNA isolation.

##### Whole Blood collection

Mice were sacrificed by a lethal 100 µL i.p. injection containing ketamine (5 µL), xylazine (45 µL) and saline (50 µL) (Patterson Veterinary). Upon ceasing of all reflexes, whole blood was collected directly from the heart in EDTA tubes. The blood was processed immediately by centrifugation at 1500 x *g* for 15 min to pellet the blood cells. The blood cell pellet was washed with PBS carefully and used for RNA isolation to determine the expression of CD8 in whole blood cells. The supernatant was carefully centrifuged again at 2500 x *g* to collect the plasma. The plasma was further analyzed at the Pathology core at MGH, with the comprehensive blood toxicology panel.

##### AAV plasmid constructs and production

The m4-1BBL expression construct was cloned into a GFAP-GFP AAVF vector plasmid (AltaBiotech), using the restriction enzymes NheI-HF and NcoI-HF (New England Biolabs) followed by Gibson assembly with NEBuilder® HiFi DNA Assembly Master Mix (New England Biolabs). Both the AAVF-m41BBL and AAVF-null vector plasmids were then transformed into SURE Electroporation Competent cells (Agilent Technologies) by two pulses at 1700 V. Plasmid DNA was isolated in nuclease-free water (Ambion Life Technologies) using the Qiaprep® Spin miniprep kit (Qiagen) after selection with 1 µg/mL ampicillin (ampicillin sodium salt, Sigma). Both plasmid constructs were fully sequenced with Next Generation Sequencing at the MGH CCIB DNA core and analyzed with Snapgene software version 6.0.2. Upon confirmation of the sequence, the plasmid constructs were isolated at a large scale by AltaBiotech at a concentration of 2 µg/µL. Subsequently, scAAVF vectors were produced by Packgene at a titer of 1.0 × 10^13 genome copies (gc)/mL.

##### Western blots

Total protein was extracted from cultured cells using RIPA lysis buffer (Thermo Scientific). The tissue samples were homoginized in RIPA lysis buffer with a tissue homogenizer. RIPA buffer was supplemented with a protease inhibitor cocktail (Sigma-Aldrich). To remove non-soluble cell debris, samples were sonicated using a probe sonicator (Sonic Dismembrator Model 100, Fisher Scientific) at a setting of 3.0 for 5 sec and centrifuged at 15,000 *x* g for 10 min at 4°C. Protein concentration was determined using the Pierce^TM^ BCA Protein Assay Kit (Thermo Fisher Scientific). Absorbance was measured at 562 nm using the SynergyHI microplate reader (BioTek). Equal amounts of protein (20 µg) mixed with Laemmli SDS-Sample buffer (Boston BioProducts) were loaded and resolved by electrophoresis on NuPage® 4-12% Bis-Tris polyacrylamide gels (Thermo Fisher Scientific) in NuPage® MES SDS Running Buffer (Thermo Fisher Scientific). After transfer onto nitrocellulose membranes using the iBlot 2 (Thermo Fisher Scientific), samples were subsequently incubated for one hr at RT in 5% non-fat dry milk (Labscientific) in Tris-buffered saline (pH = 7.4) with 0.05% Tween 20 (TBS-T) and probed with primary antibody mouse 3xFLAG-tag 1:1000 (Merck, F3165) or goat-α-β-actin (Santa Cruz Biotechnology, I-19) overnight at 4°C. After washing three times with TBS-T for 10 min, membranes were incubated for one hr at RT with secondary antibodies ECL^TM^ donkey-anti-goat immunoglobulin G (IgG) (Sigma-Aldrich) and ECL^TM^ sheep-anti-mouse IgG (Thermo Fisher Scientific) (1:5000) corresponding to the primary antibodies. Membranes were developed with ECL or Femto staining (Thermo Fisher Scientific) and imaged on an Azure Biosystems C300 gel imager.

##### Luminex

To quantify protein concentrations in C57BL/6J mice and IL-12p40 KO (*Il12^−/−^*) mice, frozen tumor-bearing brains were cut in pieces around the tumor and weight in microcentrifuge tubes and 100µL of RIPA lysis buffer (Thermo Scientific) was added per 100 mg of tissue. Stainless steel 3mm tungsten carbide beads (Qiagen) were added to homogenize the tissue using the TissueLyser system (Qiagen) for 3 min at 0.25 Hz speed. Samples were centrifuged at 16,000 × g for 10 min at 4°C. Supernatant were transferred to new microcentrifuge tubes. Protein concentration was determined using the Pierce^TM^ BCA Protein Assay Kit (Thermo Fisher Scientific) and samples were diluted to 10 mg protein/mL with 1X PBS. To proceed with ProcartaPlex mouse basic kit (Invitrogen) protocol, 25 µL of Universal Assay Buffer (Cat. No. EPX-11111-000) was added to 25 µL of the diluted sample per sample well. Samples were incubated with beads overnight and analysed by flowcytometry to detect events in the PE channel.

##### RT-qPCR

Total RNA was extracted using the Direct-Zol RNAmini kit (Zymo-research). RNA concentrations were measured using the Nanodrop Spectrophotometer ND-1000 (Thermo Fisher Scientific). For gene expression analysis using RT-qPCR, cDNA was synthesized from 200ng total RNA and prepared using the SuperScript® Vilo^TM^ cDNA Synthesis Kit (Thermo Fisher Scientific). cDNA samples were diluted 10-fold with nuclease-free water. Gene expression was determined using the manufacturing protocol of PowerUp^TM^ SYBR^TM^ Green PCR Master Mix (Applied Biosystems). The cycling conditions using the standard protocol were: 2 min at 50°C, 10 min at 95°C, 40 cycles of 95°C for 15 sec and 60°C for 1 min, followed by a melt curve from 60 to 95 °C at 0.1 °C/sec, with 15 sec hold at 95°C. Twenty-five sets of primers (**Table 4**) obtained from Origene (https://www.origene.com/) were used to specifically target the genes of interest by RT-qPCR. Gene expression was normalized to the housekeeping mRNA β-Actin.

##### Tissue digestion

The mice were exsanguinated further with PBS perfusion. Tumor Tissue Dissociation Kit (MiltenyiBiotec) was used to process the brain into a single-cell suspension. Brains were placed into a GentleMacs C-tube (Miltenyi Biotec) with 2.35 mL RPMI 1640 (Corning) containing enzymes D (100 μl), R (30 μl) and A (3.5 μl). According to the manufacturer’s protocol, the brains were dissociated using the gentle MACS Dissociator (Miltenyi Biotec) on the brain program settings. Samples were run through a 70 µm filter to obtain a single-cell suspension. Myelin removal was achieved using magnetic separation and anti-myelin beads (Miltenyi Biotec). The final cell suspension was resuspended in 1X Dulbecco’s (D)PBS without calcium (Ca^2+^) or magnesium (Mg^2+^) (Corning), supplemented with 2 mM EDTA (Thermo Fisher) and 0.5% BSA (Sigma). Samples were then loaded onto a series of LS columns containing microbeads conjugated to anti-mouse CD11b and anti-mouse CD45 (Miltenyi Biotec), respectively, and separated into CD11b^POS^, CD11b^NEG^CD45^POS^, and CD11b^NEG^CD45^NEG^ (non-immune) cell populations using the MACS multi-stand (Milteny Biotec).

##### Antibody staining and flow cytometry

Cell surface proteins were stained for 20 min at 4°C. Intracellular and nuclear proteins were stained for 60 min at RT after permeabilization and fixation (Thermo Fisher Scientific) for 30 min at RT. To investigate T cells, samples were stained with different antibodies (**Table 5**). Stained cell samples were re-suspended in 200 µL FACS buffer (Dulbecco’s PBS supplemented with 2 mM EDTA and 0.5% FBS) and transferred to FACS tubes (Stellar Scientific). A mixture of isolated lymph nodes derived from the thigh and spleens were passed through 70 mm cell strainers, pellets were then incubated with red blood cell lysis buffer (Boston Bioproducts) two times for 5 min and washed with PBS. These lymph nodes and splenocyte mixtures were used as single-stained controls. In all experiments, lymph nodes, spleens and ipsilateral hemispheres implanted with CT-2A cells were mixed to measure the Fluorescence Minus One (FMOs). For all studies, dead cells were stained using the fixable viability violet dyes - Zombie Red or Zombie Blue (Invitrogen) for 10 min at room temperature (RT), followed by blocking of Fc receptors with TruStain fcX (Biolegend) for 15 min at 4°C. Cells were analyzed on LSRFortessa^TM^ or LSRFortessa X-20 flow cytometers (BD Biosciences) and data was analyzed with FlowJosoftware version 10.8.1.

##### Immunohistochemistry

Whole brains from mice were fixed overnight in 4% paraformaldehyde at 4°C. The following day, brains were transferred to a 30% sucrose (Sigma) solution and incubated until they sank, indicating proper cryoprotection. Brains were then embedded in optimal cutting temperature compound (Fisher Scientific) and snap frozen. Serial coronal sections (12 µm thick) were prepared using a cryostat and mounted onto Fisherbrand microscope slides (Canada). The sections were fixed again with 4% paraformaldehyde for 10 min at RT, followed by three 5 min rinses in PBS. Blocking was performed for one hr at RT in blocking buffer consisting of 5% goat serum and 0.1% Tween-20 in PBS (PBS-T). Brain slices were then incubated with the primary antibodies (GFP 1:400, Invitrogen Cat#A11120; GFAP 1:400, Invitrogen Cat#13-0300; CD8 1:400, Novus Biologicals Cat#NBP2-29475; IL12Rb1 1:400, Invitrogen Cat#PA5-95976; anti-4-1BB 1:100, Absolute Antibody Cat# Ab01052; 3xFLAG-tag 1:400, Abcam Cat# ab245893; 4-1BBL 1:100, Invitrogen Cat#MA529838; CD11c 1:400, Abcam Cat#ab33483), diluted in blocking buffer at 4°C overnight. Slices were rinsed three times in PBS-T for 5 min each. Secondary antibodies (goat anti-rabbit 1:400 Invitrogen Cat#A11008; goat anti-rat 1:400 Abcam Cat#ab150157; 1:400 goat anti-mouse Invitrogen Cat#A11001) were diluted in PBS-T and incubated for one hr in the dark at RT. Slices were mounted with DAPI (Vectashield, Vector Labs, San Francisco, CA).

##### Hematoxylin & Eosin staining

For H&E staining, brain slices were air dried under a fan for 20 min, before fixation in 100% ethanol for 10 min. Brains were rinsed briefly in MilliQ (EMD Millipore), then stained for 10 min at RT with Harris Hematoxylin (Poly Scientific R&D). Slides were washed twice with MilliQ for 2 min, then de-stained in 1% acetic acid (Sigma-Aldrich) for 6 sec, followed by washing twice in MilliQ. Samples were differentiated in 0.05% aqueous lithium carbonate (Poly Scientific R&D) for 30 sec, after which they were washed in warm tap water for 2 min. 1% Eosin Y solution (Electron Microscopy Sciences) was pipetted on top of the sections to counterstain for 4 sec. Next, brains were de-stained in 95% ethanol for 20 sec, followed by further de-staining and dehydration in 100% ethanol for 5 min. Brain sections were cleared in Xylene (Sigma-Aldrich) for 15 min, mounted with Permount (Electron Microscopy Sciences) and imaged on a Keyence microscope at 4x magnification.

##### IFN-γ Elispot assay

GL261 tumor-bearing mice were sacrificed on day 14 post-tumor implantation and the ipsilateral hemisphere was dissociated into single-cell suspensions. Tumor single cell suspensions were separated from myelin using magnetic separation and anti-myelin beads (Miltenyi Biotec). The myelin-negative cell pellet was incubated with microbeads conjugated to anti-mouse CD45 (Miltenyi Biotec). After magnetic separation, the CD45 positive cell pellet was incubated with CD8 microbeads (Miltenyi Biotec) to isolate for positive CD8 T cells. CD8^POS^ T cells were cultured in RPMI 1640 (Corning) with 10% FBS and 1% p/s, 1% Glutamax (Gibco) and 0.01% 2-mercaptoethanol (Thermo Fisher) stimulated with IL-2 overnight. Splenocytes were derived from a spleen that was filtered in PBS through a 70mm cell strainer followed by incubation with red blood cell lysis buffer (Boston Bioproducts) two times for 10 min. CD8^POS^ T cells and splenocytes were counted 150,000 CD8^POS^ T cells combined with 25,000 splenocytes were plated in a 3:1 ratio either with or without mImp3, GL261-specific neopeptide (AALLNKLYA) together with either 50ng FC or rIL-12 overnight in 200ul RPMI 37C on a pre-coated murine IFN-γ detection plate (ImmunoSpot). After following manufacturers’ protocol, wells were dried overnight, and images were quantified by Image J.

##### Cell viability assay

Cell proliferation was assessed *in vitro* by the WST reduction assay to determine cell viability (cell counting kit-8; Dojindo, Rockville, MD) of FACS-sorted GFP^POS^ cells. Cells were seeded at a low density (2 × 10^3^ cells/well) in a 96-well plate. After 24 hrs, the medium was removed, and 10% WST solution was added to the cells. The cells were incubated at 37°C for one hr, and absorbance levels at wavelength 450 nm were measured using a microplate reader (SynergyH1; BioTek, Winooski, VT). Thereafter, the medium was changed, and cells were measured repeatedly every 24 hrs up until 90% confluency on day 5.

#### Single-cell RNA sequencing analysis

##### scRNAseq datasets tumor-bearing mice/human samples enriched for immune cells

For the scRNAseq analysis, publicly available datasets, or datasets provided by co-authors were used (**Table 1**). The Seurat v4-v5 R package was used to preprocess and analyze the data (Hao et al., 2021; Hao et al., 2024). Unless otherwise stated, the Seurat Pipeline was followed. Low-quality cells were excluded from the analysis. The count matrix and cell metadata were used to create a Seurat object, of which the standard Seurat Pipeline was followed by running NormalizeData, FindVariableFeatures (using variance stabilizing transformation), ScaleData, RunPCA, FindNeighbors, FindClusters and RunUMAP. Cell types were annotated using published cell annotation matrices (Pombo Antunes et al., 2021) and projected on the other datasets to homogenize analyses between datasets. To examine the expression levels of genes of interest, “VlnPlot”, “FeaturePlot” and “AverageExpression” functions from Seurat were used. The proportion of cells per cluster that expressed genes of interest (normalized counts > 0) was also calculated.

For feature plots, UMAP visualizations and heat map analysis, we utilized a subset of the scRNAseq data from our preprint Miller et al., 2023, specifically incorporating data derived from Johnson *et al.,* (Johnson et al., 2021) and Abdelfattah *et al.,* (Abdelfattah et al., 2022). Cell annotations were applied according to Miller et al. 2023 to ensure consistent classification. The data was normalized to 10,000 counts per cell, log-transformed, and the top 3,000 most highly expressed genes were selected for dimensionality reduction and downstream analysis. Principal Component Analysis (PCA) was used for dimensionality reduction, and a nearest neighbors’ graph was constructed with standard parameters (n_pcs=40, n_neighbors=10). Uniform Manifold Approximation and Projection (UMAP) was subsequently applied for visualization. All analyses and visualizations were conducted in Python using the Numpy, Pandas, and Scanpy (Wolf et al., 2018) libraries.

The survival information (Survival time) and Events were obtained from the G-SAM (Hoogstrate et al., 2023) and GLASS (Consortium, 2018) cohorts. IDH-WT GBMs were exclusively considered for this analysis. Duplicate patient entries were excluded, and the values were maintained from the primary tumor only for the patient. For genes, the CPM-normalized value was used. For gene sets, the CPM-normalized and log-transformed matrix was uploaded to Seurat, and Module scores of gene sets were calculated using the AddModuleScore() function. CIBERSORTx (Newman et al., 2019) was used to normalize the expression of genes or module scores to the myeloid contents in the cohorts. Discretized matrix was utilised from our preprint (Miller et al., 2023) as a reference matrix for CIBERSORTx. We removed any library with a CIBERSORTx value of 0 for the myeloid lineage. Samples in the top 33% in terms of expression of genes of interest (or module scores) were labeled as “high.” The bottom 33% were considered the “low” group. We used ggsurvfit (https://github.com/pharmaverse/ggsurvfit) to generate the Kaplan-Meier survival curve. A Cox Proportional Hazard Model (https://github.com/therneau/survival) was used to determine differences in survival probabilities.

### Quantification and statistical analysis

Bar graphs, heat maps, and survival plots were made in GraphPad Prism 9.5.1. Error bars show the mean ± standard error of the mean (SEM). A one way ANOVA, two way ANOVA, multiple t-tests, and Log-rank tests were applied to determine if conditions significantly differed. Statistical significance was specified as p <0.05. Sequences and plasmid constructs were analyzed with Snapgene software version 6.0.2.

## Supporting information

List of Primers for the manuscript

Resource Table for Antibodies

## Acknowledgments

We thank all members of the Breakefield laboratory for their suggested ideas during laboratory meetings. We thank all laboratories within the Molecular Neurogenetics Unit at MGH for their input. We thank the Dunn’s laboratory for generously providing the mImp3 peptide. We would like to thank Dr. Mark Issa for his expert advice on immunology within the tumor microenvironment. Special thanks to members of the Mempel laboratory for sharing their expertise on the immune system and El Khoury’s laboratory for sharing their insight on tumor-associated microglia. We thank Mrs. Suzanne McDavitt for her skilled editorial assistance. X.O.B. acknowledges grant support from National Institute of the Neurological Disorders and Stroke (NINDS) NS122163, the National Institutes of Health (NIH) National Cancer Institute (NCI) CA179563, CA069246, and CA232103 and NINDS NS122163 grants for supporting this work. U19 CA179563 was supported by the NIH Common Fund, through the Office of Strategic Coordination/Office of the NIH Director. T.R.M. acknowledges grant support from NIH grant R01 AL123349. K.B. acknowledges grant support from Cell-immunotherapy for brain tumors R01 NS122163-01A1. T.R.L. acknowledges grant support from the Norwegian Research Council (Grant Number: 315566). L.N. acknowledges support from Prins Bernhard Cultuurfonds. S.M.L. acknowledges the Jo-Kolk scholarship.

## Author contributions

X.O.B. and K.B. conceived the study and designed experiments. X.O.B. and K.B. supervised the project. T.S.v.S. initiated the project. K.B., T.R.L., L.N., S.M.L, T.X., and E.D. performed and analyzed the experiments. K.B. designed the plasmids and D.R.R., S.M, and E.G., cloned the constructs. T.R.M., provided transgenic mice strains. T.R.M, J.K.L, and M.S., provided their expertise on the flow cytometry experiments, interpretation and analysis. K.B., V.M., and A.J.E.M., performed bioinformatic analyses. K.B., C.P.C., C.A.F., and T.E.M., analyzed human scRNAseq datasets. E.W., Y.S., and R.W.J., performed organotypic culture experiments. G.D., provided the mImp3 peptide and M.L., supported with the Elispot assay. L.N., T.R.L., and K.B. prepared the figures. T.R.L., L.N., and K.B. wrote the manuscript. All authors edited or commented on the manuscript.

---

**Supplemental Figure 1:**
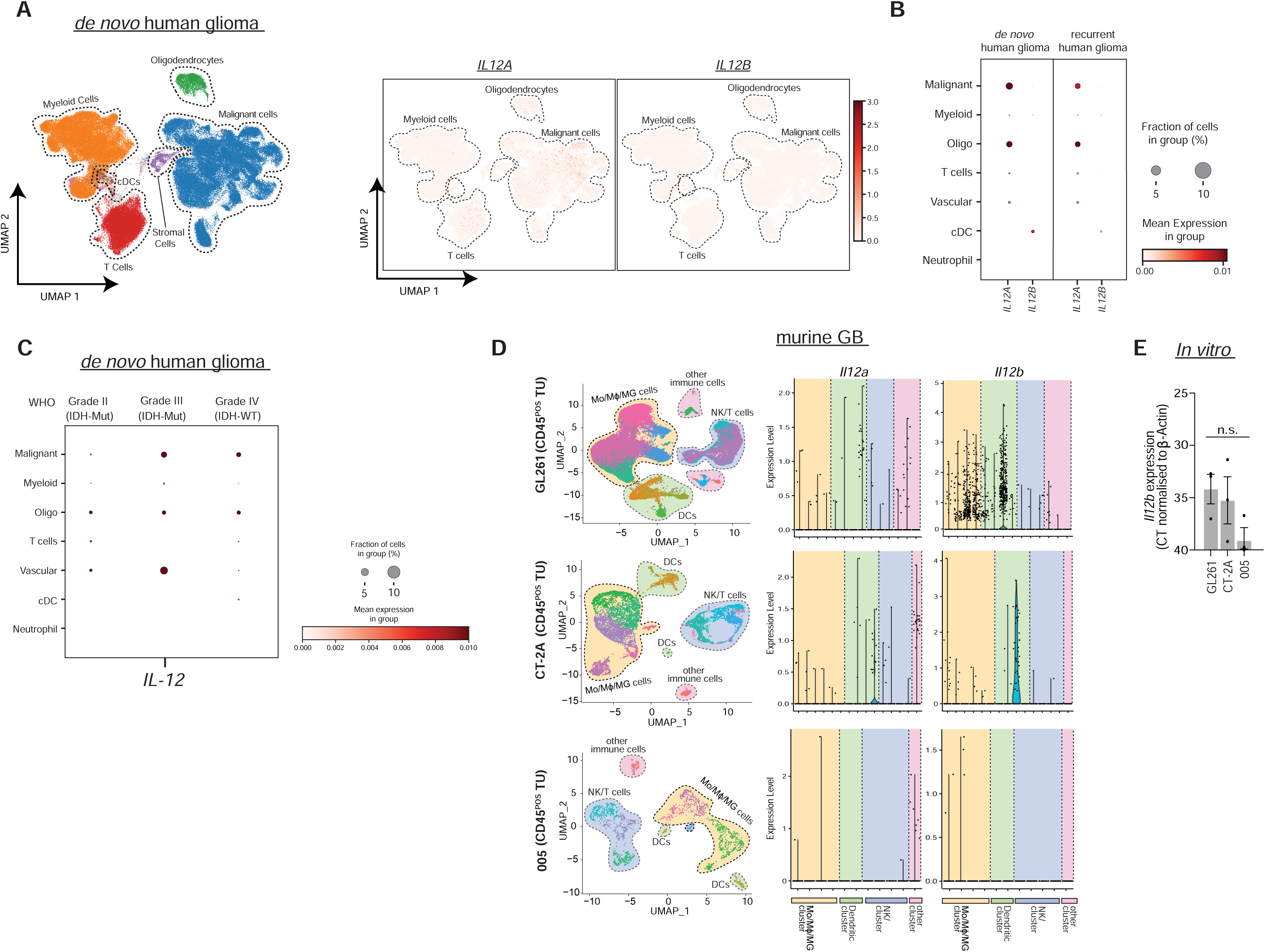

**Supplemental Figure 2:**
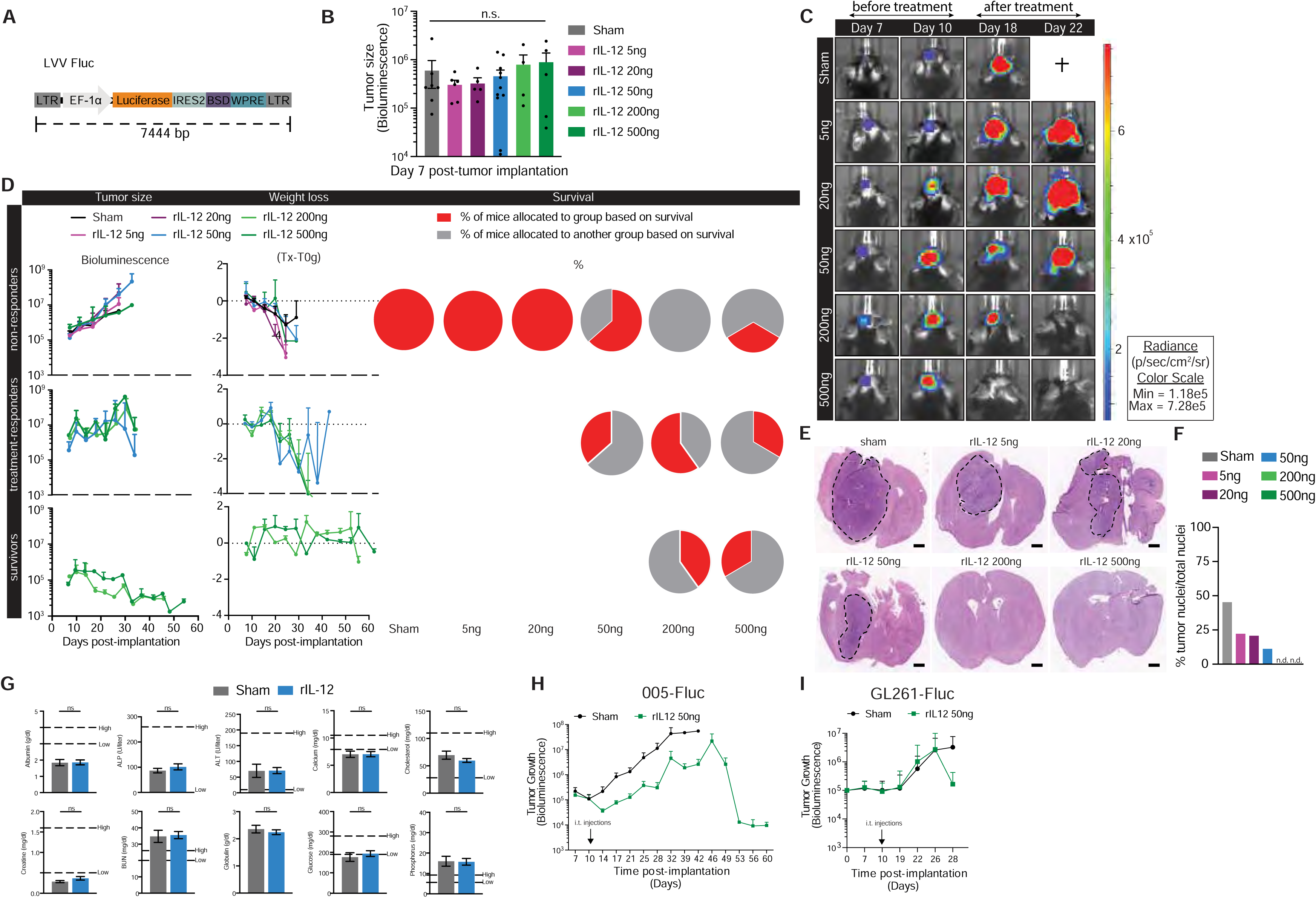

**Supplemental Figure 3:**
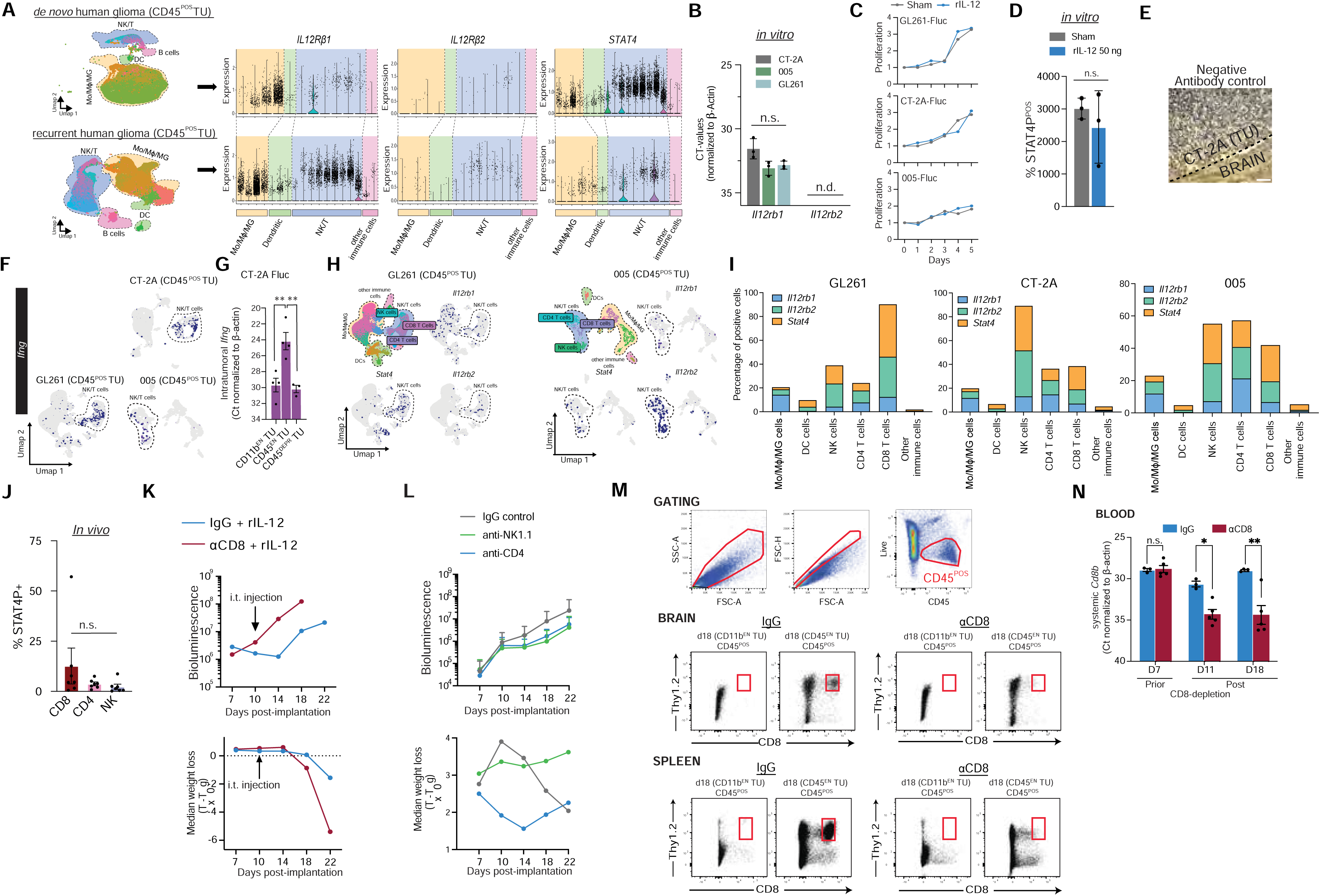

**Supplemental Figure 4:**
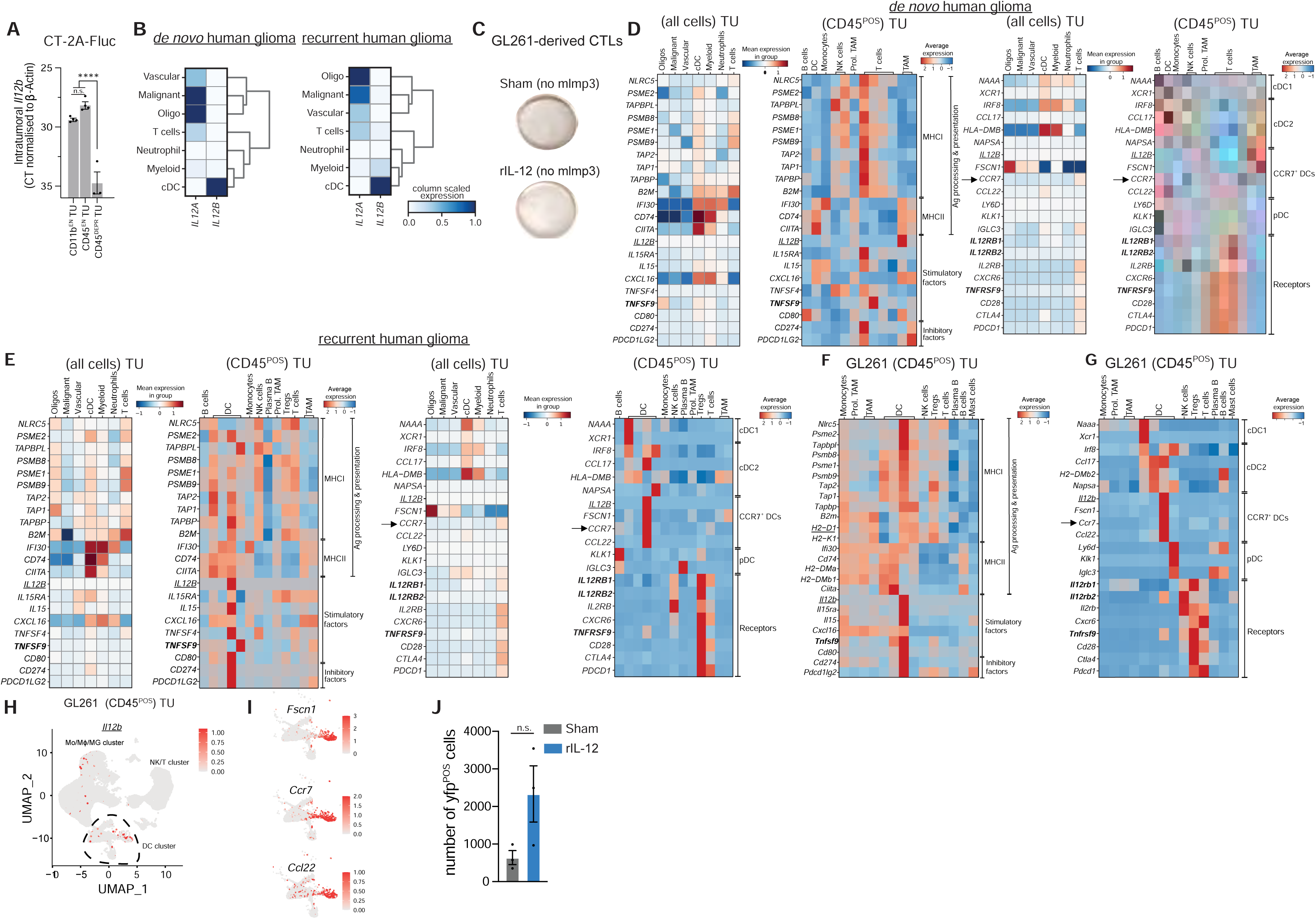

**Supplemental Figure 5:**
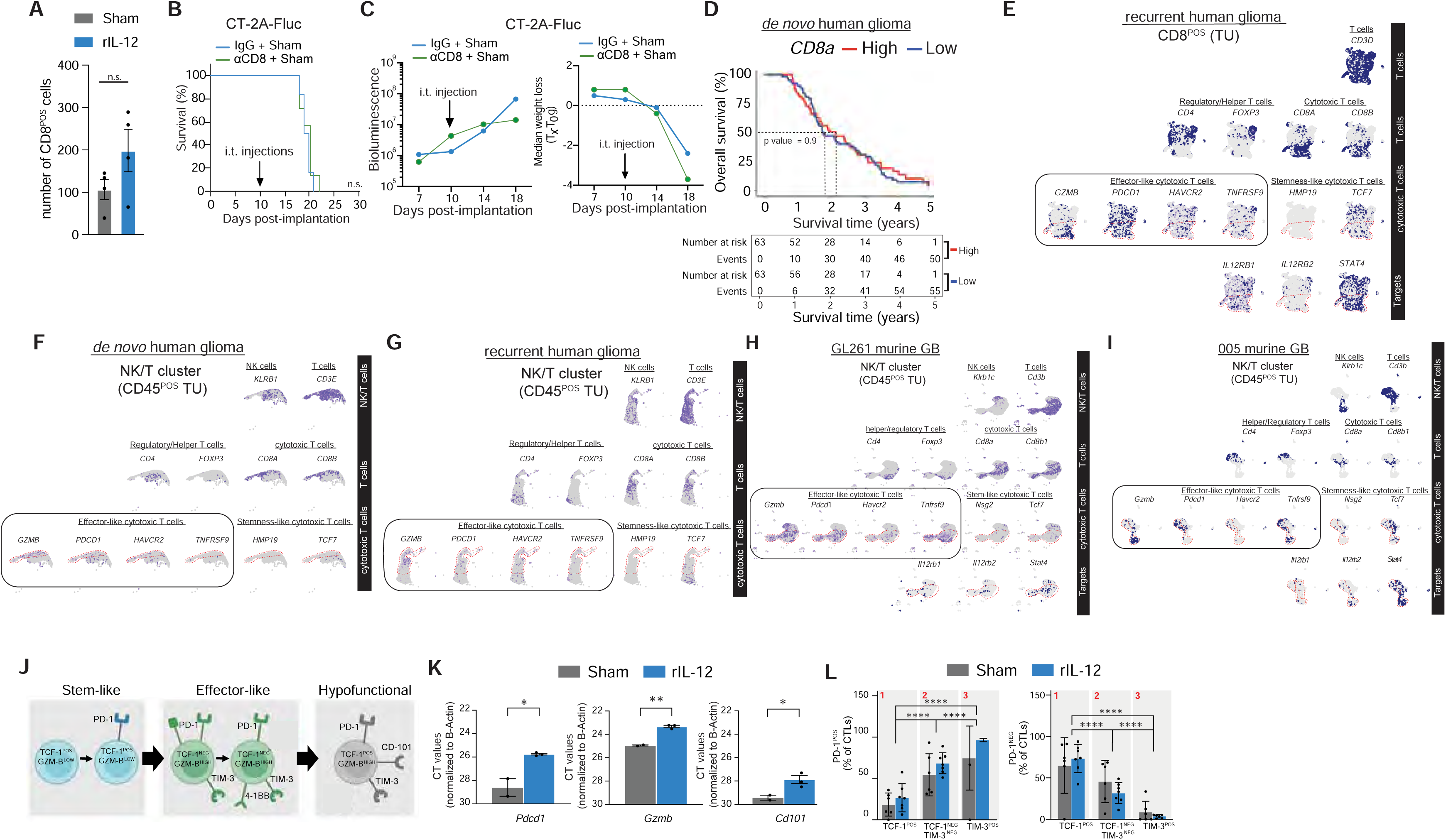

**Supplemental Figure 6:**
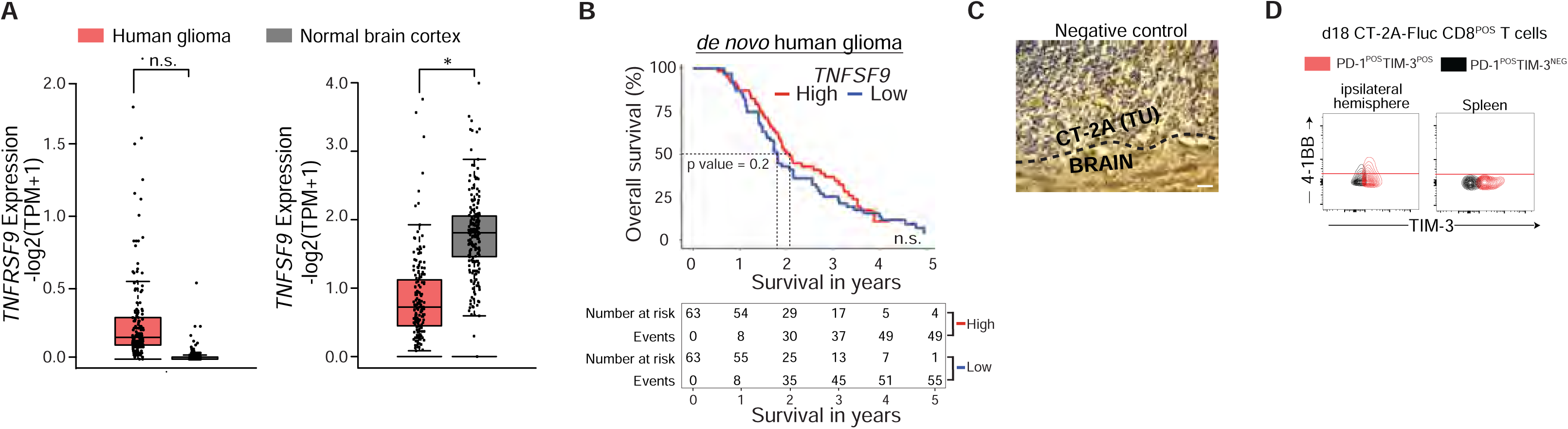

**Supplemental Figure 7:**
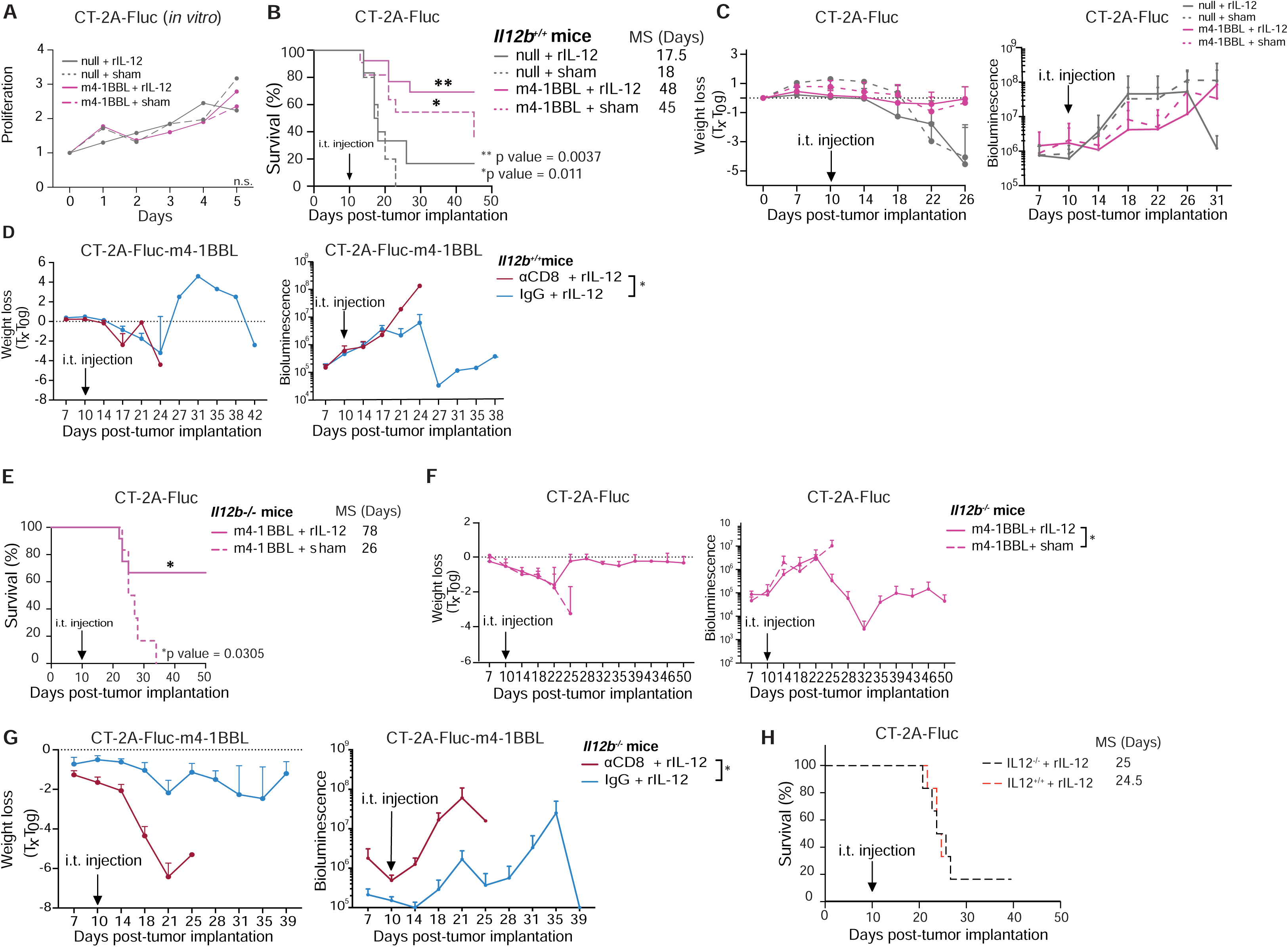

**Supplemental Figure 8:**
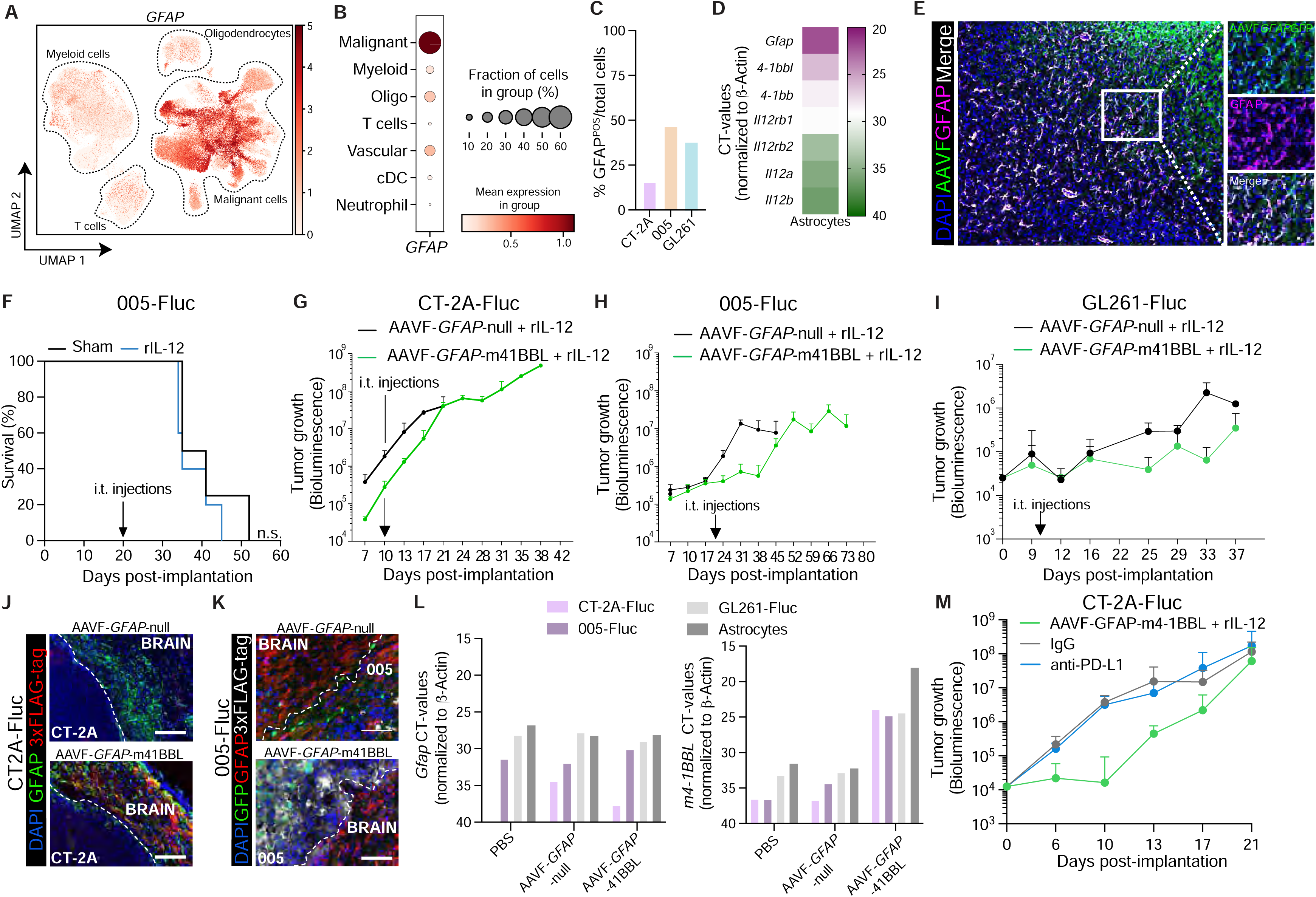

