## Supplementary material for "Intratumoral gene delivery of 4-1BBL boosts IL-12-triggered anti-glioblastoma immunity": Resource Table for Antibodies

**Key resources table**

| REAGENT or RESOURCE | SOURCE | IDENTIFIER |
| --- | --- | --- |
| Antibodies | | |
| Alexa Fluor® 647 anti-mouse TCF1/TCF7 Rabbit mAb (clone C63D9) | Cell Signaling Technology | [Cat# 6709, RRID:AB_2916074](http://antibodyregistry.org/AB_2797627) |
| PE/Cy7 anti-mouse IFN-γ (clone XMG1.2) | BioLegend | [Cat# 505826, RRID:AB_2295770](http://antibodyregistry.org/AB_2295770) |
| BV 605™ anti-mouse CD366 (Tim-3) (clone RMT3-23) | BioLegend | [Cat# 119721, RRID:AB_2616907](http://antibodyregistry.org/AB_2616907) |
| PE anti-mouse CD366 (Tim-3) (clone RMT3-23) | BioLegend | [Cat# 119704, RRID:AB_345378](http://antibodyregistry.org/AB_345378) |
| PerCP/Cy5.5 anti-mouse CD366 (Tim-3) (clone RMT3-23) | BioLegend | [Cat# 119717, RRID:AB_2571934](http://antibodyregistry.org/AB_2571934) |
| Alexa Fluor® 700 anti-mouse CD45 (clone 30-F11) | BioLegend | [Cat# 103128, RRID:AB_493715](http://antibodyregistry.org/AB_493715) |
| BV 421™ anti-mouse CD45 (clone 30-F11) | BioLegend | [Cat# 103134, RRID:AB_2562559](http://antibodyregistry.org/AB_2562559) |
| PE/Cy7 anti-mouse CD45 (clone 30-F11) | BioLegend | [Cat# 103114, RRID:AB_312979](http://antibodyregistry.org/AB_312979) |
| BV 605™ anti-mouse CD45R/B220 (clone RA3-6B2) | BioLegend | [Cat# 103244, RRID:AB_2563312](http://antibodyregistry.org/AB_2563312) |
| Alexa Fluor® 488 anti-mouse CD45.1 (clone A20) | BioLegend | [Cat# 110717, RRID:AB_492863](http://antibodyregistry.org/AB_492863) |
| Alexa Fluor® 647 anti-mouse CD45.1 (clone A20) | BioLegend | [Cat# 110720, RRID:AB_492864](http://antibodyregistry.org/AB_492864) |
| PerCP/Cy5.5 anti-mouse CD45.1 (clone A20) | BioLegend | [Cat# 110727, RRID:AB_893348](http://antibodyregistry.org/AB_893348) |
| BV 510™ anti-mouse CD45.1 (clone A20) | BioLegend | [Cat# 110741, RRID:AB_2563378](http://antibodyregistry.org/AB_2563378) |
| APC/Cy7 anti-mouse CD45.1 (clone A20) | BioLegend | [Cat# 110716, RRID:AB_313505](http://antibodyregistry.org/AB_313505) |
| APC/Cy7 anti-mouse CD45.2 (clone 104) | BioLegend | [Cat# 109824, RRID:AB_830789](http://antibodyregistry.org/AB_830789) |
| APC/Cy7 anti-mouse CD4 (clone GK1.5) | BioLegend | [Cat# 100414, RRID:AB_312699](http://antibodyregistry.org/AB_312699) |
| Alexa Fluor® 700 anti-mouse/human CD11b (clone M1/70) | BioLegend | [Cat# 101222, RRID:AB_493705](http://antibodyregistry.org/AB_493705) |
| PerCP/Cy5.5 anti-mouse CD3 (clone 17A2) | BioLegend | [Cat# 100217, RRID:AB_1595597](http://antibodyregistry.org/AB_1595597) |
| PE/Cy7 anti-mouse CD11c (clone N418) | BioLegend | [Cat# 117318, RRID:AB_493568](http://antibodyregistry.org/AB_493568) |
| Alexa Fluor® 488 anti-rat CD90/mouse CD90.1 (Thy-1.1) (clone OX-7) | BioLegend | [Cat# 202505, RRID:AB_492883](http://antibodyregistry.org/AB_492883) |
| Alexa Fluor® 647 anti-rat CD90/mouse CD90.1 (Thy-1.1) (clone OX-7) | BioLegend | [Cat# 202507, RRID:AB_492885](http://antibodyregistry.org/AB_492885) |
| Alexa Fluor® 700 anti-rat CD90/mouse CD90.1 (Thy-1.1) (clone OX-7) | BioLegend | [Cat# 202528, RRID:AB_1626241](http://antibodyregistry.org/AB_1626241) |
| BV 421™ anti-rat CD90/mouse CD90.1 (Thy-1.1) (clone OX-7) | BioLegend | [Cat# 202529, RRID:AB_10899572](http://antibodyregistry.org/AB_10899572) |
| BV 605™ anti-mouse CD90.2 (clone 30-H12) | BioLegend | [Cat# 105343, RRID:AB_2632889](http://antibodyregistry.org/AB_2632889) |
| BV 510™ anti-mouse CD90.2 (clone 30-H12) | BioLegend | [Cat# 105335, RRID:AB_2566587](http://antibodyregistry.org/AB_2566587) |
| Alexa Fluor® 700 anti-mouse CD90.2 (clone 30-H12) | BioLegend | [Cat# 105320, RRID:AB_493725](http://antibodyregistry.org/AB_493725) |
| PerCP/Cy5.5 anti-mouse CD90.2 (Thy1.2) (clone 53-2.1) | BioLegend | [Cat# 140322, RRID:AB_2562696](http://antibodyregistry.org/AB_2562696) |
| PE/Cy7 anti-mouse CX3CR1 (clone SA011F11) | BioLegend | [Cat# 149015, RRID:AB_2565699](http://antibodyregistry.org/AB_2565699) |
| PE anti-mouse CD185 (CXCR5) (clone L138D7) | BioLegend | [Cat# 145504, RRID:AB_2561968](http://antibodyregistry.org/AB_2561968) |
| BV 421™ anti-mouse CD185 (CXCR5) (clone L138D7) | BioLegend | [Cat# 145512, RRID:AB_2562128](http://antibodyregistry.org/AB_2562128) |
| APC anti-mouse CD191 (CCR1) (clone S15040E) | BioLegend | [Cat# 152503, RRID:AB_2629810](http://antibodyregistry.org/AB_2629810) |
| PE anti-mouse CD192 (CCR2) (clone SA203G11) | BioLegend | [Cat# 150609, RRID:AB_2616981](http://antibodyregistry.org/AB_2616981) |
| BV 421™ anti-mouse CD192 (CCR2) (clone SA203G11) | BioLegend | [Cat# 150605, RRID:AB_2571913](http://antibodyregistry.org/AB_2571913) |
| APC anti-mouse CD196 (CCR6) (clone 29-2L17) | BioLegend | [Cat# 129813, RRID:AB_187714](http://antibodyregistry.org/AB_187714) |
| PE anti-mouse CD195 (CCR5) (clone HM-CCR5) | BioLegend | [Cat# 107006, RRID:AB_313301](http://antibodyregistry.org/AB_313301) |
| APC anti-mouse CD195 (CCR5) (clone HM-CCR5) | BioLegend | [Cat# 107011, RRID:AB_2074528](http://antibodyregistry.org/AB_2074528) |
| PE/Cy7 anti-mouse CD186 (CXCR6) (clone SA051D1) | BioLegend | [Cat# 151119, RRID:AB_2721670](http://antibodyregistry.org/AB_2721670) |
| BV 421™ anti-mouse CD186 (CXCR6) (clone SA051D1) | BioLegend | [Cat# 151109, RRID:AB_2616760](http://antibodyregistry.org/AB_2616760) |
| BV 510™ anti-mouse CD183 (CXCR3) (clone CXCR3-173) | BioLegend | [Cat# 126528, RRID:AB_2650922](http://antibodyregistry.org/AB_2650922) |
| PE/Cy7 anti-mouse CD183 (CXCR3) (clone CXCR3-173) | BioLegend | [Cat# 126515, RRID:AB_2086740](http://antibodyregistry.org/AB_2086740) |
| BV 421™ anti-mouse CD197 (CCR7) (clone 4B12) | BioLegend | [Cat# 120119, RRID:AB_10897811](http://antibodyregistry.org/AB_10897811) |
| Alexa Fluor® 488 anti-mouse CD197 (CCR7) (clone 4B12) | BioLegend | [Cat# 120112, RRID:AB_492842](http://antibodyregistry.org/AB_492842) |
| BV 510™ anti-mouse CD8a (clone 53-6.7) | BioLegend | [Cat# 100751, RRID:AB_2561389](http://antibodyregistry.org/AB_2561389) |
| PerCP/Cy5.5 anti-mouse CD8a (clone 53-6.7) | BioLegend | [Cat# 100734, RRID:AB_2075238](http://antibodyregistry.org/AB_2075238) |
| APC/Cy7 anti-mouse CD8a (clone 53-6.7) | BioLegend | [Cat# 100714, RRID:AB_312753](http://antibodyregistry.org/AB_312753) |
| PerCP/Cy5.5 anti-mouse CD25 (clone PC61) | Biolegend | [Cat# 102030, RRID:AB_893288](http://antibodyregistry.org/AB_893288) |
| APC/Cy7 anti-mouse/rat XCR1 (clone ZET) | BioLegend | [Cat# 148223, RRID:AB_2783117](http://antibodyregistry.org/AB_2783117) |
| PerCP/Cy5.5 anti-mouse CD172a (SIRPa) (clone P84) | BioLegend | [Cat# 144009, RRID:AB_2563547](http://antibodyregistry.org/AB_2563547) |
| APC anti-mouse Ly-6G (clone 1A8) | BioLegend | [Cat# 127613, RRID:AB_1877163](http://antibodyregistry.org/AB_1877163) |
| APC anti-mouse CD335 (NKp46) (clone 29A1.4) | BioLegend | [Cat# 137607, RRID:AB_10612749](http://antibodyregistry.org/AB_10612749) |
| APC anti-mouse Ly-6C (clone HK1.4) | BioLegend | [Cat# 128015, RRID:AB_1732087](http://antibodyregistry.org/AB_1732087) |
| APC anti-mouse CD19 (clone 6D5) | BioLegend | [Cat# 115511, RRID:AB_313646](http://antibodyregistry.org/AB_313646) |
| PE/Cy7 anti-mouse CD69 (clone H1.2F3) | BioLegend | [Cat# 104512, RRID:AB_493564](http://antibodyregistry.org/AB_493564) |
| PerCP/Cy5.5 anti-mouse CD279 (PD-1) (clone RMP1-30) | BioLegend | [Cat# 109119, RRID:AB_2566640](http://antibodyregistry.org/AB_2566640) |
| BV 421™ anti-mouse CD279 (PD-1) (clone RMP1-30) | BioLegend | [Cat# 109121, RRID:AB_2687080](http://antibodyregistry.org/AB_2687080) |
| BV 510™ anti-mouse CD279 (PD-1) (clone 29F.1A12) | BioLegend | [Cat# 135241, RRID:AB_2715761](http://antibodyregistry.org/AB_2715761) |
| APC anti-mouse CD279 (PD-1) (clone 29F.1A12) | BioLegend | [Cat# 135210, RRID:AB_2159183](http://antibodyregistry.org/AB_2159183) |
| PerCP/Cy5.5 anti-mouse CD279 (PD-1) (clone 29F.1A12) | BioLegend | [Cat# 135208, RRID:AB_2159184](http://antibodyregistry.org/AB_2159184) |
| Alexa Fluor® 700 anti-mouse CD8b (Ly-3) (clone YTS156.7.7) | BioLegend | [Cat# 126618, RRID:AB_2563949](http://antibodyregistry.org/AB_2563949) |
| BV 605™ anti-mouse I-A/I-E (clone M5/114.15.2) | BioLegend | [Cat# 107639, RRID:AB_2565894](http://antibodyregistry.org/AB_2565894) |
| BV 421™ anti-mouse CD64 (FcgRI) (clone X54-5/7.1) | BioLegend | [Cat# 139309, RRID:AB_2562694](http://antibodyregistry.org/AB_2562694) |
| PE anti-mouse CD64 (FcgRI) (clone X54-5/7.1) | BioLegend | [Cat# 139304, RRID:AB_10612740](http://antibodyregistry.org/AB_10612740) |
| BV 510™ anti-mouse F4/80 (clone BM8) | BioLegend | [Cat# 123135, RRID:AB_2562622](http://antibodyregistry.org/AB_2562622) |
| PE anti-mouse Ly108 (clone 330-AJ) | BioLegend | [Cat# 134605, RRID:AB_1659258](http://antibodyregistry.org/AB_1659258) |
| APC anti-mouse Ly108 (clone 330-AJ) | BioLegend | [Cat# 134609, RRID:AB_2728154](http://antibodyregistry.org/AB_2728154) |
| PE/Cy7 anti-Bcl-2 antibody (clone BCL/10C4) | BioLegend | [Cat# 633512, RRID:AB_2565247](http://antibodyregistry.org/AB_2565247) |
| Alexa Fluor® 647 anti-Bcl-2 (clone BCL/10C4) | BioLegend | [Cat# 633510, RRID:AB_2274702](http://antibodyregistry.org/AB_2274702) |
| TruStain fcX™ (anti-mouse CD16/32) (clone 93) | BioLegend | [Cat# 101320, RRID:AB_1574975](http://antibodyregistry.org/AB_1574975) |
| BV 785™ anti-mouse CD86 (clone GL-1) | BioLegend | [Cat# 105043, RRID:AB_2566722](http://antibodyregistry.org/AB_2566722) |
| BV 785™ anti-mouse CD274 (B7-H1, PD-L1) (clone 10F.9G2) | BioLegend | [Cat# 124331, RRID:AB_2629659](http://antibodyregistry.org/AB_2629659) |
| Alexa Fluor® 647 anti-mouse Blimp-1 (clone 5E7) | BioLegend | [Cat# 150004, RRID:AB_2565618](http://antibodyregistry.org/AB_2565618) |
| BV 421™ anti-T-bet (clone 4B10) | BioLegend | [Cat# 644815, RRID:AB_10896427](http://antibodyregistry.org/AB_10896427) |
| APC/Cy7 anti-mouse/human KLRG1 (MAFA) (clone 2F1/KLRG1) | BioLegend | [Cat# 138426, RRID:AB_2566554](http://antibodyregistry.org/AB_2566554) |
| BV 605™ anti-mouse CD25 (PC61) | BioLegend | [Cat# 102035, RRID:AB_11126977](http://antibodyregistry.org/AB_11126977) |
| PE anti-mouse CD215 (IL-15Rα) (clone 6B4C88) | BioLegend | [Cat# 153504, RRID:AB_2721342](http://antibodyregistry.org/AB_2721342) |
| Alexa Fluor® 647 anti-human/mouse Granzyme B (clone GB11) | BioLegend | [Cat# 515406, RRID:AB_2566333](http://antibodyregistry.org/AB_2566333) |
| Alexa Fluor® 700 anti-mouse CD3 (clone 17A2) | BioLegend | [Cat# 100215, RRID:AB_493696](http://antibodyregistry.org/AB_493696) |
| Alexa Fluor® 594 anti-mouse CD31 (clone MEC13.3) | BioLegend | [Cat# 102520, RRID:AB_2563319](http://antibodyregistry.org/AB_2563319) |
| PE-Cy7 anti-Foxp3 (clone FJK-16 s), | eBioscience™, Thermo Fisher Scientific | [Cat# 25-5773-82, RRID:AB_891552](http://antibodyregistry.org/AB_891552) |
| PE anti-Foxp3 (clone FJK-16 s) | eBioscience™, Thermo Fisher Scientific | [Cat# 12-5773-82, RRID:AB_465936](http://antibodyregistry.org/AB_465936) |
| Anti-CD3e (clone 145-2C11) | eBioscience, | [Cat# 14-0031-86, RRID:AB_467051](http://antibodyregistry.org/AB_467051) |
|  | Thermo Fisher Scientific |  |
| InVivoMab anti-mouse CD8a (clone YTS 169.4) | Bio X Cell | [Cat# BE0117, RRID:AB_10950145](http://antibodyregistry.org/AB_10950145) |
| PE Rat anti-mouse CXCL16 (clone 12-81) | BD Biosciences | [Cat# 566740, RRID:AB_2869842](https://www.antibodyregistry.org/AB_2869842) |
| PE-Cy7™ hamster anti-mouse CD11c (clone HL3) | BD Biosciences | [Cat# 561022, RRID:AB_2033997](http://antibodyregistry.org/AB_2033997) |
| PE mouse anti-Ki-67 (clone B56) | BD Biosciences | [Cat# 556027, RRID:AB_2266296](http://antibodyregistry.org/AB_2266296) |
| FITC mouse anti-Ki-67 (clone B56) | BD Biosciences | [Cat# 556026, RRID:AB_396302](http://antibodyregistry.org/AB_396302) |
| Alexa Fluor® 647 mouse anti-Ki-67 (clone B56) | BD Biosciences | [Cat# 561126, RRID:AB_10611874](http://antibodyregistry.org/AB_10611874) |
| BV 786™ rat anti-mouse CD103 (clone M290) | BD Biosciences | [Cat# 564322, RRID:AB_2738744](https://www.antibodyregistry.org/AB_2738744) |
| BV 395™ rat anti-mouse CD273 (PD-L2) (clone TY25) | BD Biosciences | [Cat# 565102, RRID:AB_2739068](http://antibodyregistry.org/AB_2739068) |
| BV 395™ hamster anti-mouse CD80 (clone 16-10A1) | BD Biosciences | [Cat# 740246, RRID:AB_2739993](http://antibodyregistry.org/AB_2739993) |
| BV 480™ hamster anti-mouse CD11c (N418) | BD Biosciences | [Cat# 746392, RRID:AB_2743706](http://antibodyregistry.org/AB_2743706) |
| PE Anti-Fascin1 (clone 55K-2) | Santa Cruz Biotechnology | [Cat# sc-21743, RRID:AB_627580](https://www.antibodyregistry.org/AB_627580) |
| Alexa Fluor® 700 anti-mouse IFN-γ Antibody | Biolegend | [Cat# 505823; RRID: AB_2561299](https://www.antibodyregistry.org/AB_2561299) |
| Alexa Fluor® 647 anti-STAT4 Phospho (Tyr693) Antibody | Biolegend | [Cat# 941203, RRID:AB_2936723](https://www.antibodyregistry.org/AB_2936723) |
| PE/Cyanine7 anti-mouse CD274 (B7-H1, PD-L1) | Biolegend | [Cat# 124313, RRID:AB_10639934](https://www.antibodyregistry.org/AB_10639934) |
| Alexa Fluor® 700 anti-mouse CD90.2 (Thy-1.2) Antibody | Biolegend | [Cat# 140324, RRID:AB_2566740](https://www.antibodyregistry.org/AB_2566740) |
| PE/Cyanine5 anti-mouse CD11c Antibody | Biolegend | [Cat# 117316, RRID:AB_493566](https://www.antibodyregistry.org/AB_493566) |
| BUV395 Rat Anti-Mouse CD8a | BD Horizon | [Cat# 565968, RRID:AB_2739421](https://www.antibodyregistry.org/AB_2739421) |
| Pacific Blue™ anti-mouse CD90.2 (Thy-1.2) Antibody | Biolegend | [Cat# 140306, RRID:AB_10641693](https://www.antibodyregistry.org/AB_10641693) |
| BUV395 Rat Anti-Mouse CD90.2 | BD Horizon | [Cat# 565257, RRID:AB_2739136](https://www.antibodyregistry.org/AB_2739136) |
| BUV395 Mouse Anti-Human CD11c | BD Horizon | [Cat# 563787, RRID:AB_2744274](https://www.antibodyregistry.org/AB_2744274) |
| PE anti-mouse 4-1BB Ligand (CD137L) Antibody | Biolegend | [Cat# 107105, RRID:AB_2256408](https://www.antibodyregistry.org/AB_2256408) |
| PE/Cyanine5 anti-mouse CD274 (B7-H1, PD-L1) Antibody | Biolegend | [Cat# 124343, RRID:AB_2894674](https://www.antibodyregistry.org/AB_2894674) |
| APC/Fire™ 750 anti-mouse CD45 Antibody | Biolegend | [Cat# 147714, RRID:AB_2750441](https://www.antibodyregistry.org/AB_2750441) |
