## Supplementary fig legends for "Intratumoral gene delivery of 4-1BBL boosts IL-12-triggered anti-glioblastoma immunity"

**Supplemental Information**

**Supplemental Figure 1.**

(A) *UMAP projections of expression of IL12A and IL12B in de novo human glioma.* The scRNAseq dataset (Miller et al., 2023) of human glioma cells were analyzed. Distinct cell type subsets were clustered, annotated and visualized with a high-resolution color coded UMAP projection. Malignant cells, Myeloid cells, oligodendrocytes, DCs, T cells and stromal cells are depicted (left). *IL12A* and *IL12B* UMAP projections show low expression in each cell cluster (right).

(B) *IL12A and IL12B expression in de novo human glioma and recurrent glioma.* Dot plot showed that the expression of IL12A and IL12B were present at low levels in both primary and recurrent glioma (Miller et al., 2023).

(C) *Expression of IL12A and IL12B at different grade levels of de novo human glioma.* Human *IL12A* and *IL12B* were expressed at low levels in IDH mutant human glioma grade II, III and IV (including WT glioma). No significant difference was observed between the glioma grades.

(D) *UMAP projections of immune cell populations in murine GB.* scRNAseq datasets of murine GB were analyzed. Distinct cell types were clustered, annotated and visualized with a high-resolution color coded UMAP projection. To visualize *IL-12a* and *IL-12b* expression in different cell lines, single cell violin plots were used to compare transcript levels in Mo/Mφ/MG cells (TAM, proliferative TAM and monocytes), dendritic cell cells (DC1, DC2, DC3 and DC4), NK/T cell cells (reg T cells, NK cells), T cells and other cell cluster (B cells, plasma B cells, mast cells). (Chen et al., 2023; Pombo Antunes et al., 2021; Tomaszewski et al., 2022). The data suggested that *Il12b* was expressed at minimal levels in NK/T and other cell clusters

(E) *Murine GB cell lines express minimal levels of Il12b.* GL261, CT-2A and 005 cells show low expression levels of *Il12b.* Data represent three independent experiments and are presented as the mean with SEM (error bars). Data were analyzed using one way ANOVA, not significant (n.s.).

**Supplemental Figure 2.**

(A) *Lentiviral construct.* Schematic display of LVV-Firefly Luciferase (FLuc) containing luciferase, IRES2, BSD and WPRE driven by the EF-1α promoter.

(B) *Bioluminescence measurements.* To ensure equal tumor size between groups, CT-2A-FLuc bioluminescence levels were measured in GB-bearing mice at day 7 post-tumor implantation, prior to i.t. treatment of sham or rIL-12. No significant differences were observed between groups (50 ng sham, n= 7; 5 ng rIL-12, n = 6; 20 ng rIL-12, n= 5; 50 ng rIL-12, n= 11; 200 ng rIL-12, n=5, and 500 ng rIL-12, n=6). Multiple t-test, not significant (n.s.)

(C) *Regression of tumors in GB-bearing mice with rIL-12 over time.* FLuc readings demonstrate that tumors were established in mice brains before treatment on days 7 and 10. Different outcomes were observed after treatment on days 18 and 22 post tumor-implantation depending on the rIL-12 dose that GB-bearing mice received. The shown images are representative in vivo imaging system (IVIS) images of GB-bearing mice from each treatment condition and “+” indicates that animals died prior to the imaging timepoint.

(D) *Classification of rIL-12 treated GB-bearing mice based on multiple parameters.* Based on tumor growth and weight loss, three categories could be distinguished. Non-responders (n=27) represent mice that had similar results as the sham treatment. Treatment-responders (n=9) represent mice that performed better than the sham-treated mice, but still died. Survivors (n=4) represent mice that had visible tumor regression due to treatment. In the left graphs, CT-2A-FLuc-bearing mice treated rIL-12 or sham were monitored every 3-4 days by IVIS imaging, which is representative of the tumor size in the brain. Dotted line represents the background signal. In the middle graphs, the weight of the mice was tracked over time. The dotted line represents the weight at start of the experiment (T_0_), while Tx denotes specific time points at which weights were measured. The pie graphs on right represent the percentage of mice that are allocated to a certain category based on survival. Data represents two independent experiments and are presented as the mean with SEM (error bars).

(E) *Tumor sizes in brains of rIL-12 treated GB-bearing mice*. Brain sections of mice 22 days post-i.c. implantation with CT-2A-FLuc and i.t. treatment with sham or rIL-12 were stained for hematoxylin and eosin (H&E) (4x magnification, scale bar = 5 µm). The black dotted line indicates the tumor border.

(F) *Hematoxylin & Eosin.* Quantification of H&E histology images shown as the percentage of tumor nuclei over the total number of nuclei. Images were analyzed by Image J using the color deconvolution plugin (n=1 per group; n.d. – not determined).

(G) *Blood chemistry analysis.* Whole blood from GB bearing mice treated with 50 ng rIL-12 or sham showed no significant differences for systemic toxicity at post-tumor implantation. Albumin, Alkaline Phosphatase (ALP), Alanine transaminase (ALT), calcium, cholesterol, creatine, blood urea nitrogen (BUN), globulin, glucose and phosphorus were determined. Data represents three independent experiments and are presented as the mean with SEM (error bars). Data were analyzed using unpaired t-test, not significant (n.s.).

(H&I) Average bioluminescence intensity (BLI) levels of 005-FLuc and GL261-FLuc tumor-bearing mice were measured over time comparing sham (solid black) and rIL-12 (solid green) (n=4-5 mice per group).

**Supplemental Figure 3.**

(A) *Expression of IL12Rβ1, IL12Rβ2 and STAT4 in de novo and recurrent human glioma in immune cell populations*. scRNAseq cluster datasets of CD45^POS^-sorted cells derived from *de novo* human glioma tumors (n=7) and human recurrent glioma tumors (n=4) were analyzed (left). Distinct cell type subsets were clustered, annotated and visualized with a high-resolution color coded UMAP projection. To visualize IL12Rβ1, IL12Rβ2 and STAT4 expression in different datasets, single cell violin plots were used to compare transcript levels in Mo/Mϕ/MG cells (TAM, proliferative, TAM and monocytes), dendritic cells (DC1, DC2, DC3 and DC4), NK/T cells (reg T cells, NK cells, T cells) and other cell cluster (B cells, plasma B cells, mast cells). (Chen et al., 2023; Pombo Antunes et al., 2021; Tomaszewski et al., 2022).

(B) *Expression of Il12rb1/2 in mouse cell lines.* The expression of *Il12rb1* was determined in three different mouse glioma cell lines (CT-2A, 005 and GL261) using qRT-PCR. The expression of *Il12rb1* did not show any significant difference, whereas Il12rb2 was not determined (n.d.) in any of the cell lines. Data represent three independent experiments and were analyzed using two-way ANOVA multiple comparison test, not significant (n.s.).

(C) *Cell viability assay.* CT-2A-FLuc, 005-FLuc and GL261-FLuc cell lines were exposed to 50 ng rIL-12 or sham control and cell viability (proliferation) was measured over a period of 5 days. Data represents triplicates. No significant differences were observed between groups, Unpaired t-test.

(D) *Quantification of STAT4P protein expression in CT-2A-FLuc cultured cells.* CT-2A-FLuc cells were exposed to sham or rIL-12 for 24 hours, STAT4P protein levels were quantified by flowcytometry. Data represent three independent experiments and are presented as the mean with SEM (error bars). Data was analyzed using paired t-test, not significant (n.s.).

(E) *Primary antibody (IL12rb1) control staining in brain tissues implanted with a CT-2A tumor.* Immunohistochemistry negative control lacking primary antibody (IL12rb1) in the CT-2A tumor (TU) cells in the brain TME (magnification 40x; scale bar = 100 µm).

(F) *Expression of Infy in immune cells populations of mouse GB cell lines*. scRNAseq cluster datasets of CD45^POS^-sorted tumor cells derived from GL261 (Pombo Antunes et al., 2021), CT-2A (Tomaszewski et al., 2022) and 005 (Chen et al., 2023) cells were analyzed. Distinct cell type subsets were clustered, annotated and visualized with a coded UMAP projection and NK/T cells were highlighted with a dotted line.

(G) *Expression of Ifn-γ in CT-2A-FLuc implanted mice.* TH brain tissue was analyzed eight days after rIL12-treatement. *Ifn-γ* was expressed significantly higher in CD45^EN^ TU compared to CD11b^EN^ TU (p value = 0,0075) and CD45DEPR TU cells (p value = 0.0067). Data represent CT values normalized to β-actin. Data represent three independent experiments and are presented as the mean with SEM (error bars), Data were analyzed using one-way ANOVA, **p < 0.01.

(H) *Expression of Il12rb1, Il12rb2 and Stat4 in GL261 and 005 murine GB cell lines*. scRNAseq cluster datasets of CD45^POS^-sorted cells derived from GL261 (n=3) (Pombo Antunes et al., 2021), and 005 (n=4) (Chen et al., 2023) tumor-bearing C57BL6 mice were analyzed (left). Distinct cell type subsets were clustered, annotated and visualized with a high-resolution color coded UMAP projection. To visualize Il12rb1, Il12rb2 and Stat4 expression in different datasets, single cell feature plots were used to compare transcript levels in NK/T cells cluster (reg T cells, NK cells, T cells) as demonstrated by dotted lines.

(I) *Percentages of positive cells in each immune cell cluster is shown by bar graphs.* Datasets acquired (**Table 1**) were analyzed for *Il12rb1, Il12rb2* and *Stat4* expression in GL261, CT-2A and 005 GB cells, including Mo/Mϕ/MG cells, dendritic cells, natural killer cells, Regulatory T cells, T cells, and other immune cells clusters. (Chen et al., 2023; Pombo Antunes et al., 2021; Tomaszewski et al., 2022).

(J) *Quantification of STAT4p percentages in CD8, CD4 T cells and NK1.1 cells.* Bar graphs represent quantification of the percentages of STAT4p positive cells within CD8/CD4/NK cells (n=6 mouse/group). CD8 T cells showed the highest percentage of STAT4p (12.6%), followed by CD4 T cells (3.7%) and NK1.1 cells (2.3%). Data represent three independent experiments and are presented as the mean with SEM (error bars). Data was analyzed using multiple comparison one-way ANOVA, not significant (n.s.).

(K) Tumor growth and weight were measured over time in tumor-bearing mice injected with IgG and rIL-12 (solid blue), anti-CD8 and rIL-12 (solid red). After T-cell depletion, mice had increased tumor sizes as measured by BLI. Weights of all mice dropped starting day 14 after tumor cell injection. (n = 6-8 mice per group).

(L) Tumor growth and mice weight were measured over time in tumor-bearing mice injected with anti-CD4 or anti-NK1.1 compared to IgG control all treated i.t. with rIL-12 at day 10 post-tumor implantation. BLI signal did not differ between groups. Weights of NK1.1 depleted mice increased over time whereas CD4-depleted mice had reduced weights over time. (n = 6-8 mice per group).

(M) Validation of CD8^POS^ T cell depletion shown by the absence of CD8^POS^ cells in representative flow cytometry plots showing cell fractions for CD11b^EN^ TU or CD45^EN^ TU, pre-gated for CD45^EN^ cells, after treatment with anti-CD8 compared to IgG control for brain and spleen samples.

(N) *Systemic CD8b expression levels measured over time.* Gene expression levels in retro-orbital blood samples obtained from CD8-depleted mice (n=5) show a significant reduction in *CD8b* at day 11 and 18 (*p = 0.014, **p = 0.0013, respectively), post CD8-depletion compared to IgG control mice (n=3); no significant drop of CD8b was observed at day 7 (prior to CD8-depletion). Data represent two independent experiments and are presented as the mean with SEM (error bars). Data was analyzed using two-way ANOVA, *p<0.05, **p<0.01, not significant (n.s.).

**Supplemental Figure 4.**

(A) Intratumoral *Il12b* expression in CT-2A-FLuc cell populations. *Il12b* was predominantly expressed by CD11b^EN^ TU and CD45^EN^ TU cells compared to CD45^DEPR^ TU cells. Data represent three independent experiments and are presented as the mean with SEM (error bars). Data were analyzed using one way ANOVA, ****p < 0.0001, not significant (n.s.).

(B) *IL12A and IL12B expression in de novo human glioma and recurrent glioma.* Using hierarchical clustering, *IL12B* expression was found to be highest in the dendritic cell (DC) cluster across both primary and recurrent glioma samples (Miller et al., 2023) .

(C) *Representative images of* IFN-*γ Elispot assay.* Control groups showing minimal number of IFN-*γ* positive spots when CD8^POS^ T cells and naïve splenocytes were co-cultured without mlmp3 (GL261-specific peptide), comparing rIL-12 and sham control.

(D) Heatmaps of *de novo* human glioma showing regulatory factors of interest for CTL differentiation by CCR7^POS^ DC cluster genes identified in T cells, Neutrophils, Myeloid, cDCs, Vascular, Malignant and Oligodendrocytes cell clusters of all cells (TU) (Miller et al., 2023). B cells, DCs, Monocytes, NK cells, Proliferative tumor-associated macrophages (TAMs), T cells and TAMs clusters post-CD45 (TU) enrichment (Pombo Antunes et al., 2021). The arrow points at the CCR7 gene.

(E) Heatmaps of recurrent human glioma showing regulatory factors of interest for CTL differentiation by CCR7^POS^ DC cluster genes identified in T cells, Neutrophils, Myeloid, cDCs, Vascular, Malignant and Oligodendrocytes cell clusters of all cells (TU) (Miller et al., 2023). B cells, DCs, Monocytes, NK cells, Plasma B, Proliferation tumor-associated macrophages (TAMs), Regulatory T cells (Tregs), TAMs post-CD45 (TU) enrichment (Pombo Antunes et al., 2021). The arrow points at the CCR7 gene.

(F & G) Heatmaps showing co-expression of CCR7^POS^ DC cluster genes identified in Mo/Mϕ/MG, dendritic and NK/T cells clusters and other cell clusters in GL261 CD45^POS^ tumor cells (Pombo Antunes et al., 2021). *H2-D1* and *IL12b* are highlighted. The arrow points at the CCR7 gene.

(H) *IL-12 expression profile matches CCR-7^POS^-DCs*. IL12b was highly expressed in subcluster CCR-7^POS^-DCs which had marked expression of *Fsn1, Ccr7* and *Ccl22* in GL261 (CD45^POS^) TU (Pombo Antunes et al., 2021).

(I) *Visualization of CCR-7^POS^-DCs in scRNAseq dataset*. The cells positive in panel H (marked with dotted line) match with the dendritic specific markers *Fscn1, Ccr7,* and *Ccl22* in GL261 (CD45^POS^) TU (Pombo Antunes et al., 2021).

(J) *Quantification of IHC images showing the number of yfp^POS^ cells.* The number of IL-12b^YFP^ to label CCR7^POS^ DCs were quantified at the tumor border (CT-2A-FLuc) using Image J comparing sham and rIL-12 treated mice (n=3 per group). Although a higher number of yfp^POS^ cells was observed post rIL-12 therapy, no significant differences between groups were observed. Data are presented as the mean with SEM (error bars). Data were analyzed using Unpaired t-test, not significant (n.s.)

**Supplemental Figure 5.**

(A) *Quantification of the number of CD8^POS^ cells as determined by IHC.* The number of CD8^POS^ T cells were quantified at the tumor border (CT-2A-FLuc) using Image J comparing sham and rIL-12 treated mice. Although a higher number of CD8^POS^ cells was observed post rIL-12 therapy, no significant differences between groups were observed. Data are presented as the mean with SEM (error bars). Data were analyzed using Unpaired t-test, not significant (n.s.)

(B) *Importance of CD8 depletion for survival benefit in anti-GB therapy.* Kaplan-Meier curves showing survival outcome of tumor-bearing mice injected without CD8-depletion (IgG) (solid blue) and with CD8 depletion (anti-CD8b) (solid green), (n = 6-8 mice/group). Both groups had a median survival of 20 days. Log-rank (Mantel-Cox) test, not significant (n.s)

(C) Tumor growth and mice weight were measured over time in tumor-bearing mice injected with IgG and sham, anti-CD8 and sham. After T-cell depletion, mice had increased tumor sizes as measured by BLI. Weights of all mice dropped starting day 14 after tumor cell injection.

(D) *Survival probability of de novo human glioma differentiating high and low CD8a.* Kaplan-Meier survival curves showing the survival outcomes over a period of 5 years of 63 glioma patients per group with high (red) or low (blue) levels of *CD8a*, each group had a median of ~2 years (Miller et al., 2023). No differences were observed between groups. Log-rank (Mantel-Cox) test, p-value = 0.9, not significant (n.s).

(E, F & G) Differentiating stem- and effector-like CTLs at human primary and recurrent glioma tumor site. scRNAseq analysis distinguishes NK cells from T cells based on *KLRB1* and *CD3E* expression, respectively. Then, helper and regulatory T cells, expressing *CD4* and *FOXP3* genes, were discriminated from the CTLs marked by *CD8A/B* expression. In the CTL cluster, we observed naïve and stem-like CTLs, expressing *TCF7* (encoding TCF-1) and *HMP19* genes, that were different from the CTLs with an effector-like phenotype expressing *HAVCR2* (encoding TIM-3), *PDCD1* (encoding PD-1), *GZM-B* (encoding cytotoxic granzyme-B) and *TNFRSF9* (encoding 4-1BB) in *de novo* and recurrent human glioma (Mathewson et al., 2021). The targets IL12RB1, IL12RB2 and STAT4 expression in CTLs were also identified based on the marked red dotted border.

(H & I) *Differentiating stem- and effector-like CTLs in GL261 and 005 murine GB*. scRNAseq analysis GL261 (Pombo Antunes et al., 2021), and 005 (Chen et al., 2023) distinguishes NK cells from T cells based on *Klrb1c* and *Cd3b* expression, respectively. Then, helper and regulatory T cells, expressing *Cd4* and *Foxp3* genes, were discriminated from the CTLs by marked *Cd8a/b* expression. In the CTL cluster, we observed naïve and stem-like CTLs, expressing *Tcf7* (encoding TCF-1) and *Nsg2* genes, that were different from the CTLs with an effector-like phenotype expressing *Havcr2* (encoding TIM-3), *Pdcd1* (encoding for PD-1), *Gzmb* (encoding for cytotoxic granzyme-B) and *Tnfrsf9* (encoding for 4-1BB) in GL261 GB and 005 GB, respectively. The targets Il12rb1, Il12rb2 and Stat4 expression in CTLs were also identified based on the marked red dotted border.

(J) *Schematic overview to illustrate the stages of T cell differentiation*. Stem-like CTLs (TCF-1^POS^, GZM-B^LOW^) gain PD-1, effector-like CTLs (TCF-1^NEG^, GZM-B^HIGH^) express PD-1, TIM-3 and 4-1BB, and hypofunctional (TCF-1^POS^, GZM-B^HIGH^) CTLs express CD101.

(K) Gene expression levels showed that *Pdcd1* (the PD-1 gene – average CT values 25.7 sham; 28.6 rIL-12), *Gzmb* (average CT values 23.3 sham; 24.9 rIL-12), and *Cd101* (average CT values 27.9 sham; 29.4 rIL-12), transcripts were increased with 5.6-, 3.0- and 2.9-fold, respectively rIL-12 treated (blue) compared to Fc control (grey) measured in RNA from total mouse brain. Data represent three independent experiments and are presented as the mean with SEM (error bars). Data were analyzed using paired student t-test, *p < 0.05, **p < 0.01.

(L) Quantification of flow cytometry PD-1^POS^ and PD-1^NEG^ comparing rIL-12 (blue) and sham control (grey) within TCF-1^POS^ (panel 1), TCF-1^NEG^TIM-3^NEG^ (panel 2) and TIM-3^POS^ (panel 3) cell populations. TIM-3^POS^ cells expressed 4-fold higher PD-1 levels compared to TCF-1^POS^ and 2-fold increased PD-1 levels compared to TIM-3^NEG^TCF-1^NEG^ cells. TIM-3^POS^ cells expressed significantly less PD-1 compared to TCF-1^POS^ and TIM-3^NEG^TCF-1^NEG^ cells. Data represent two independent experiments and are presented as the mean with SEM (error bars). Data were analyzed using two-way ANOVA, ****p < 0.0001.

**Supplemental Figure 6.**

(A) *Expression of TNFRSF9 and TNFSF9 in glioma patients.* Box plots showing the expression of *TNFRSF9* (4-1BB) and *TNFSF9* (4-1BBL) gene in glioma patients (n = 163) compared to tissue of normal brain cortex (n = 207). The plot showed significant low expression of *TNFSF9* in glioma tissue as compared to normal brain. The data was generated from GEPIA 2.0 (<http://gepia2.cancer-pku.cn/#index>) (Tang et al., 2019) using the TCGA portal. *p < 0.05, not significant (n.s.).

(B) *Survival probability of de novo human glioma differentiating high and low expressing TNFSF9.* Kaplan-Meier survival curves showing the overall survival outcomes over a period of 5 years of a total of 63 glioma patients per group with high (red) or low (low) levels of *TNFSF9*, both groups had a median of ~2 years (Miller et al., 2023). No differences were observed between groups. Log-rank (Mantel-Cox) test, p-value > 0.9, not significant (n.s).

(C) *Primary antibody (4-1BB) control staining in brain tissues implanted with a CT-2A tumor.* Immunohistochemistry lacking primary antibody (4-1BB) in the CT-2A tumor (TU) cells in the brain TME (magnification 40x; scale bar = 100 µm).

(D) *Representative flow cytometry counter plots.* Overlaid counter plots of 4-1BB expression within the PD-1^POS^ TIM-3^POS^ and PD-1^POS^ TIM-3^NEG^ populations showed the increased presence of 4-1BB receptor in the ipsilateral hemisphere compared to the spleen.

**Supplemental Figure 7.**

(A) *Cell viability assay.* CT-2A-FLuc-null and CT-2A-FLuc-m4-1BBL cell lines were exposed to 50 ng rIL-12 or sham control and cell viability (proliferation) was measured over a period of 5 days. Data represents triplicates and no significant differences were observed between groups, Unpaired t-test, not significant (n.s.).

(B) *Survival benefit of local m4-1BBL expression in CT-2-FLuc-bearing mice post-rIL-12 treatment*. Kaplan-Meier curves showing survival outcomes following treatment of CT-2A-FLuc-null with sham (dashed grey) or CT-2A-FLuc-m4-1BBL treated with sham (dashed pink) and CT-2A-FLuc-null with rIL-12 (solid grey) or CT-2A-FLuc-m4-1BBL treated with rIL-12 (solid pink) (n= 4-5 mice per group). Mice injected with CT-2A-FLuc-m4-1BBL tumor cells treated with rIL-12 (50 ng) had a median survival of 48 days (p-value = 0.0431), mice treated with sham control had a median survival of 45 days. Compared to mice implanted with tumor cells lacking m4-1BBL treated with rI-12 or sham had a median survival of 17.5 or 18 days, respectively. Data represent at least two independent experiments and are presented as the mean with ± SEM (error bars). Data were analyzed using the Log-rank (Mantel-Cox) test *p < 0.05, **p < 0.01.

(C) Weight (left) and tumor growth (right) were measured over time in *Il12^+/+^* mice injected with CT-2A-FLuc-null or CT-2A-FLuc-m4-1BBL tumor cells and treated with rIL-12 (solid red) or sham (dashed red). Mice treated with Fc control showed a weight drop at day 22, while rIL-12 treated mice maintained their weight over 50 days. Mice treated with rIL-12 showed a decrease in tumor size starting at day 22, with increasing size in Fc treated mice to day 28 - time of death (n=9-12 mice per group).
(D) Weight (left) and tumor growth (right) were measured over time in tumor-bearing *Il12b^+/+^* mice injected with IgG and rIL-12 (solid blue) or anti-CD8 and rIL-12 (solid red). After T-cell depletion, weights of mice dropped at day 21 and these mice had significantly increased tumor sizes compared to IgG control. (n=5-6 mice per group).

(E) *GB mouse survival upon m4-1BBL and rIL-12 combination treatment is not dependent on endogenous IL-12.* Kaplan-Meier curves of *Il12^-/-^* mice showing survival outcome of CT-2A-FLuc-m4-1BBL tumor-bearing mice injected intratumorally with rIL-12 (solid pink), or the sham (dashed pink). Mice (n= 9-12 mice per genotype) treated with rIL-12 had a median survival of 78 days compared to 26 days for sham treated. Data represent at least two independent experiments and are presented as the mean with ± SEM (error bars). Data were analyzed using the Log-rank (Mantel-Cox) test *p < 0.05.

(F) Weight (left) and tumor growth (right) were measured over time in WT mice injected with CT-2A-FLuc-control (grey) or CT-2A-FLuc-m4-1BBL (red) tumor cells comparing rIL-12 (solid) to sham control (dashed) treatment (n = 5-11 mice per group).

(G) Weight (left) and tumor growth (right) were measured over time in tumor-bearing *Il12b^-/-^* mice injected with IgG and rIL-12 (solid blue) or anti-CD8 and rIL-12 (solid red). After T-cell depletion, weights of mice dropped at day 14 and these mice had significantly increased tumor sizes compared to IgG control. (n=5-6 mice per group). Data represents at least two independent experiments and were analyzed using multiple t-test, *p < 0.05.

(H) *Survival curves of GB-bearing mice in Il12b^+/+^ and Il12b^-/-^ mice treated with rIL-12.* Kaplan-Meier survival curves showing no overall survival benefit of CT-2A tumor-bearing *Il12^+/+^* mice (red) and *Il12^-/-^* mice (blue) (n = 5-6 mice per group) (median survival of 24.5 days and 25 days, respectively). Data represents at least two independent experiments. No differences were observed between the groups. Log-rank (Mantel-Cox) test, not significant (n.s).

**Supplemental Figure 8.**

(A) *UMAP projections of expression of GFAP in human glioma.* The scRNAseq dataset (Miller et al., 2023) of human glioma cells were analyzed. Distinct cell type subsets were clustered, annotated and visualized with a high-resolution color coded UMAP projection. *GFAP* was most expressed in the malignant cluster, compared to the oligodendrocytes, DCs, T cells and stromal clusters.

(B) *Expression of GFAP in human glioma.* Human *GFAP* was expressed at high levels in malignant cluster as depicted in a dot plot (Miller et al., 2023).

(C) Quantification of the number of GFAP^POS^ shown as a percentage of total cells as stained for by DAPI comparing CT-2A, 005 and GL61 GB models. Images were analyzed by Image J (n=1 per group).

(D) Gene expression levels shown for *Gfap*, *Il12a*, *Il12b*, *Il12rb1*, *Il12rb2* *41bb* and *41bbl* mRNA measured in primary mouse astrocytes. Data are plotted as CT values normalized to β-actin and displayed as a heatmap.

(E) Immunofluorescence of a brain without tumor i.c. injected at day 7 with AAVF-*GFAP*-GFP backbone vector showed successful targeting of GFAP astrocytes after 14 days post-injection. (10x magnification, scale bar = 10 µm, left; 40x magnification, scale bar = 50 µm, right).

(F) *Survival benefit is reduced by delayed treatment of 005-FLuc-bearing mice with rIL-12.* Kaplan-Meier curves displaying the percentage of survival of 005-FLuc-bearing mice (100,000 cells at the time of injection) with treatment at day 20 post-tumor implantation, comparing i.c. injection of 50 ng rIL-12 (blue) to sham control (black) (n=4-5 mice per group). No significant difference was observed for rIL-12 treated mice with a median survival of 35 days compared to sham with a median survival of 38 days.

(G) Average bioluminescence levels of CT-2A-FLuc tumor-bearing mice were measured over time comparing AAVF-*GFAP*-null treated with rIL-12; (solid black) and AAVF-*GFAP*-m4-1BBL treated with rIL-12 (solid green) (n=4-6 mice per group).

(H) Average bioluminescence levels of 005-FLuc tumor-bearing mice were measured over time comparing AAVF-*GFAP*-null treated with rIL-12; (solid black) and AAVF-*GFAP*-m4-1BBL treated with rIL-12 (solid green) (n=4-5 mice per group).

(I) Average bioluminescence levels of GL261-FLuc tumor-bearing mice were measured over time comparing AAVF-*GFAP*-null treated with rIL-12; (solid black) and AAVF-*GFAP*-m4-1BBL treated with rIL-12 (solid green) (n=4-5 mice per group).

(J) Immunofluorescence of brain sections from mice implanted with CT-2A tumor (12,500 cells) and i.t. injected with AAVF-GFAP null and m-4-1BBL vector (three times) showed successful targeting of GFAP astrocytes and 3x FLAG-tag (red) in TME in the tumor vicinity. The white dotted line represents the tumor border. (40x magnification, scale bar = 50 µm).

(K) Immunofluorescence of brain sections from mice implanted with 005-GFP tumor (50,000 cells) and i.t. injected with AAVF-GFAP null and m-4-1BBL vector (three times) showed successful targeting of GFAP astrocytes (red) and 3xFLAG-tag (white) in TME in the tumor vicinity. The white dotted line represents the tumor border. (40x magnification, scale bar = 50 µm).

(L) CT-2A cells (light purple), 005 cells (dark purple), GL261 (light grey) and primary derived astrocytes (grey) were transduced with AAVF-*GFAP*-m4-1BBL or AAVF-*GFAP*-null control and maintained for 7 days in culture. mRNA levels showed increased *Gfap* expression in astrocytes compared to CT-2A and 005 cells; all four cell types showed increased levels of the *m4-1BBL* transgene only after incubating with AAVF-*GFAP*-m4-1BBL, and not with the AAVF-*GFAP*-null control, compared to PBS control. Data are plotted as CT values normalized to β-actin. Data represents one independent experiment.

(M) Average bioluminescence levels of CT-2A-FLuc tumor-bearing mice were measured over time comparing AAVF-*GFAP*-m4-1BBL treated with rIL-12; (solid green); anti-PD-L1 treated with rIL-12 (solid blue) and IgG control (solid grey) treated with rIL-12 (n = 4-5 mice per group). Multiple t-test, not significant (n.s.).

**Tables**

Table 1. scRNAseq datasets

| **Dataset** | **Sample type** | **Enriched for specific cell types** | **Reference** |
| --- | --- | --- | --- |
| **Human glioma** | | | |
| Mathewson 2021 | IDH mutant and wildtype | Yes - T cells | (Mathewson et al., 2021) |
| Miller 2023 | IDH mutant and wildtype | No - all cells | (Miller et al., 2023) |
| Pombo Antunes 2021 | IDH wildtype | Yes - CD45^POS^ cells | (Pombo Antunes et al., 2021) |
| **Murine glioblastoma** | | | |
| Tomaszewski 2022 | CT-2A | Yes - CD45^POS^ cells | (Tomaszewski et al., 2022) |
| Chen 2023 | 005 | Yes - CD45^POS^ cells | (Chen et al., 2023) |
| Pombo Antunes 2021 | GL261 | Yes - CD45^POS^ cells | (Pombo Antunes et al., 2021) |

Table 2. Murine GB cell line characterization and genetics

| **GB model** | **Established** | **Proliferative** | **Invasive** | **Histology** | **Genetic** | **Reference** |
| --- | --- | --- | --- | --- | --- | --- |
| CT-2A | Chemical induction of methylcholanthrene Subcutaneous | +++ | - | Astrocytoma | P53 WT/Pten deficient | (Haddad et al., 2021; Khalsa et al., 2020) |
| GL261 | Chemical induction of methylcholanthrene Intracranially | ++ | - | Ependymoblastoma | K-Ras mutant/p53 mutant; Pten deficient | (Haddad et al., 2021; Khalsa et al., 2020) |
| 005 | Retroviral transduction | + | +++ | High-grade glioma | H-ras/AKT activation; Pten elevated; p53+/- | (Khalsa et al., 2020; Marumoto et al., 2009) |

Table 3. Re-implanted mice

| **Strain** | **First implant** | **Survival time** | **Second implant** | **Outcome** |
| --- | --- | --- | --- | --- |
| BL6 WT mice | 8/2/23 |  | 4/12/23 |  |
| 1 | CT-2A-FLuc-m4-1BBL | 10 months | CT-2A-FLuc-m4-1BBL | tumor at day 16 |
| 2 | CT-2A-FLuc-m4-1BBL | 10 months | CT-2A-FLuc-m4-1BBL | tumor at day 16 |
| 3 | CT-2A-FLuc-m4-1BBL | 10 months | CT-2A-FLuc-m4-1BBL | No tumor |
| **Strain** | **First implant** | **Survival time** | **Second implant** | **Outcome** |
| BL6 WT mice | 11/2/22 |  | 6/7/22 |  |
| 4 | CT-2A-FLuc-m4-1BBL | 5 months | CT-2A-FLuc | no tumor |
| 5 | CT-2A-FLuc-m4-1BBL | 5 months | CT-2A-FLuc | no tumor |
| 6 | CT-2A-FLuc-m4-1BBL | 5 months | CT-2A-FLuc | no tumor |
| 7 | CT-2A-FLuc-m4-1BBL | 5 months | CT-2A-FLuc | tumor at day 14 |
| **Strain** | **First implant** | **Survival time** | **Second implant** | **Outcome** |
| BL6 WT mice | 8/2/22 |  | 15/01/24 |  |
| 8 | CT-2A-FLuc-m4-1BBL | 23 months | CT-2A-FLuc | tumor at day 10 |
| 9 | CT-2A-FLuc-m4-1BBL | 23 months | CT-2A-FLuc | tumor at day 30 |
| 10 | CT-2A-FLuc-m4-1BBL | 23 months | CT-2A-FLuc | no tumor |
| 11 | CT-2A-FLuc-m4-1BBL | 23 months | CT-2A-FLuc | no tumor |
| 12 | CT-2A-FLuc-m4-1BBL | 23 months | CT-2A-FLuc | no tumor |
| 13 | CT-2A-FLuc-m4-1BBL | 23 months | CT-2A-FLuc | no tumor |
| **Strain** | **First implant** | **Survival time** | **Second implant** | **Outcome** |
| BL6 WT mice | 12/1/24 |  | 14/03/2024 |  |
| 1 | 005-FLuc-m4-1BBL | 3 months | 005-FLuc | No tumor |
| 2 | 005-FLuc-m4-1BBL | 3 months | 005-FLuc | No tumor |
| 3 | 005-FLuc-m4-1BBL | 3 months | 005-FLuc | No tumor |
| 4 | 005-FLuc-m4-1BBL | 3 months | 005-FLuc | No tumor |
